# A spatiotemporal map of the aging mouse brain reveals white matter tracts as vulnerable foci

**DOI:** 10.1101/2022.09.18.508419

**Authors:** Oliver Hahn, Aulden G Foltz, Micaiah Atkins, Blen Kedir, Patricia Moran-Losada, Ian H Guldner, Christy Munson, Fabian Kern, Róbert Pálovics, Nannan Lu, Hui Zhang, Achint Kaur, Jacob Hull, John R Huguenard, Sebastian Grönke, Benoit Lehallier, Linda Partridge, Andreas Keller, Tony Wyss-Coray

## Abstract

Aging is the key risk factor for cognitive decline, yet the molecular changes underlying brain aging remain poorly understood. Here, we conducted spatiotemporal RNA-seq of the mouse brain, profiling 1,076 samples from 15 regions across 7 ages and 2 rejuvenation interventions. Our analysis identified a brain-wide gene signature of aging in glial cells, which exhibited spatially defined changes in magnitude. By integrating spatial and single-nucleus transcriptomics, we found that glia aging was particularly accelerated in white matter compared to cortical regions, while specialized neuronal populations showed region-specific expression changes. Rejuvenation interventions, including young plasma injection and dietary restriction, exhibited distinct effects on gene expression in specific brain regions. Furthermore, we discovered differential gene expression patterns associated with three human neurodegenerative diseases, highlighting the importance of regional aging as a potential modulator of disease. Our findings identify molecular foci of brain aging, providing a foundation to target age-related cognitive decline.

## Introduction

Aging is the predominant risk factor for cognitive dysfunction ^1, 2^ and several neurodegenerative disorders, including Alzheimer’s disease (AD) and Parkinson’s disease (PD) ^3–5^. It remains unclear though, how aging contributes to the development of these distinct diseases of the brain, given their differences in pathological hallmarks, time of onset, and, notably, the regions affected^4^. A quantitative understanding of the dynamics of aging across the brain may provide new insight into the relationship between aging and neurodegeneration. Interestingly, neuroimaging studies using structural and functional magnetic resonance imaging (MRI) data indicate that aging impacts the brain in a region-specific manner ^6, 7^. However, these structural manifestations provide limited insight into the underlying molecular alterations occurring during brain aging. In contrast, changes in gene expression can be a readout of cellular deterioration and molecular processes accompanying aging, permitting quantitative comparisons of aging rates between tissues ^8^ and cell types ^9^. Previous studies have profiled age-related gene expression changes in human brain tissue, yet these microarray-based experiments capture a limited set of transcripts and cover usually one to four regions ^10, 11^ or quantify the transcriptome at low temporal resolution ^12, 13^. Expression profiling during human brain aging is particularly challenging since it can take hours to days before postmortem tissue is stabilized ^13–15^. Alternatively, expression profiling in model organisms like *M. musculus* enables quantitative data with minimal confounding factors, but comprehensive studies covering more than a few regions and at high temporal resolution ^16–19^ do not – to our knowledge – yet exist. In consequence, this limitation also complicates the dissection of molecular mechanisms which mediate the effects of experimental disease models or interventions targeting the aging process, such as dietary restriction or young plasma injection, which delay molecular and cognitive phenotypes of brain aging ^20^.

## Results

### Spatiotemporal quantification of age-related gene expression across the mouse brain

To obtain a deep molecular understanding of the spatiotemporal changes of the aging mammalian brain, we selected 15 brain regions in the mouse with critical functions in cognition, whole-body metabolic control, or human disease. We punched out brain regions from coronal brain sections from the left and right hemisphere, including three cortical regions (motor area, visual area and entorhinal cortex; Mot.cor., Vis.cor and Ent.cor, respectively), anterior (dorsal) and posterior (ventral) hippocampus (Hipp.ant and Hipp.post., respectively), hypothalamus (Hypoth.), thalamus, caudate putamen (part of the striatum; Caud.put.), pons, medulla, cerebellum (Cereb.) and the olfactory bulb (Olf.bulb). We further isolated three regions that were enriched with the corpus callosum (Corp.cal.), choroid plexus (Chor.plx.) and the neurogenic subventricular zone (SVZ), (Figure S1A). We then applied our method to 59 mice (Figure 1A; n = 3-6 males per age; aged 3, 12, 15, 18, 21, 26 and 28 months; n = 5 females per age; aged 3, 12, 15, 18 and 21 months; all C57BL/6JN strain), resulting in a total of 1,770 samples (885 samples from each hemisphere). Isolated regions from the left hemisphere were stored, while all right hemisphere regions were processed through a custom-built bulk RNA-seq (bulk-seq) pipeline (Figure 1B, STAR Methods). We achieved robust tissue sampling with high RNA quality while minimizing perfusion artifacts, as indicated by consistent RNA yields across samples from the same region (Figure S1B), median RNA integrity numbers of 9.45 out of 10 (Figure S1C) ^21^, and a neglectable fraction of reads mapping to hemoglobin genes (Figure S1D).

**Figure 1.**
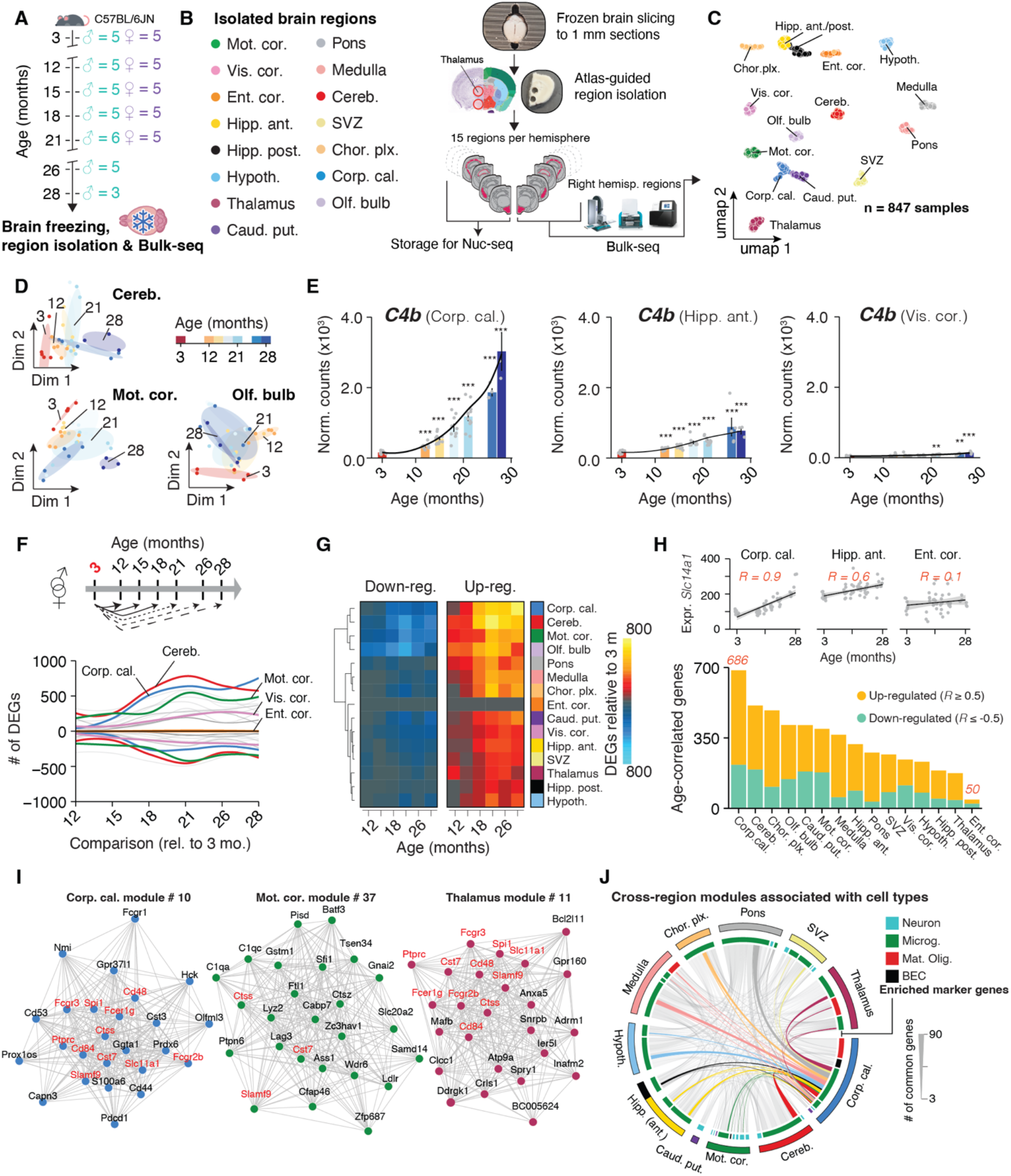
Brain regions exhibit distinct transcriptional patterns of aging independent of anatomical proximity (A) Cohort overview. Whole brains were collected from male (n = 3-5, 3–28 months) and female (n = 5, 3–21 months) C57BL/6JN mice. (B) Study outline. 15 brain regions were isolated from each hemisphere of the brains collected in (A). Regions from the right hemisphere were analyzed using Bulk-seq. (C) UMAP representation of brain region transcriptomes (n = 847 total samples), based on the first 40 principal components. (D) Diffusion maps of region transcriptomes from male cerebellum, motor cortex, and olfactory bulb. Dim., dimension. (E) *C4b* expression in corpus callosum, anterior hippocampus and visual cortex. Black lines indicate averaged-smoothed gene expression. Differential expression relative to the 3 months group is indicated. Data are mean ± s.e.m. Two-sided Wald test, adjusted for multiple testing. *** p < 0.001, ** p < 0.01, * p < 0.05.

After quality control, we obtained 847 single-region transcriptomes. Visualization in uniform manifold approximation and projection (UMAP) space separated samples by region (Figure 1C), but not sex or age, which concurred with deterministic shared-nearest-neighbors graph clustering and hierarchical clustering (Figure S2A-E). However, within individual regions, samples segregated transcriptionally by age. The comparatively subtle effect of aging on gene expression highlights the necessity for precise isolation of brain tissue to avoid confounding cross-region contamination (Figure 1D and S2A).

To assess if the isolated regions accurately captured a given brain structure’s transcriptome, we analyzed the expression of region-enriched genes (‘marker genes’; STAR Methods and Table S1) in a publicly-available spatial transcriptomics dataset of an adult male mouse brain ^22–24^. To this end, we combined marker genes of a given region into ‘signatures’ ^25^ that represent its transcriptional fingerprint. For each signature, we calculated a score per spatial transcriptome spot, summarizing the expression of marker genes into a single value. Signature scores were distinctly elevated in areas corresponding to the anatomical structures annotated in the Allen brain reference atlas (Figure S3A) ^26^. Notably, the corpus callosum-derived signature demarcated fiber tracts throughout the brain, indicating that the sampled transcriptome of this region could be a proxy for white matter tracts in general (Figure S3A). To assess the isolation of the SVZ, we built a signature of activated neural stem cells (aNSC) based on marker genes from single-cell data ^27^ and calculated the score per region. Not only was the score elevated in the aNSC-rich olfactory bulb and SVZ region but we also found a significant decline with age, indicating a loss of aNSCs with age, that is in agreement with diminished neurogenic capacity in aged mice. (Figure S3B) ^27^. In summary, our tissue isolation and bulk-seq workflow yielded high-quality transcriptomes that robustly captured a region’s gene expression profile across a cohort of mice. The data can be interactively explored at https://twc-stanford.shinyapps.io/spatiotemporal_brain_map/.

### Region identity is linked to expression dynamics during aging

RNA-seq permits quantitative comparisons of aging rates between organs and cell types ^8, 19^ based on timing and effect size of gene expression shifts. For instance, we found substantial region-dependence in the magnitude and timing of *C4b* expression (Figure 1E), a complement component and major schizophrenia risk factor ^28^ that is robustly up-regulated in aged mice ^29^ and models of neurodegeneration ^30^. Notably, recent single-cell sequencing and spatial imaging studies revealed that the composition of major cell types remains almost constant throughout the aging mouse brain between 3 and 21 months ^31, 32^, thus the expression dynamics observed in bulk are unlikely to be driven primarily by shifts in cell type abundance. In cases of stable cell populations and substantial numbers of biological replicates, bulk RNA-seq is particularly suitable to investigate subtle, yet robust expression changes as they occur during aging. This is in part due to well-established, replicate-sensitive statistical approaches ^33^ that currently do not exist for single-cell data ^34^. Thus, we were able to use our temporally resolved data to probe the per-region impact of aging on gene expression over time, as this could help to identify structures with selective vulnerability to old age.

We performed pairwise differential expression between 3 months and every following age group to determine when differentially expressed genes arise (DEGs; used from hereon to refer to genes that change with age). Given that very few genes exhibit a significant interaction between sex and age (Figure S4A-C) – thus indicating a general collinearity of aging effects in both sexes – we treated sex as a covariate to maximize statistical power for identifying DEGs and mapping out transcriptional shifts that occur in both males and females. To focus on high-confidence expression changes that persist with advancing age, a gene had to pass the statistical cutoff in at least two comparisons to be classified as a DEG (Figure 1E-G). The general trend across regions indicated an increase of DEGs over time that plateaued around 21 months (Figure 1F,G), yet individual regions varied profoundly with respect to their total number of DEGs and the trajectory of DEG accumulation (Figure 1F and Table S2). For instance, the visual cortex showed a steady increase of DEGs until late age, while the transcriptome of the motor cortex already exhibited significant perturbation at 12 months, but there was little increase until a sudden jump at 21 months (Figure 1F,G). In contrast, the transcriptome of the entorhinal cortex appeared largely refractory to the effects of age altogether, with only 13 detectable DEGs in total (Figure 1F,G). This is in line with human MRI ^7^ and microarray ^35^ studies demonstrating that the entorhinal cortex displays only mild alterations during cognitively normal human aging, whereas it frequently exhibits the first amyloid deposition in AD patients ^36^. Together, these results reveal that the effect size of expression shifts during brain aging are strikingly region- and time-dependent, highlighting the necessity for region-resolved quantification and analysis. Notably, the regions with the most profound and earliest shifts in gene expression were the white matter-rich caudate putamen, cerebellum, and corpus callosum, the latter showing a tenfold increase in the number of DEGs between 12 and 18 months. Since pairwise comparisons treat every gene and age group independently, we next validated these results with two independent analyses. First, we probed all genes in the genome for positive or negative correlation with age (Spearman’s rho ≥ 0.5 or ≤ -0.5, respectively; padj ≤ 0.05; Table S3), thus taking all age groups into account (Figure 1H). Not only did the regions differ in the number of age-correlated genes, confirming that the effect size of age depends on the region, but also the corpus callosum and cerebellum were the most impacted, while the entorhinal cortex remained largely unaffected (Figure 1H). As a second validation, we performed weighted gene co-expression network analysis (WGCNA) ^37^ for each region (STAR Methods; Table S4), clustering genes into modules which might be driven by similar regulation during aging. We filtered for modules exhibiting significant association with age and found the number of modules differed between regions. In line with the above results, we found seven or more modules in the corpus callosum, cerebellum and motor cortex, whereas we detected no age-related modules in the entorhinal cortex (Table S4). To gain biological insight, we performed cell type- and pathway-enrichment for each age-related module and compiled summarized reports for each region as a quick reference resource for the scientific community (https://twc-stanford.shinyapps.io/spatiotemporal_brain_map/). Interestingly, we discovered in 10 regions at least one module with increased expression over time that was enriched for microglia- and inflammation-related genes (Fig 1I,J). Consistent with these findings, we found a small, common set of differentially regulated genes, including neuroinflammatory markers *Fcgr2b*, *Ctss*, *Cst7* ^38^ in modules across regions, suggesting the presence of a minimal group of co-regulated genes changing throughout the brain. In summary, we found the results of three independent analyses (pairwise tests, age-correlation and WGCNA) congruent, demonstrating that the observed effects of aging on the transcriptome are region specific.

### A minimal gene set forms a common fingerprint of brain aging

WGCNA analysis results indicated the possibility of a shared gene set that changes during aging throughout the brain. Such a minimal age-related gene signature would permit quantitative comparisons of the rates of change in a region’s transcriptional age. While the vast majority of DEGs appeared to change only in three or less regions, indicative of region-selective expression patterns, we found 82 genes that were differentially regulated in 10 or more regions (Figure 2A,B; Table S2). These were strongly enriched for up-regulated genes with immune-modulatory functions (Table S5), including MHC I-mediated antigen presentation, interferon-response, cell adhesion and complement cascade, as well as regulators of mouse microglia activity (Figure 2C) including *Cd22* ^39^, *Trem2* and *Tyrobp* ^40^. Interestingly, of the only 7 down-regulated genes in this set, we found protein homeostasis genes *Dnajb1*, *Hsph1* and *Ahsa1*, as well as collagen synthesis gene *P4ha1*, which is in line with perturbed protein quality control mechanisms as a hallmark of aging ^3^ (Figure 2B). We combined these 82 genes into a common RNA aging signature to calculate their expression as a single ‘common aging score’ (CAS; STAR Methods) for each mouse and region. While the CAS expectedly showed significant increases in every region (Figure 2D, Figure S5A), the shape and amplitude of the trajectories varied profoundly. The pace and direction with which the CAS changed with age is defined from here on as a region’s ‘CAS velocity’. We employed linear models to approximate these trajectories, using the slope of the linear fit as a metric to comparatively assess the CAS velocity across regions (Figure 2D,E). Of note, the CAS at baseline (i.e. the offset of the linear fit) did not predict a region’s CAS velocity (Figure 2F). Our analysis revealed a gradient of velocities across regions, with the three cortical areas and the olfactory bulb ranking last, at approximately one-third of the velocity of the corpus callosum, the ‘fastest’ region (Figure 2G,H,I). Other white matter-rich areas such as the caudate putamen also exhibited high velocities, while the hippocampus, thalamus and hypothalamus – some of the most investigated regions in mouse and human brain aging research ^18^ – ranked slightly below average. The median CAS across all regions associated with the animals’ chronological age (Figure S5B).

**Figure 2.**
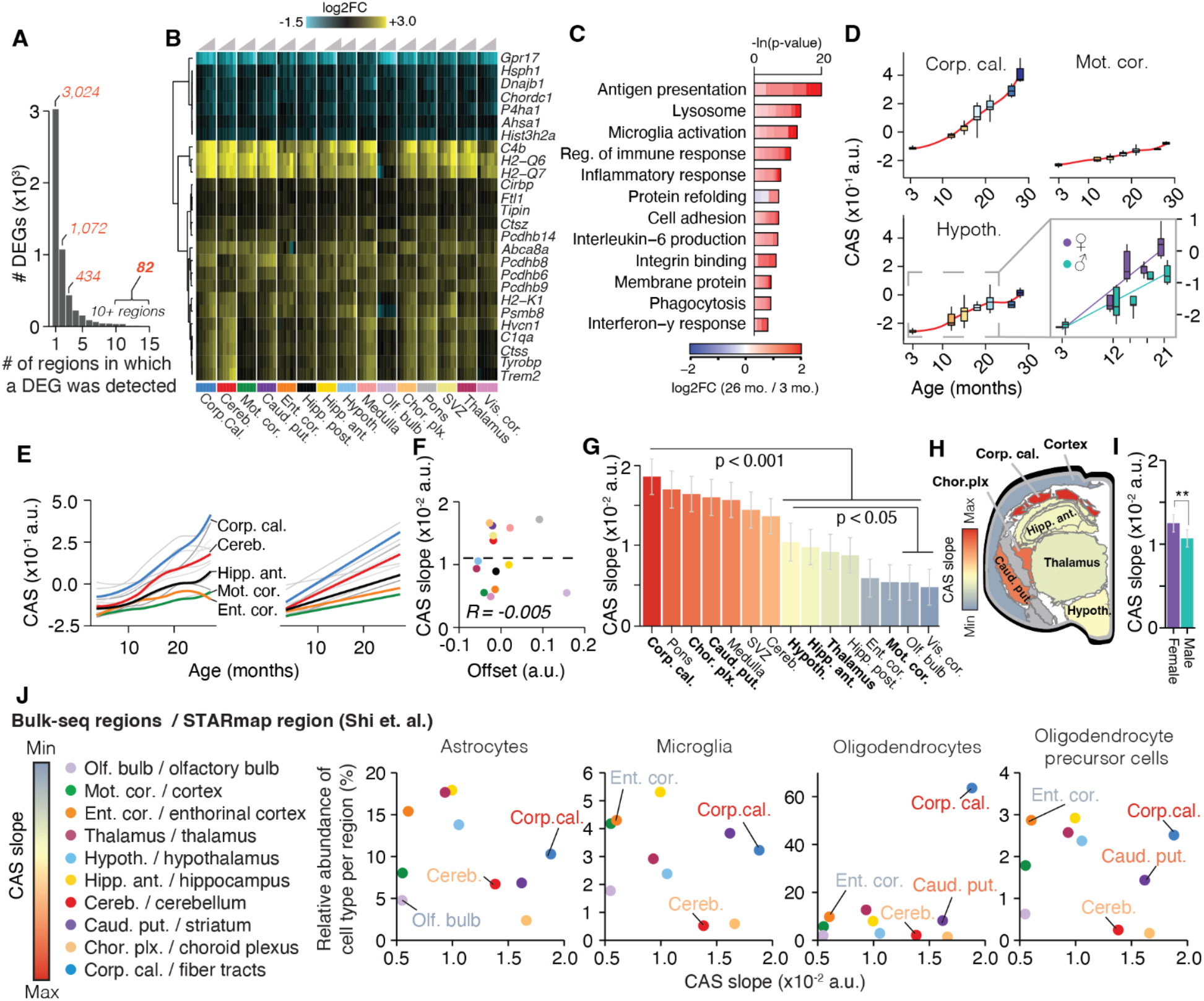
A common gene signature identifies regions with accelerated aging (A) Bar graph indicating the number of regions in which a given DEG was detected (Table S2). (B) Region-wise expression changes with age (column-wise from left to right) for genes with shifts in at least 10 of the 15 collected regions. (C) Representative GO analysis of 82 genes with shifts in at least 10 of the 15 collected regions that make up the CAS. Lengths of bars represent negative ln-transformed Padj using two-sided Fisher’s exact test. Colors indicate gene-wise log2 fold-changes (log2(FC)) between 26 and 3 months old mice as measured in the corpus callosum. The complete list of enriched GO terms can be found in Table S5. (D) CAS trajectories in the corpus callosum, motor cortex, and hypothalamus. Insert indicates trajectories for male and females in the hypothalamus from 3 to 21 months. (E) CAS trajectories of all regions approximated via local estimate (LOESS) and linear regression, colored by region; gray lines represent non-labelled regions. (F) Offset and slope comparison for linear models in (E), colored by region. Linear regression (dashed line) and Spearman correlation coefficient are indicated. (G) Slope of linear regressions in (D), colored by slope. Data are mean ± 95% confidence intervals. Bolded regions are highlighted in the following panel. (H) Coronal cross-section sketch of the mouse brain, with regions colored according to CAS linear slopes. Corpus callosum was chosen to represent white matter tracts. (I) Slope of linear regression across all brain regions from 3 to 21 months, colored by sex. Data are mean ± 95% confidence intervals. Two-sided Tukey’s HSD test, adjusted for multiple testing, *** p < 0.001, ** p < 0.01, * p < 0.05. The highest (least significant) Pval is indicated. (J) Correlation of the relative abundance of major glia cell types (astrocytes, microglia, oligodendrocytes and OPCs) with the regions’ respective CAS slopes. Each plot shows the relative abundance of a different cell type in each of the regions, (determined from the equivalent region in ^41^, with CA1, CA2, CA3, dentate gyrus etc. were grouped together into a ‘hippocampus’ region) plotted against the CAS slope (determined from our Bulk dataset). Each point represents a different region. Significance of this relationship tested through spearman correlation and linear regression, with no significant trends found.

Yet, the regions’ differing velocities resulted in increased per-animal variance, indicating that the transcriptional state of this gene set becomes profoundly desynchronized across the brain. This appeared to be independent of the regions’ anatomical location, as the fast-aging corpus callosum stretches between the slow-aging cortical areas and hippocampus (Figure 2H). Importantly, we found no association between the CAS velocity and each region’s relative cell composition at young age as quantified in a recent brain-wide in-situ single-cell sequencing dataset ^41^ (Figures 2J, S6A STAR Methods). This suggests that the heterogeneous CAS velocities are unlikely to result from differences in cell proportions across regions.

When we examined the CAS trajectories for the interval between 3 and 21 months, we observed moderate but significant acceleration of the CAS in females compared to males (Figure 2I, S7A,B). In particular, the hypothalamus exhibited the most pronounced acceleration in females (Figure S7B). Aiming to identify which exact CAS genes would contribute to this sexual dimorphism, we analyzed age-correlated genes in the male and female hypothalamus separately (Figure S7C, STAR Methods). While overall age-related expression changes were well-correlated between both sexes (*P* value for Fisher’s Exact test < 2.2x10^-16^), genes related to lipid metabolism, stress response and unfolded protein response, including CAS genes *Rbm3* and *Cirbp* ^42^, tended to exhibit stronger regulation in males (Figure S7D,E). In contrast, females exhibited a more accentuated regulation of markers of neuroinflammation (such as *Gfap*) and antigen-presentation genes (Figure S7F), as well as several chemokines and CAS genes related to immune response. In line with this, we found *Cish*, a known regulator of T cell immune response increasing in females ^43, 44^, being the only gene exhibiting significant, opposite regulation in both sexes. Critically, the female-specific regulation of pro-inflammatory genes was not observable in other regions with similar CAS slopes (Figure S7F). These findings are in line with human studies reporting more pronounced expression of immune-related genes in the hippocampus and cortex of aged women ^35, 45^. Of note, adenomas in the pituitary gland of female C57BL/6J mice, which is directly connected to the hypothalamus, can develop at high frequency in female C57BL/6J mice older than 20 months of age^46^. Since we did not record adenomas in our study, we cannot exclude the possibility that this phenomenon could contribute to the accelerated aging patterns observed in the hypothalamus. Our data could advance the understanding of several sexual-dimorphisms observed in the brain, including the higher age-specific risk of dementia among women ^47^ in general, and the dynamics of reproductive aging in particular, given the hypothalamus’ critical role in regulating reproduction, development and metabolism ^48^.

### Fiber tracts are foci of accelerated brain aging

Bulk-seq data – even with the regional dissection conducted here – could mask transcriptional changes with age that occur in sub-structures of regions, such as specific layers of the cortex. We thus aimed to validate our CAS analysis with a fine-resolution method that would still capture multiple regions in the same assay. To this end, we performed spatial transcriptomics (10X Visium) of the brain across aging, isolating coronal sections from an independent cohort of male mice aged 6, 18, and 21 months (Figure 3A). Using a clustering-based approach to annotate the regional identity of Visium spots (Figure S8A,B; STAR Methods; Table S6) we identified them as belonging to the hippocampus, cortex (motor and somatosensory area), thalamus, hypothalamus, striatum (including the caudate putamen), choroid plexus and white matter fiber tracts (including the corpus callosum) (Figure 3B and S8C-F). Our data demonstrated robust capture of the same regions across age groups and individuals (Figure S8G-L), thus enabling the comparison of DEGs found in bulk-seq with Visium data (Table S7). We confirmed a more pronounced regulation of DEGs in the white matter cluster (equivalent to the dissected corpus callosum region) compared to the cortex cluster (equivalent to the motor cortex region), including several of the 82 CAS genes (Figure 3C, Table S7) such as *Trem2* (Figure 3D). Calculating CAS for each Visium spot identified a distinct, spatially-confined increase of the score along the white matter tracts, including not only the corpus callosum but also other fiber tract sub-structures such as the fimbria and internal capsule (Figure 3E,F). In the cortex, however, we observed only a mild increase of CAS with age. In general, CAS velocities calculated via bulk-seq and those calculated via spatial transcriptomics were well-correlated (Figure 3G), confirming vastly differing aging velocities between proximal regions *in-situ*.

**Figure 3.**
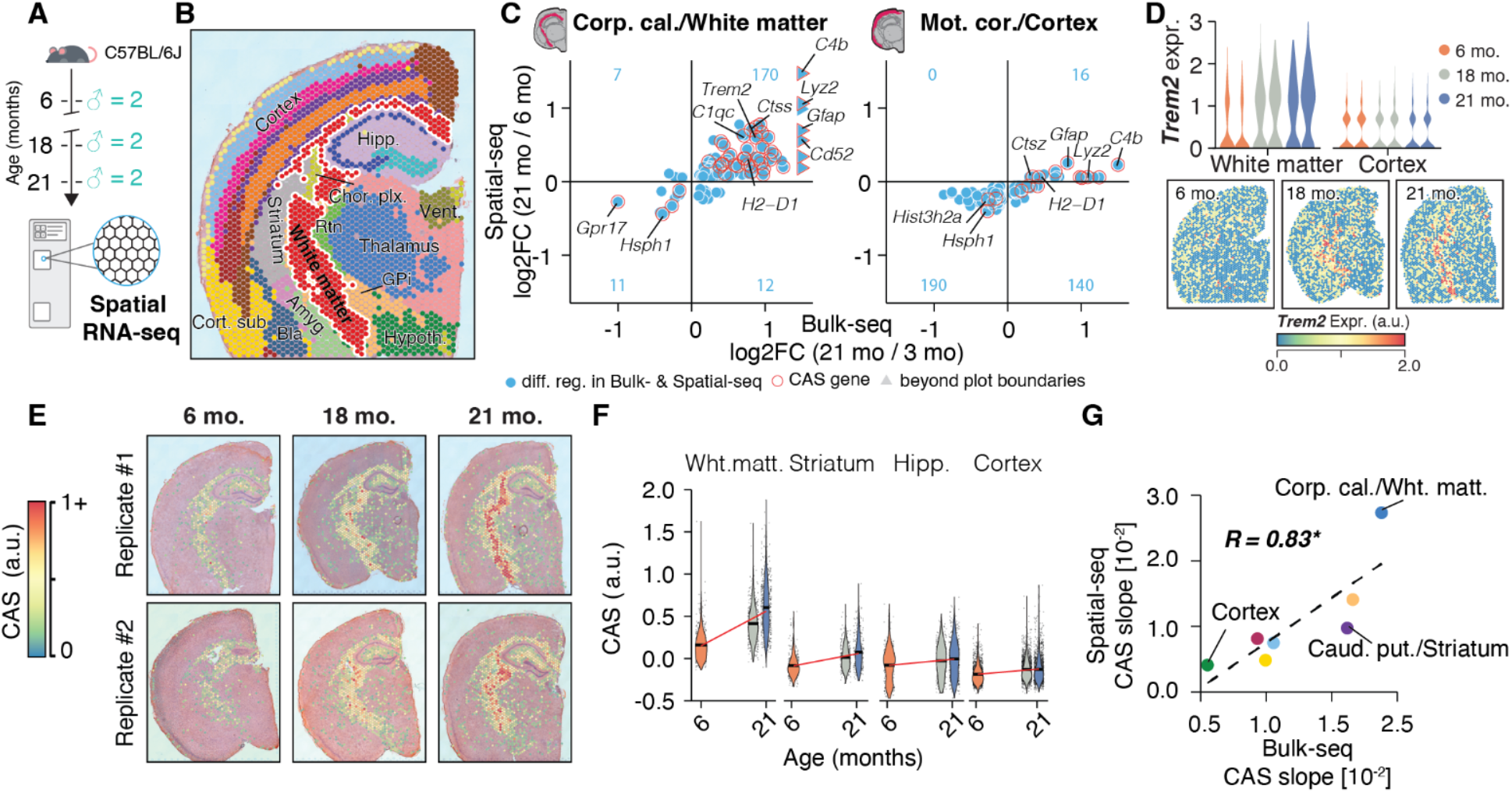
Spatially-resolved CAS detects accelerated aging in white matter tracts (A) 10X Visium experiment overview. Brain tissue was collected from an independent male C57BL/6J mouse cohort (n = 2 mice; 6, 18 and 21 months). (B) Representative spatial transcriptome data (6 months replicate #2), colored by cluster-based annotation, according to Fig S8. Labels represent region-level annotation according to Fig S8. Complete data description and abbreviations are in Fig S8. (C) Comparison of Bulk-seq and Visium differential expression results in white matter cluster/corpus callosum punch; cortex cluster/motor cortex punch. DEGs (Padj < 0.05) found in both datasets are shown, with their log2-transformed expression ratios (21 rel. to 3 months) in Bulk-seq and Visium data. CAS genes are highlighted. The number of overlapping DEGs in each quadrant is indicated in blue. (D) Spatially-resolved expression of *Trem2* across age. Violin plots represent expression in white matter- and cortex-associated spots, split by replicates. (E) Spatial representation of CAS. Spots with values ≥ 0 are shown. (F) Violin plot representing CAS across spatial clusters of white matter, striatum, hippocampus and cortex. Red line indicates linear regression fit. (G) Comparison of CAS slopes for linear models in Bulk-seq and Visium data, colored by region. Linear regression (dashed line) and Spearman correlation coefficient are indicated. Corpus callosum, caudate putamen and motor cortex regions were chosen to represent white matter, striatum and cortex, respectively.

### Heterogeneous velocity of CAS is encoded by glial transcripts

We next sought to quantify the activity of the CAS genes at the single-cell level to identify the cell type(s) that shape the heterogeneous expression dynamics across brain regions. We chose the anterior hippocampus as a representative region given its intermediate CAS velocity (Figure 2G), utilizing frozen punches from the left hemispheres of the bulk-seq cohort (Figure 4A). Single-nuclei sequencing (nuc-seq) yielded all major cell types, with no evidence for a shift in cell type composition with age or sex. We observed the highest baseline CAS in microglia, in line with several of the CAS genes being known immune-response genes (Figure 2B,C). While the CAS showed a statistically significant increase in all cell types (Figure 4B), including a mild elevation in several neuronal populations, the most accentuated increase was observed in microglia (Figure 4C), followed by mature oligodendrocytes, brain endothelial cells (BECs), astrocytes and oligodendrocyte progenitor cells (OPCs).

**Figure 4.**
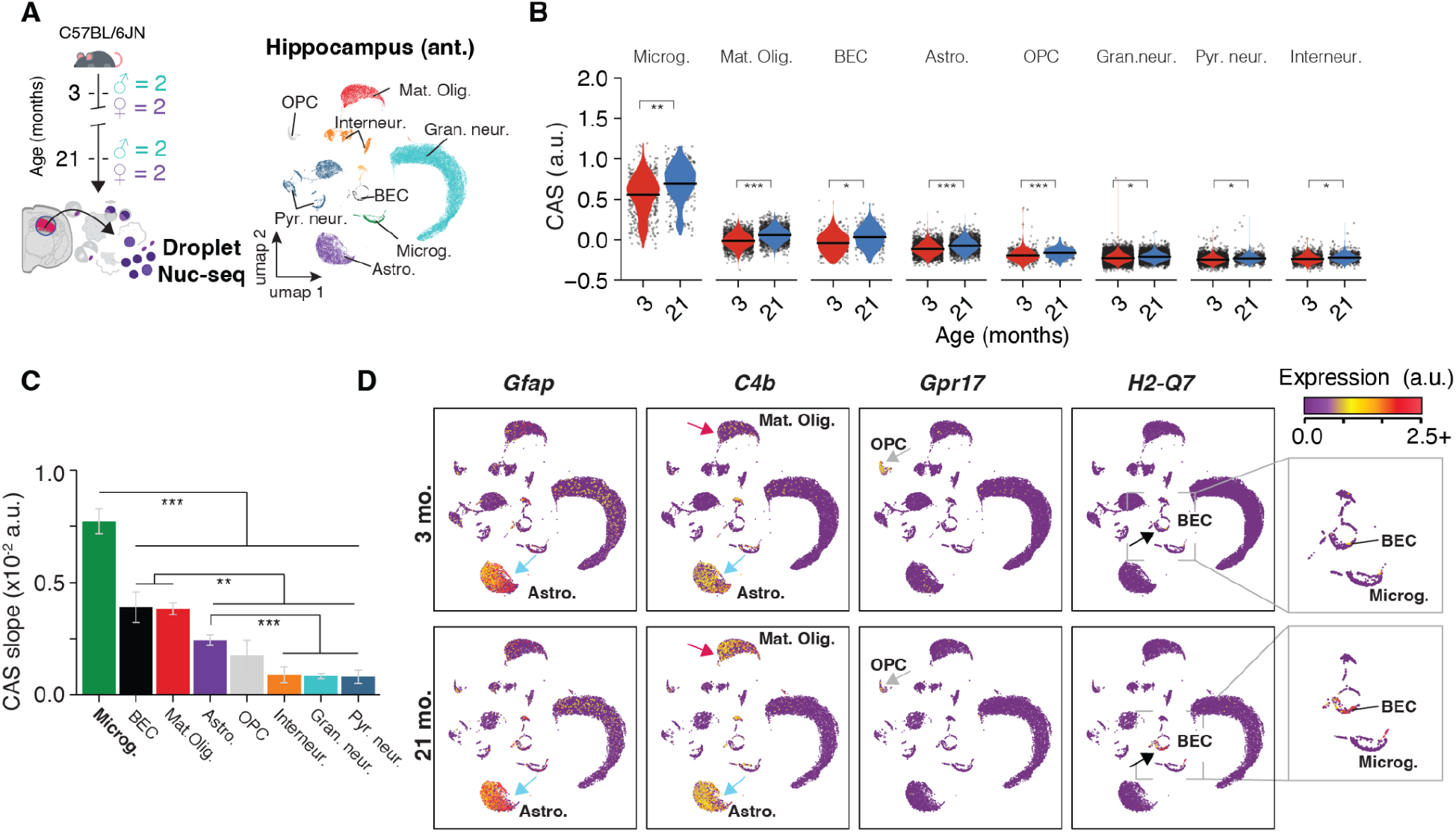
Aging in glia and endothelial cells is the major contributor to CAS increase (A) Nuc-seq experiment overview. Nuc-seq of left-hemisphere regions of the anterior hippocampus from the same mice used for bulk RNA-seq (n = 2 males, n = 2 females; 3, and 21 months). UMAP representation of all nuclei populations (n = 36,339 cells). (B) Violin plot representing CAS across hippocampal cell types. Points indicate nuclei-wise expression levels, and the violin indicates average distribution of expression split by age. *P* values calculated with two-tailed t-test on per-replicate median of CAS, adjusted for multiple testing. *** p < 0.001, ** p < 0.01, * p < 0.05 (C) CAS slope of linear regressions in (B), colored by cell type. Data are mean ± 95% confidence intervals. Two-sided Tukey’s HSD test, adjusted for multiple testing, *** p < 0.001, ** p < 0.01, * p < 0.05. The highest (least significant) Pval is indicated. (D) Expression of CAS genes *Gfap*, *C4b*, *Gpr17* and *H2-Q7*. Quantification and statistical analysis can be found in Figures S9 and S10.

Upon closer examination of the 82 genes, it became clear how the CAS could reflect aging dynamics for several cell types beyond microglia by cell type-specific or cell type-selective gene expression shifts (Figure 4D), including *Gfap* (Astrocytes; Figure S9A), *C4b* (Astrocytes and mature oligodendrocytes; Figure S9B-E), *Gpr17* (OPCs; Figure S10A-E) and *H2-Q7* (BECs; Figure S10F). Of note, this analysis also demonstrated that aging can induce expression of genes that are not detected at young age. For instance, *C4b* was mostly expressed in astrocytes at young age, however its expression became detectable and increased foremost with age in mature oligodendrocytes (Figure S9E). Similarly, expression of *H2-Q7* only became detectable in BECs with old age (Figure S10F). We validated our findings in an independent dataset, using publicly available scRNA-seq data from dissected SVZ of young and old male mice ^27^ (Figure S11A). Though generated using a different cohort, region and method, the CAS increase was most pronounced in microglia which is consistent with our data (Figure S11B-G). There was also a profound increase of CAS in aNSCs, though the very low number of cells at 28 months (less than 50 per animal) complicates robust calculations of CAS at this age. Thus, the region-to-region differences in CAS velocity are predominantly reflecting age effects in non-neuronal cell types, with microglia having the strongest contribution.

### Transcriptional aging of microglia is region-dependent

We finally examined if there were varying CAS dynamics between microglia from regions with fast or slow CAS velocity. To this end, we analyzed Smartseq2 scRNA-seq data from the *Tabula Muris* consortium (Figure 5A), where comparable numbers of microglia were collected from the freshly-isolated cerebellum, striatum, hippocampus and cortex, which we considered sufficient equivalents to the cerebellum, caudate putamen (both areas with high velocity), anterior hippocampus (medium velocity) and motor/visual cortex (low velocity) regions (compare Figure 2G). The Smartseq2 protocol is particularly suitable to examine subtle transcriptional effects occurring during aging – especially those observed at the bulk level – due to its efficient per-cell transcript capture rate ^49–51^. Indeed, as predicted by our bulk-seq results, the CAS in aged microglia increased in all four regions significantly, though with greater magnitude in the cerebellum and striatum, followed by the hippocampus and cortex, respectively (Figure 5B). These shifts were well-represented by all biological replicates (Figure S12A). The same trend was detectable on the level of individual CAS genes, like *Trem2* (Figure 5C). Notably, there was no detectable CAS difference among microglia at young age across the striatum, hippocampus and cortex, while the cerebellum-derived microglia exhibited a slightly higher CAS at baseline. In agreement with our data, the CAS of microglia isolated from white matter of aged mice ^51^ were significantly elevated compared to microglia derived from gray matter (Figure S12B,C). Further, we meta-analyzed a well-powered bulk microarray dataset of microglia isolated from cerebellum, striatum, hippocampus and cortex ^52^, derived from mice at 4, 12 and 22 months of age (Figure S12D). Here, we found more differentially expressed genes with age in the cerebellum and striatum (Figure S12E), together with a more pronounced up-regulation of CAS genes (Figure S12F-H), particularly in the period of 12 to 22 months.

**Figure 5.**
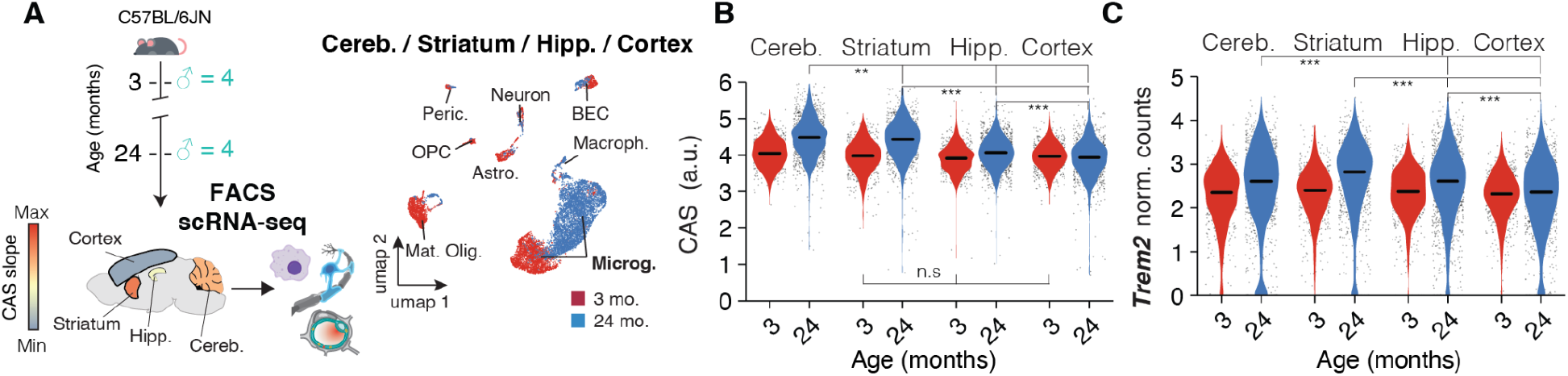
CAS analysis reveals that transcriptional aging of microglia depends on their region of origin. (A) Meta-analysis of scRNA-seq data from ^19^ of microglia from cerebellum, striatum, hippocampus and cortex. UMAP representation of all cell populations (n = 6,373 cells), colored by age. Regions are colored according to CAS slopes in Figure 2G. (B,C) Violin plot representing (B) CAS and (C) *Trem2* expression across microglia from four different brain regions. Points indicate nuclei-wise expression levels, and the violin indicates average distribution of expression split by age. (MAST, Benjamini–Hochberg correction; false discovery rate (FDR) < 0.05 and logFC > 0.2 to be significant). *** p < 0.001, ** p < 0.01, * p < 0.05.

We conclude that the CAS velocities observed in bulk-seq and Visium data are, in part, representing microglia that exhibit region-specific magnitudes of aging. Future studies should examine if other non-neuronal cells, like mature oligodendrocytes or BECs, would exhibit a similar CAS heterogeneity.

### Neuronal transcripts encode region-specific expression patterns

Given that the CAS genes represent only 1.5% of all DEGs (Figure 2A; Table S2), we hypothesized that the remainder could represent region-specific expression shifts. We first compared age-related DEGs across mouse organs to construct organ-specific signatures of aging (Figure S13, STAR Methods). The presence of gene sets with specific regulation found in functionally distinct organs led us to investigate whether individual brain regions exhibit a similar degree of specificity during aging. We found that the number of region-specific DEGs varies greatly (Figure 6A), which we utilized to build aging signatures for each region before calculating the respective score across all other regions (Figure 6B,C, Figure S14A,B). As exemplified by the specific signature of the caudate putamen – a region marked by intertwined gray and white matter structures – we found that most region-specific scores increased with age predominantly in the region on which they were based (Figure 6B-D and Figure S14A,B). Except for the thalamus, pons and SVZ, a given region’s signature velocity outperformed those of all other regions. Thus, dozens to hundreds of genes in the brain are regulated in a region-specific or -selective manner, revealing highly compartmentalized effects of aging within a single organ.

**Figure 6.**
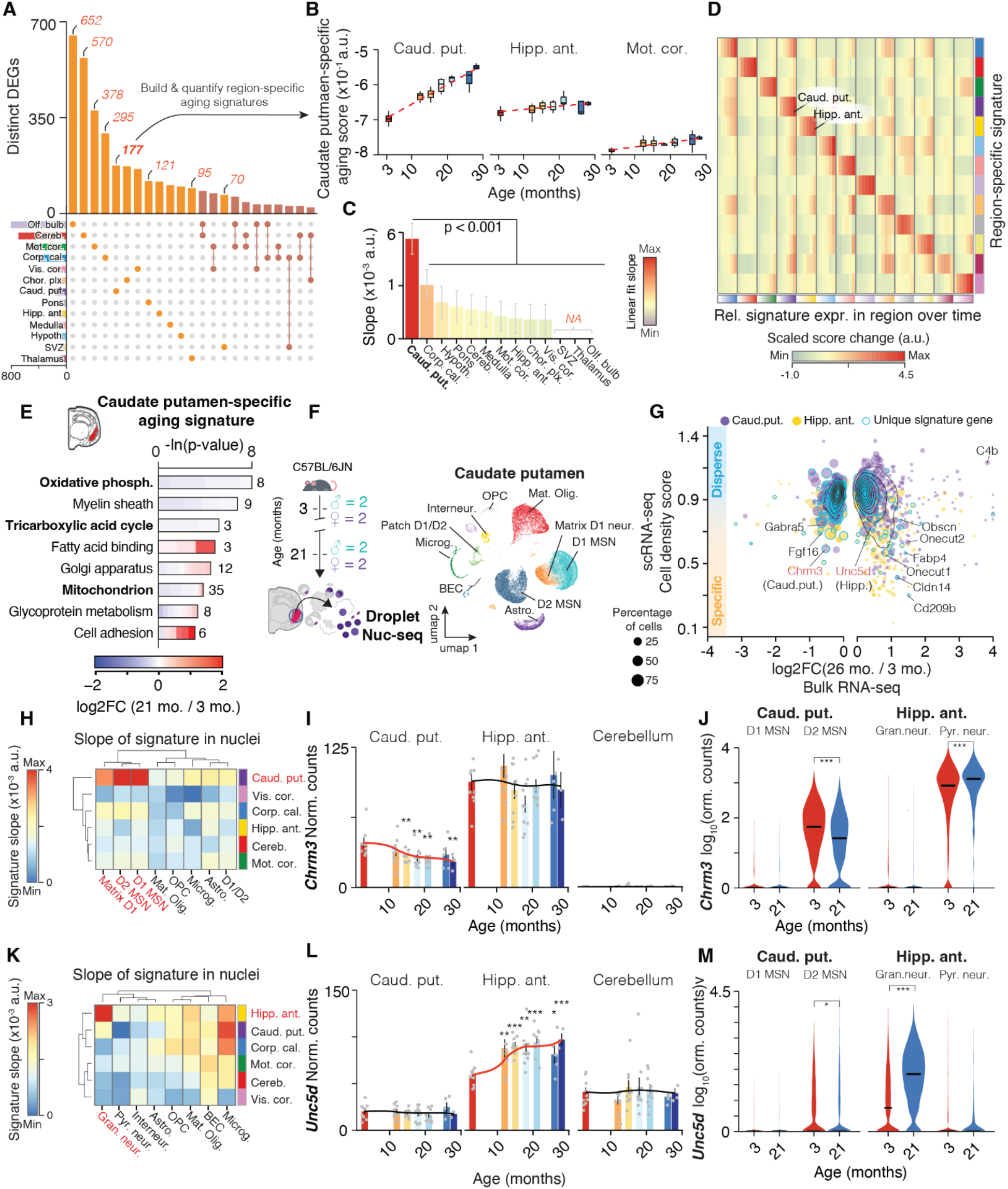
Region-specific expression shifts are encoded by neuronal transcripts (A) Regional specificity of DEGs. UpSet plot showing a matrix layout of DEGs shared across and specific to each region. Each matrix column represents either DEGs specific to a region (single circle with no vertical lines) or DEGs shared between regions, with the vertical line indicating the regions that share that given DEG. Top, bar graph displays the number of DEGs in each combination of regions. Left bottom, bar graph displays the total number of DEGs for a given region. Gene sets with ≥ 25 genes are shown. Unique gene sets were used to construct region-specific aging signatures. (B) Trajectories of caudate putamen-specific aging score in the caudate putamen, anterior hippocampus and motor cortex. Linear fit is indicated as dashed lines. (C) Slope of linear regressions in (B), colored by slope. Data are mean ± 95% confidence intervals. Two-sided Tukey’s HSD test, adjusted for multiple testing, *** p < 0.001, ** p < 0.01, * p < 0.05. The highest (least significant) Pval is indicated. (D) Region-wise score changes with age relative to 3 months (column-wise from left to right) for region-specific signatures. Score changes are z-scaled within a row. Quantification and statistical analysis can be found in Figure S14. (E) Representative GO enrichment as in (Figure 2C) for 177 DEGs unique to the caudate putamen that make up its specific signature. The complete list of enriched GO terms can be found in Table S8. (F) Nuc-seq experiment overview. Nuc-seq of left-hemisphere regions of the caudate putamen from the same mice used for bulk RNA-seq (n = 2 males, n = 2 females; 3, and 21 months). UMAP representation of all nuclei populations (n = 45,277 cells). (G) Single-nuclei dispersion scores plotted against log2-transformed expression ratios between 21 and 3 months (bulk RNA-seq) for the caudate putamen and anterior hippocampus. The colors represent different organ types and size of the dots corresponds to the percentage of cells that express a given gene. Genes that make up the region-specific score are highlighted. (H) Slope of cell type-wise changes with age for representative region-specific signatures from (D). D1 and D2 Medium spiny neuron populations (MSN) are highlighted, as they exhibit a distinct increase with age exclusively for the caudate putamen-specific signature. (I) Bulk expression across caudate putamen, anterior hippocampus and cerebellum for *Chrm3*. Black lines indicate averaged-smoothed gene expression. The trajectory with significant age effect is highlighted. Data are mean ± s.e.m. (J) Violin plot representing *Chrm3* expression across neuronal cell types in caudate putamen and anterior hippocampus. Points indicate nuclei-wise expression levels, and the violin indicates average distribution of expression split by age. (MAST, Benjamini–Hochberg correction; false discovery rate (FDR) < 0.05 and logFC > 0.2 to be significant). *** p < 0.001, ** p < 0.01, * p < 0.05. (K) Slope of cell type-wise changes with age for representative region-specific signatures from (D). Granule cells are highlighted, as they exhibit a distinct increase with age exclusively for the anterior hippocampus-specific signature. (L) Same as (I) for *Unc5d*. (M) Same as (J) for *Unc5d*.

Notably, signature genes appeared to be functionally connected, as exemplified by the caudate putamen-specific signature which was enriched for down-regulated mitochondrial processes and up-regulated cell adhesion and lipid binding functions (Figure 6E and Table S8). To map out the cell types driving this region-specific signature, we analyzed nuc-seq data from the left hemisphere punches of the anterior hippocampus (Figure 4A) and caudate putamen (Figure 6F), where we captured non-neuronal cell types as well as striatum-specific D1- and D2-type medium spiny neurons (D1 and D2 MSNs, respectively). We were able to map several signature genes like *Fgf16*, *S100a10* and *Fabp4* (Figure S14C-E) to distinct cell populations (Figure 6G, STAR Methods and Table S9) suggesting that bulk tissue can indeed capture the expression dynamics of specific cell subsets. Similar to the cell-resolved analysis of the CAS (Figure 4B), we calculated several region-specific signature scores for each cell type in young and old individuals. We found a distinct increase of the caudate putamen-specific signature in D1 and D2 MSNs which was not seen with signatures from other regions (Figure 6H). For instance, expression of muscarinic acetylcholine receptor gene *Chrm3* dropped significantly in the caudate putamen, reflecting down-regulation of this gene specifically in D2 MSNs (Figure 6I,J). In comparison, dentate gyrus granule cells of the hippocampus exhibited a distinct increase of the hippocampus-specific signature (Figure 6K), and we found granule cell-specific regulation of several signature genes such as axon-guidance receptor *Unc5d* ^53^ as well as transcription factor *Onecut1* (Figure 6L,M, Figure S15A-D). Of note, granule neurons are also highly abundant in the cerebellum ^54^ yet the hippocampus-specific signature, as well as expression levels of *Unc5d* or *Onecut1*, exhibited no age-related change in the bulk data of the cerebellum. Our approach was thus able to identify aging signatures of a given cell type that occur selectively in a specific region.

Finally, we explored whether the biological processes associated with signature genes could indicate differential transcriptional activity across whole pathways or organelles. We observed a significant down-regulation of several mitochondria-related genes in the caudate putamen, including several members of the electron transport chain, which could be indicative of impaired mitochondrial function (Figure 6E). We identified in this region a global, gradual down-regulation of all genes coding for mitochondria-related proteins (Figure S15E), as well as a significant drop in scores for a corresponding mitochondrial signature in aged D2 MSNs, mature oligodendrocytes, and astrocytes (Figure S15F). This was not detected in cell types from the hippocampus or the SVZ (Figure S15F). This specific down-regulation of mitochondrial processes in aged striatum could help to explain previous observations demonstrating selective vulnerability to mitochondrial toxins and stresses in the striatum of old animals ^55, 56^.

In conclusion, we discovered extensive region-specific transcriptional signatures of aging that are largely encoded by expression shifts in distinct neuronal subpopulations reflective of a region’s specialization.

### Rejuvenating interventions act on distinct regions and cell types affected during normal aging

Given the substantial region-specific expression changes during normal aging, we wondered if interventions known to stave off age-related pathologies may also act in a region-specific manner. To this end, we performed region-resolved bulk-seq on brains of 19-month-old mice having experienced either four weeks of acute dietary restriction (aDR), a well established nutritional intervention ^57, 58^, or recurring injections of young mouse plasma (YMP) ^59^, a paradigm to administer circulatory factors found at young age. Both aDR and YMP have previously been shown to exert molecular, structural and cognitive improvements even at this relatively late age ^58, 59^. Understanding the nature and magnitude of the brain’s transcriptional shifts in response to ‘rejuvenating’ interventions, as well as the regions where they occur, may help to decipher the mechanism mediating their effects.

For the dietary intervention, 19-month-old female mice were treated either with four weeks of aDR or *ad libitum* feeding (AL; n = 4-5 female C57BL/6JN per group; Figure 7A). The 25% aDR paradigm (i.e. food reduction to 75% of the AL group) resulted in the expected metabolic shifts, marked by weight loss (Figure S16A,B) and induction of well-recognized expression changes in the liver, albeit to a milder degree than those observed in studies employing chronic DR over years (Figure S16C-E, ^57^). For the young plasma intervention, we profiled the brains of 19-month-old male mice having received recurring injections of either YMP or PBS (n = 3-4 male C57BL/6JN per group; Figure S16F). Critically, the resulting 229 single-region transcriptomes clustered well with the bulk-seq data from the aging cohort (Figure S16G), suggesting a robust sampling of regions across experimental cohorts.

**Figure 7.**
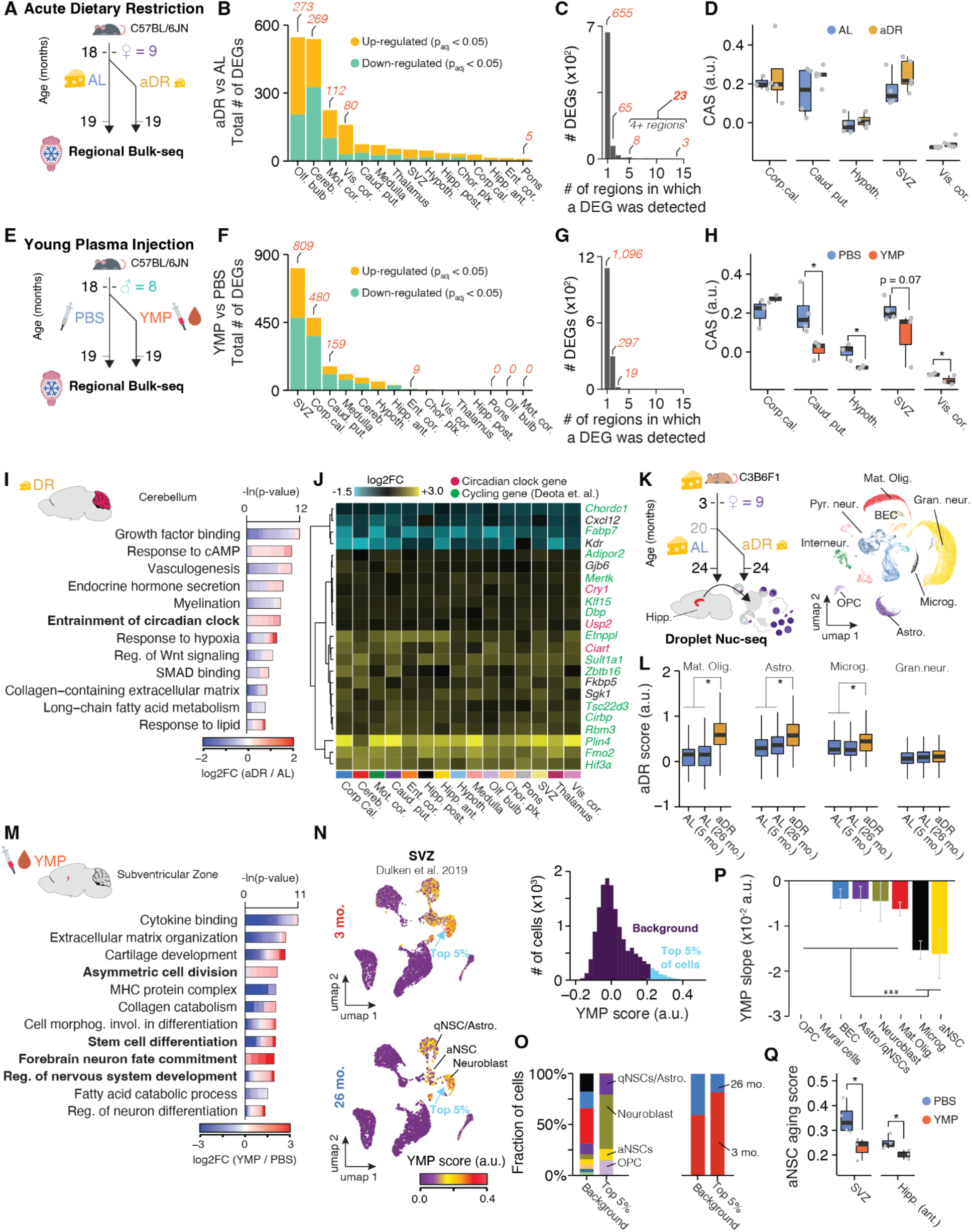
Young plasma injection and acute dietary restriction induce distinct spatial gene expression changes in the aged brain (A) Experiment overview. Aged female mice were either put on acute dietary restriction (aDR) for 5 weeks or maintained on ad libitum (AL) feeding (n = 4-5, 19 months at end of experiment) before whole brains were collected and 15 regions were isolated and subjected to Bulk-seq. (B) Number of differentially expressed genes, split by region, colored by up- and down-regulation. padj < 0.05 Wald-test. List of DEGs can be found in Table S10 (C) Bar graph indicating the number of regions in which a given DEG was detected. (D) CAS shifts in response to aDR across selected regions. P values calculated with one-tailed t-test. Bonferroni correction; padj < 0.05. *** p < 0.001, ** p < 0.01, * p < 0.05. (E) Experiment overview. Aged male mice were injected with either young mouse plasma (YMP) or PBS for 4 weeks (n = 3-4, 19 months at end of experiment) before whole brains were collected and 15 regions were isolated and subjected to Bulk-seq. (F) same as (B) for YMP experiments. (G) same as (C) for YMP experiments. (H) same as (D) for YMP experiments. (I) Representative GO analysis of DEGs with shifts in cerebellum in response to aDR. Lengths of bars represent negative ln-transformed Padj using two-sided Fisher’s exact test. Colors indicate gene-wise log2 fold-changes (log2(FC)) between aDR and ad libitum-fed. The complete list of enriched GO terms can be found in Table S11. (J) Region-wise expression changes in aDR for 24 genes with shifts in at least 4 of the 15 collected regions. (K) Experiment overview. Female C3B6F1 mice were raised on ad libitum diet. At 20 months of age, half of the AL-fed mice were switched to DR (aDR). At 24 months, whole hippocampi were isolated, frozen and Nuc-seq was performed. UMAP representation of all nuclei is depicted (n=69,253 nuclei). (L) Boxplot representation of common aDR scores in four cell types. P values calculated with two-tailed t-test on per-replicate median of score. *** p < 0.001, ** p < 0.01, * p < 0.05. (M) Same as (I) for YMP-induced DEGs in SVZ. The complete list of enriched GO terms can be found in Table S12. (N) UMAP representation of single-cell data of young and old SVZ from (Dulken et al. 2019). Cells are colored with scores for YMP signature (representing DEGs up-regulated in response to YMP). Histogram of score distribution is depicted on the right hand side. The top 5% of cells with the highest scores are indicated. Signature genes can be found in Table S13. (O) Composition of cell types (left) and age (right) groups in cells exhibit the highest YMP scores as compared to all other cells. (P) YMP score slope of linear regressions against age, colored by cell type. Data are mean ± 95% confidence intervals. Two-sided Tukey’s HSD test, adjusted for multiple testing, *** p < 0.001, ** p < 0.01, * p < 0.05. The highest (least significant) Pval is indicated. (Q) Boxplot representation of scores for aNSC aging in SVZ and hippocampus in YMP- or PBS-injected mice. aNSC signature genes can be found in Table S13. P values calculated with two-tailed t-test on per-replicate median of score. *** p < 0.001, ** p < 0.01, * p < 0.05.

Remarkably, aDR and YMP exerted highly distinct expression changes across the brain (Figure 7B-D, F-H). aDR was primarily marked by substantial differential regulation in the olfactory bulb, cerebellum and cortical areas, as well as lesser expression changes across all regions (Figure 7A-C). Interestingly, DEGs under aDR exhibited little overlap with DEGs occurring during aging (Figure S16H-J) and the CAS remained unaffected across the brain (Figures 7D and S16K). In several regions, particularly the cerebellum, we found a strong functional enrichment for aDR-induced genes related to regulation of the circadian clock. Indeed, a set of 23 genes was differentially regulated in at least four regions (Figure 7C,J), which involved three direct members of the circadian clock (*Cry1*, *Usp2* and *Ciart*), as well as 15 genes that have been previously identified to exhibit cycling gene expression in the brain ^60^. We utilized these 23 genes to construct an aDR signature, which was robustly and evenly induced across all brain regions examined (Figure S17A,B). To map out the cell types driving the aDR signature, we performed nuc-seq on whole, frozen hippocampus tissue of 24-month-old female C3B6F1 mice that had been fed AL or subjected to 40% aDR since 20 months of age (Figure 7K; n = 3-4 female C3B6F1 per diet group; cohort previously described in ^57, 61^). Remarkably, the signature was specifically up-regulated under aDR in the same cell types affected by the CAS, namely in mature oligodendrocytes, astrocytes and to a lesser degree in microglia and OPCs, but was unaffected in any neuronal subpopulation (Figure 7L, S17C-E). Together, these results reveal that aDR induces a brain-wide transcriptional program that acts on the same cell types affected by the CAS, albeit through molecular pathways that are orthogonal to those changing during aging.

In contrast to these brain-wide effects of aDR, YMP caused region-selective expression shifts, affecting the SVZ in particular. (Figure 7E-H). Here, we observed profound up-regulation of pathways related to stem cell differentiation and neuronal maturation (Figure 7M). We mapped a signature representing all up-regulated genes under YMP to single-cell data of the SVZ ^27^, where it demarcated neuroblasts, quiescent, and activated neural stem cells (aNSCs), which were primarily found in young mice (Figure 7N,O). SVZ cells from aged mice decreased the YMP signature, and, conversely, age-related DEGs found in aNSCs were down-regulated in YMP-treated mice (Figure 7P,Q). Thus, YMP injection reactivates an expression pattern in the neurogenic lineage that becomes down-regulated with age. In addition to effects on the SVZ, YMP caused significant down-regulation of genes like *C4b*, *B2m*, *Trem2* or *Gfap* in selected regions, and led to a significant CAS reduction in caudate putamen, hypothalamus, SVZ and several cortical areas (Figure S16I,L-M). These findings suggest profoundly different modes of action for aDR and YMP, wherein the latter causes ‘actual’ rejuvenation by reverting molecular shifts occurring during normal aging.

In summary, we uncovered that the rejuvenating interventions aDR and YMP act in distinct, region-specific manners. While aDR instigates a reprogramming of genes related to the circadian clock across all glia, YMP causes a selective reversal of age-related expression signatures, particularly in the neurogenic lineage of the SVZ.

### Aging results in region-specific expression changes of genes associated with human diseases

The region-specific expression level of a gene may not only impact aging of the brain, but susceptibility to disease and selective vulnerability of regional cell populations as well. This impact is the result of region-specific differences at basal levels (i.e. in the young adult brain) and trajectories of age-related changes. We demonstrate this with the example gene *C4b* which is a key genetic risk factor for schizophrenia ^28^. The motor cortex and hippocampus, for example, differ in basal expression of *C4b* at three months of age by up to 3-fold. Meanwhile age-related increases of up to 10-fold were observed from the least affected visual cortex to the corpus callosum (Fig S9). Assuming that region-specific differences in gene expression and aging trajectories exist in human brains, it is reasonable to assume this would influence the pathogenesis and clinical manifestations of a given disease.

To explore this concept in the context of neurodegenerative diseases such as AD or PD, we analyzed the expression of genes linked to autosomal dominant forms of disease or genes linked to the risk of developing sporadic forms of disease. To this end, we ranked regions based on the highly variable expression of either *Apoe*, *Trem2*, *Plcg2* or *Scna* (α-synuclein) at 3 months (Figure 8A-D). Crucially, the expression distribution across the brain for each gene became substantially rearranged in old animals as a consequence of region-specific differential regulation. To systematically assess the regulation of genes linked with disease risk, we assembled lists of Genome-wide association studies (GWAS) genes for AD or PD, and investigated whether they were significantly enriched among age-related DEGs of a given region (Table S14) ^62, 63^. We further clustered regions that share differentially expressed disease-associated genes, allowing us to find anatomical ‘hubs’. Interestingly, each of the disease-associated gene sets exhibited a different enrichment pattern (Figure 8E-G) and a varying number of associated genes (Figure 8H-I). GWAS hits for AD, including *Apoe*, *Ms4a6d* and *Plcg2* ^64^, were part of DEGs that were co-regulated in a small cluster of three regions: the choroid plexus, corpus callosum, and pons, suggesting a different region-specific regulation of these genes as a potential modulator of disease presentation (Figure 8E). In contrast, PD-related genes, like the neuroprotective gene *Ip6k2* ^65^, were not concentrated within a cluster of regions but rather distributed across the choroid plexus, cerebellum, SVZ, and visual cortex with limited overlap (Figure 8F). This pattern indicates a disperse regulation of disease-associated genes among the regions studied, which can add to our understanding of how and where PD may progress. Of note, the substantia nigra, a major region where PD typically manifests, was not quantified in our study.

**Figure 8.**
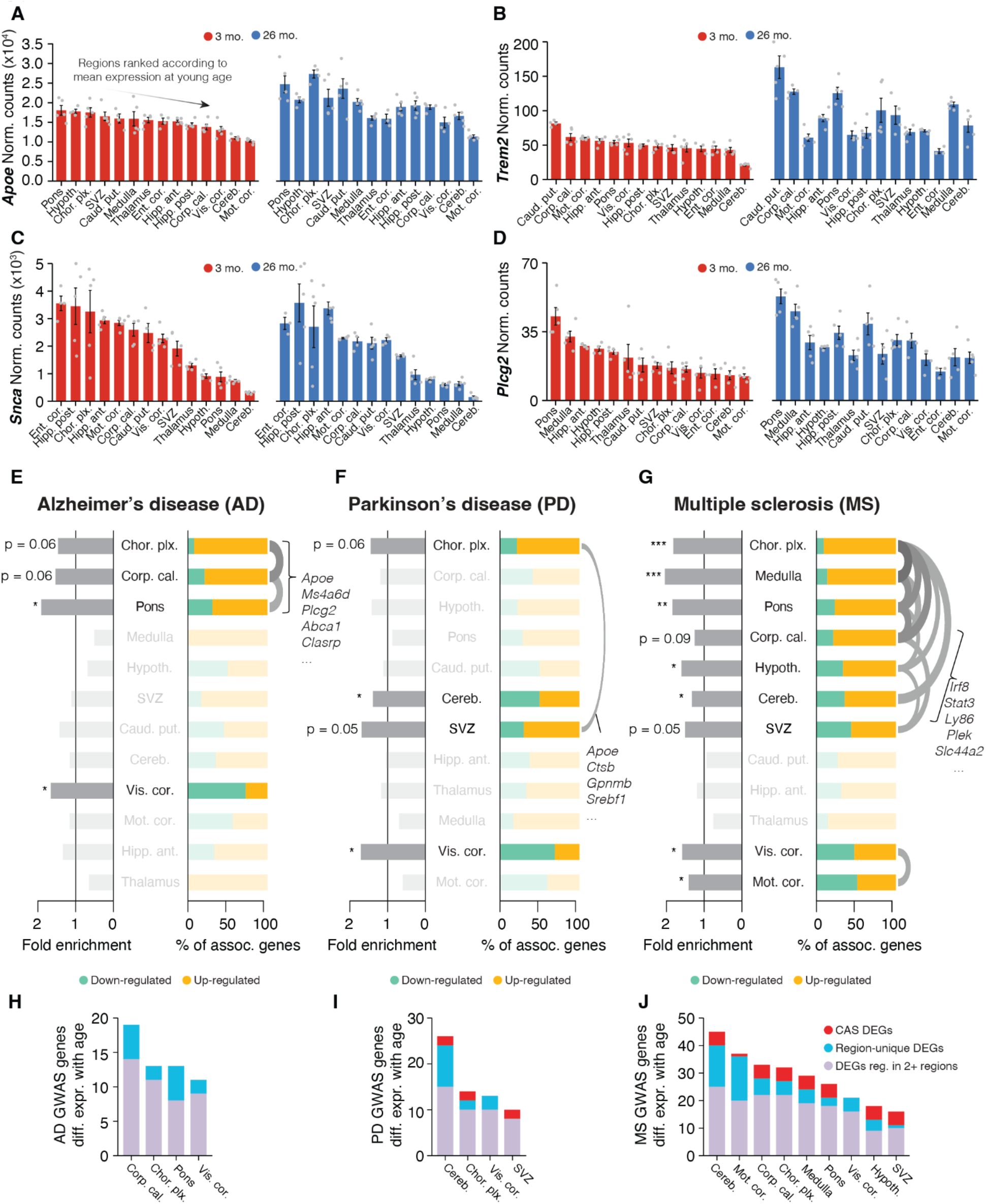
Interplay of region and age shapes expression of disease variant homologues (A-D) Bulk expression for (A) *Apoe*, (B) *Trem2*, (C) *Scna* (α-synuclein) and (D) *Plcg2* at 3 and 26 months of age. Only male samples are displayed. Regions are ordered according to mean expression at young age in descending order. Data are mean ± s.e.m. (E-G) Enrichment analysis of region-resolved DEGs for human GWAS variants for (E) AD, (F) PD and (G) MS. Associated genes are listed in Table S14. Fold enrichment (left bars) and the relative composition of disease-associated DEGs with respect to their regulation (right bars) is indicated. Regions with no significant enrichment are transparent. The vertical order of regions results from hierarchical clustering on a pairwise Jacquard Distance matrix, so regions with overlapping DEGs will cluster together. Gene overlaps with a Jaccard index ≥ 0.25 are indicated with an arc. One-sided Fisher’s exact test with hypergeometric distribution, Benjamini–Hochberg correction. *** p < 0.001, ** p < 0.01, * p < 0.05. (H-J) Number of DEGs per region that are homologues of human GWAS variant for (H) AD, (I) PD and (J) MS. Colors group the genes into CAS DEGs, region-specific DEGs or other (DEG in 2 or more but less than 10 regions).

Given the age-related effects observed in white matter-rich regions, we also analyzed GWAS genes for multiple sclerosis (MS), exhibiting significant associations with DEGs from nine different regions that fell into two clusters, indicating two disparate subsets. One cluster consisted of several regions, including the corpus callosum and cerebellum, that up-regulated a shared set of inflammation-related genes such as *Stat3*, *Ly86* and *Irf8*, all of which were also part of the CAS (Figure 8G). This raises the intriguing possibility of similarities between the pathophysiology of inflammation and demyelination associated with MS and the accelerated aging observed in white matter-rich areas. The visual and motor cortex regions formed the second cluster, exhibiting even numbers of up- and down-regulated MS genes. This supports recent evidence indicating transcriptional shifts (e.g. of *Cbln2*) in cortical areas that can occur far away from the actual lesions ^66^ and highlights the need to broadly study regional patterns of gene expression to understand the role of MS-associated genes. To better contextualize the biological relevance of the observed expression changes, we further compared the up-regulation of GWAS homologues with their baseline expression across regions (Figure S18B,C; STAR Methods), which confirmed that age-related differential expression in specific regions led to a significant redistribution of where in the brain a given GWAS homologue was predominantly expressed (Figure S18C-E).

Taken together, our data demonstrate that genetic risk factors linked to three major neurological and neurodegenerative diseases are affected by age in a region-selective manner. While we cannot predict whether the directionality of the regulation itself has a biological consequence, we consider that the region-specific differential regulation of such genes could be an additional factor modulating disease risk.

## Discussion

We report here a comprehensive spatiotemporal map of gene expression across the mouse brain throughout the adult lifespan and in response to rejuvenating interventions, consisting of 1,076 bulk, 16,277 spatial spot and 150,869 single-cell transcriptomes – supported by several publicly available bulk-spatial- and scRNA-seq datasets. We find that aging affects regional transcriptomes with widely varying magnitude and timing, including the expression of risk genes for neurodegenerative disorders. Gene expression shifts during aging fall into two general categories: a single set of 82 genes that reflects shared aging dynamics of glia cells with spatially-defined magnitude, and a dozen gene sets with region-specific activity encoded in specialized neuronal subpopulations. We establish a CAS for each mouse and region, analyzing its trajectory and rate of change over time as a quantitative measure of a region’s transcriptional age. Across analyses, white matter and white matter-rich regions emerge as the most transcriptionally impacted areas during aging. It will be particularly interesting to explore how these changes relate to developmental patterns of myelination and brain function, and whether susceptibility to brain aging and dysfunction are related to developmental processes. The advent of single-cell technologies and cell dissociation methods have enabled the exploration of an ever increasing number of cell populations in the brain ^67^, which in turn allows for cell type-specific characterization of gene expression during aging ^9^. The interplay between cell type and regional niche during aging is, however, yet to be more deeply understood. Our results emphasize the importance of region identity as a profound modulator of gene expression dynamics in the context of aging and neurodegeneration. It will be important for future studies to examine if these heterogeneous expression patterns result in corresponding shifts of the proteome or downstream functional changes in neuronal activity and plasticity. Given comparable observations that astrocytes exhibit stronger age-related expression changes in striatum and cerebellum as compared to cortical areas ^32, 68, 69^, it is likely that other glial cells contribute to the heterogeneous CAS increase on the bulk level. We thus propose further exploration of the CAS in other non-neuronal cell types. This may also help to clarify if microglia are active drivers of the regional expression dynamics described here, or rather respond to cues provided by other cell type(s) in the region.

Our data reveal that certain brain regions are selectively vulnerable to aging, with the white matter fiber tracts exhibiting a particular sensitivity. These areas are dense with myelinated axons and myelinating cells, forming the basis of neurotransmission across brain regions ^70^. The strong activation of immune- and inflammation-related genes, as well as differential expression of remyelination regulators like *Gpr17* ^71, 72^, suggest that the homeostasis of this region is compromised at old age. This could perturb myelin sheath integrity and potentially impair axonal signal transmission between regions as an early event in brain aging. In line with this hypothesis, rejuvenation of hippocampal oligodendrogenesis in aged mice via injection of young cerebrospinal fluid (CSF) improves long-term memory consolidation, thus demonstrating a causal role of compromised myelin on cognition^73^.

In line with this, we have demonstrated that the region-specific transcriptome atlas generated here can form a basis for testing rejuvenation strategies like dietary restriction and young plasma injection to quantify their spatiotemporal impact on the brain at the molecular level. Specifically, we found evidence that aDR induces brain-wide reprogramming of genes associated with circadian rhythmicity, an observation that agrees with recent findings that the lifespan-extending effects of DR depend on a shift in circadian rhythm ^74–76^. The DR expression pattern of circadian reprogramming found here appears to be independent of the timing of feeding, as we found the signature in two separate cohorts of mice treated with differing aDR regimens and feeding times (C57BL/6JN were fed in the afternoon and C3B6F1 fed in the morning ^57^). Future work should investigate if and how altered circadian rhythmicity could affect cell function, and elucidate why only glia but not neurons are affected. In contrast to aDR, YMP appeared to directly reverse age-related expression shifts, particularly in the SVZ, corpus callosum and caudate putamen. Interestingly, all three regions are proximal to the ventricles, which are highly permissive for peripheral plasma proteins^77^, which could explain their sensitivity to YMP. Decline of neurogenic capacity is a known feature of aging in mice and humans, and heterochronic parabiosis or injection of aged plasma into young individuals inhibits neurogenesis ^78^. Conversely, we provide evidence that YMP specifically reverses aging signatures of the neurogenic lineage, which indicates restoration of adult neurogenesis that should be assessed specifically with cell cycling tracing assays.

Our findings strongly support the notion that the impacts of aging on brain function are region specific, potentially relating to the regional vulnerability across different diseases as well as the varied manifestations of neurodegeneration at the level of an individual. We demonstrate that key genetic risk genes are differentially expressed in a region-specific manner, which could locally amplify, or attenuate their impact on disease pathways. Importantly, our findings also suggest that aging may drive dysfunction in brain regions that are not predominantly affected and studied by classical pathological hallmarks, highlighted by AD risk genes including *Apoe*, *Ms4a6d*, *Plcg2* and *Trem2* which are dysregulated with aging in the mouse choroid plexus, corpus callosum and pons with aging. The translation of these findings to humans may serve as a new brain cartography leading to novel treatment strategies and interventions.

## Limitations of this study

Because region-specific and age-related changes in gene expression may be distinct for each species, the conclusions drawn here from mouse data may not necessarily be translatable in their entirety to humans. Additionally, the analysis of our single-nuclei RNA-seq data computationally pools animals of the same age and cell type, and thus cannot be analyzed in a replicate-sensitive manner. We also combined sexes for most of our analyses, potentially masking subtle sex-specific gene expression differences. Relatedly, limitations in mouse availability for this study resulted in the two oldest ages being profiled only in male mice, and thus did not capture differences in very late life expression shifts between males and females. We suggest further interrogation of potential sex differences of murine brain aging late in life. Lastly, determining the exact distribution and composition of major cell type classes remains a challenge in the field, preventing us from fully eliminating the possibility that baseline differences in abundance play some role in detected aging effects.

## STAR METHODS

### RESOURCE AVAILABILITY

#### Lead Contact

Requests for resources and reagents should be directed to the lead contact, Tony Wyss-Coray.

#### Data and Code Availability

The sequencing datasets analyzed during the current study are available in the Gene Expression Omnibus repository under accession numbers GSE212336, GSE212576, GSE212903, GSE227689 and GSE227515.

##### (i)#Scripts detailing the core analyses are stored on GitHub: https://github.com/OliInTheValley/SpatioTemporal_Analysis

### EXPERIMENTAL MODEL AND SUBJECT DETAILS

#### Animal husbandry and organ collection

For Bulk-seq and nuc-seq, male and female C57BL/6JN mice were shipped from the National Institute on Aging colony at Charles River. 5-6 male and 5 female mice were used for each 3, 12, 15, 18, and 21 months group, while only 5 and 3 male mice were used for the 26 and 28 months groups, respectively. For the 10X Visium experiments, aged C57BL/6J mice (000664, Jackson Laboratory) were shipped from Jackson Laboratory. 2 male mice/age were used for the 6, 18, and 21 months groups. All mice of the aging and aDR cohorts were housed in cages of 2-3 mice at the Stanford ChEM-H animal facility under a 12 h/12 h light/dark cycle at 67–73 °F and provided with food and water ad libitum. Mice were housed in the ChEM-H animal facility for one month before euthanasia, except for mice older than 18 months, which were housed at the ChEM-H animal facility beginning at 18 months. Takedown of the bulk- and nuc-seq cohort was conducted between 10:00am-12:00pm over four days. Takedown of mice for 10X Visium was conducted between 10:00am-10:15am on a single day. Age groups and sexes were rotated through over the duration of the takedowns to average out the impact of takedown time and day. After anaesthetization with 2.5% v/v Avertin, ∼700ul blood was drawn via cardiac puncture before transcardial perfusion with 20 ml cold PBS. The brain was immediately removed and snap-frozen by submerging for 60 seconds in liquid nitrogen-cooled isopentane. Brains were stored at -80°C until further processing.

All animal care and procedures complied with the Animal Welfare Act and were in accordance with institutional guidelines and approved by the institutional administrative panel of laboratory animal care at Stanford University.

For the aDR study with C57BL/6JN mice, 18-months-old mice were randomly assigned to AL or aDR. aDR treatment was initiated by transferring mice from AL to 10% aDR for 7 days. After that, aDR was increased to 25%. aDR animals were fed once per day between 3-5 p.m., and all animals were checked daily for their well-being and any deaths. For the first 16 days, weights were checked daily. Mice were euthanized at the ages of 19 months. All mice were euthanized in the morning within a period of 6 hours prior to the regular feeding time of the DR mice.

The aDR study with C3B6F1 mice was performed in accordance with the recommendations and guidelines of the Federation of the European Laboratory Animal Science Association (FELASA), with all protocols approved by the Landesamt für Natur, Umwelt und Verbraucherschutz, Nordrhein-Westfalen, Germany (84-02.04.2015.A437). Female F1 hybrid mice (C3B6F1) were generated in-house by crossing C3H/HeOuJ females with C57BL/6NCrl males (strain codes 626 and 027, respectively, Charles River Laboratories). Lifespans of chronic DR and AL C3B6F1 mice were previously published ^61^. Pups were weaned at 3–4 weeks of age and were randomly assigned to cages upon weaning. Animals were housed in groups of 5 females in individually ventilated cages under specific-pathogen-free conditions with constant temperature (21 °C), 50–60% humidity and a 12 h/12 h light/dark cycle. For environmental enrichment, mice had constant access to nesting material and chew sticks. All mice received commercially available rodent chow (ssniff R/M-Low phytoestrogen, ssniff Spezialdiäten, Germany) and were provided with filtered water ad libitum. aDR animals received 60% of the food amount consumed by AL animals. aDR treatment was initiated at 20 months of age by directly transferring mice from AL to 40% DR. aDR animals were fed once per day, and all animals were checked daily for their well-being and any deaths. Mice were euthanized at the ages of 24 months. All mice were euthanized in the morning within a period of 3 hours prior to the regular feeding time of the DR mice. Mice were euthanized by cervical dislocation, and tissues were rapidly collected and snap-frozen in liquid nitrogen.

The cohort of mice treated with YMP or PBS were housed at the Palo Alto VA animal facility under a 12 h/12 h light/dark cycle at 68–73 °F under 40–60% humidity. All experiments were performed in accordance with institutional guidelines approved by the VA Palo Alto Committee on Animal Research. Euthanasia and organ collection was conducted in the same way as the aging cohorts.

### METHOD DETAILS

#### Processing and administration of plasma

Young Mouse Plasma (YMP) was collected following the protocol described by^59, 79^. Briefly, C57Bl/6J male mice aged 2 months were group-housed and anesthetized with 2.5% v/v Avertin. Approximately 700 μl of blood was drawn via cardiac puncture prior to transcardial perfusion. Blood was collected using 15 μl of 250 mM EDTA (Thermo Fisher Scientific, 15575020) and centrifuged at 4°C for 15 minutes at 1,000g to obtain plasma. The plasma from 20-25 mice was pooled together and dialyzed in 1X PBS using cassettes (Slide-A-Lyzer Dialysis Cassettes, 3.5 kDa molecular weight cut-off, 3-12 ml) before being frozen at -80°C.

For plasma transfer experiments, male C57BL/6JN mice aged 18 months were injected retro-orbitally with 150 μl of YMP per injection. Prior to injection, mice were habituated by being placed on the procedure table in their cage. Injections were administered every 3-4 days, alternating between the left and right eye to allow for recovery. Mice were rested for four days before tissue collection.

#### Brain region dissection

Dissociating the mouse brain at scale poses several challenges, as the tissue consists of a multitude of biologically distinct structures that require careful, time-consuming separation to avoid cross-region contamination – all while avoiding tissue degradation and loss of RNA quality. We systematically assessed several isolation, dissection, and freezing strategies, most of which yielded low-quality RNA or were not scalable to the intended set of samples and regions. We found success in perfusing the animal before isolating and freezing the whole brain in under 5 minutes, thus rapidly stabilizing the tissue and RNA. Region isolation via slicing and atlas-guided tissue punching was subsequently conducted at sub-0C temperatures. In detail, brain regions were dissected from frozen mouse brains through a modification of a previously developed protocol ^80^. Frozen brains were sliced into 1mm thick coronal slices at -20°C using a metal brain matrix and .22mm razor blades (Ted Pella, 15045; VWR, 55411-050) and were then placed on dry ice and covered to prevent condensation. One slice at a time was placed on a metal block cooled on wet ice and 1.5mm and 2mm diameter regions of interest were dissected quickly via disposable biopsy punches (Alimed, 98PUN6-2, 98PUN6-3) from the left and right hemispheres guided by visual landmarks and the Allen Mouse Brain Atlas. The same biopsy punch was used for identical regions between left and right hemispheres, but replaced between regions and mice. 15 regions were collected: three cortical regions (motor cortex, visual cortex and entorhinal cortex), anterior (dorsal) and posterior (ventral) hippocampus, hypothalamus, thalamus, caudate putamen (part of the striatum), pons, medulla, cerebellum and the olfactory bulb, corpus callosum, choroid plexus and the subventricular zone. The following regions required overlapping punches and were thus sequentially collected: (1) motor cortex, (2) caudate putamen, (3) subventricular zone, (4) corpus callosum.

Regions were stored at -80°C until further processing.

#### Bulk-seq preparation and sequencing

We isolated RNA from the right hemisphere brain regions described above using the RNeasy 96 kit (Qiagen, 74181) and a TissueLyser II (Qiagen, 85300), according to RNeasy 96 Handbook protocol “Purification of Total RNA from Animal Tissues using Spin Technology” without the optional on-plate DNase digestion. Quality control of RNA was conducted using a Bioanalyzer (Agilent) at the Stanford Protein and Nucleic Acid Facility for three randomly selected samples per brain region.

cDNA and library syntheses were performed in house using the Smart-seq2 protocol as previously described ^8, 79^ with the following modifications: Extracted RNA (2 ul at a concentration of 25 ng/ul) was reverse-transcribed and the resulting cDNA amplified using 10 cycles. After bead clean-up using 0.7x ratio with AMPure beads (Thermo Fisher, A63881), cDNA concentration was measured using the Quant-iT dsDNA HS kit (Thermo Fisher, Q33120) and normalized to 0.4 ng/ul as input for library prep. 0.8 ul of each normalized sample was mixed with 2.4 ul of tagmentation mix containing Tn5 Tagmentation enzyme (20034198, Illumina) and then incubated at 55°C for 12 minutes. The reaction was stopped by burying the plate in ice for 2 minutes followed by quenching with 0.8 ul 0.1% sodium dodecyl sulfate (Teknova, S0180). 1.6 ul indexing primer (IDT) was added and amplified using 12 cycles. Libraries were pooled and purified using two purification rounds with a ratio of 0.8x and 0.7x AMPure beads. Library quantity and quality was assessed using a Bioanalyzer (Agilent) and Qubit dsDNA HS kit. Pipetting steps were performed using the liquid-handling robots Dragonfly or Mosquito HV (SPT Labtech) using 384 well-plates and PCR reactions were carried out on a 384-plate Thermal Cycler (BioRad). Illumina sequencing of the resulting libraries was performed by Novogene (https://en.novogene.com/) on an Illumina NovaSeq S4 (Illumina). Base calling, demultiplexing, and generation of FastQ files were conducted by Novogene.

#### 10X Visium preparation and sequencing

Frozen brains (n = 2 males per age; aged 6, 18 and 21 months; C57BL/6J strain) were embedded in OCT for cryosectioning at 16 micron thickness (app. Bregma -1.655mm; Allen brain reference atlas coronal section 71). Reactions were carried out with the Visum Spatial Gene Expression (GEX) and Tissue Optimization (TO) Slide & Reagent Kits according to the manufacturer’s protocol with recommended reagents (10X Genomics, 1000193 and 1000184). Sections were placed on designated capture areas of slides for TO and GEX and stored at -80°C until further processing. TO and GEX slides were fixed with methanol and stained with hematoxylin and eosin (H&E) for visualization of tissue morphology on a AxioImager Widefield Fluorescence Microscope (Zeiss) at 10-fold magnification. To determine the optimal permeabilization time, TO slides were incubated with permeabilization enzyme for various timeframes followed by incubation with reverse transcriptase (RT) and fluorescently labeled nucleotides (FLNs) and enzymatic tissue removal. After visualizing cDNA signal via fluorescence microscopy, we selected 20 minutes as the optimal permeabilization time. GEX slides were incubated with permeabilization enzyme for 20 minutes followed by incubation with RT. cDNA was then transferred into tubes and amplified for 15 cycles using a Thermal Cycler (BioRad). Library construction steps were performed according to the manufacturer’s protocol and included cDNA fragmentation, end repair and A-tailing, adaptor ligation, and sample indexing and amplification. Quality control of the constructed library was conducted via Bioanalyzer (Agilent). Illumina sequencing of the resulting libraries was performed by Novogene (https://en.novogene.com/) on an Illumina NovaSeq S4 (Illumina). Base calling, demultiplexing, and generation of FastQ files were conducted by Novogene.

#### Nuc-seq preparation and sequencing

Single-nuclei preparation (n = 2 males and females per age and region; aged 3 and 21 months; all C57BL/6JN strain) and sequencing was performed as previously described^29^ with the following modifications: Nuclei from left hemisphere brain region punches were isolated with EZ Prep lysis buffer (Sigma, NUC-101) on ice. Single-nuclei isolation from the whole hippocampus of C3B6F1 mice (aDR) was performed similarly with the exception that tissues were not pooled. Samples were placed into 2 ml cold EZ lysis buffer in a 2 ml glass dounce tissue grinder (Sigma, D8938) and homogenized by hand 25 times with pestle A followed by 25 times with pestle B while incorporating a 180-degree twist. Tissue homogenate was transferred to a fresh 15 ml tube on ice. The tissue grinder was rinsed with 2 ml fresh lysis buffer and transferred to the tube holding the homogenate for a total volume of 4 ml. Samples were incubated on ice for 5 minutes. Nuclei were centrifuged at 500 x g for 5 minutes at 4°C, supernatant removed and pellet resuspended with 4 ml EZ lysis buffer, and incubated on ice for 5 minutes. Centrifugation at 500 x g for 5 minutes at 4°C was repeated. After removing supernatant, the pellet was resuspended with 4 ml chilled PBS and filtered through a 35-um cell strainer into a 5 ml round bottom FACS tube (Corning, 352235). Following centrifugation at 300 x g for 10 minutes at 4°C with break 3, supernatant was gently poured out leaving behind the nuclei pellet. Pellet was resuspended in 400 ul PBS containing 1% BSA (Thermo Fisher, BP9700100), 0.2 ul Hoechst dye (Thermo Fisher, H3570), and 2 ul recombinant RNase inhibitor (Takara, 2313B). Isolated nuclei were sorted on a MA900 Multi-Application Cell Sorter (Sony Biotechnology). 25,000 single nuclei per sample were collected into 1.5 ml DNA lo-bind tubes (Eppendorf, 022431021) containing 1 ml buffer mix with PBS, UltraPure BSA (Thermo Fisher, AM2618), and RNase inhibitor (Takara, 2313B). One male and one female sample from the same time point and region were pooled at this stage by FACS collecting into the same sample tube (thus yielding 50,000 nuclei per tube). Collected nuclei were centrifuged at 400 x g for 5 minutes at 4°C with break 2. Supernatant was removed leaving 40 ul suspended nuclei. Nuclei were counted using a hemocytometer (Sigma, Z359629-1EA) and assessed for concentration and quality.

Reagents of the Chromium Single Cell 3’ GEM & Gel Bead Kit v3.1 (10X Genomics, 1000121) were thawed and prepared according to the manufacturer’s protocol. Nuclei and master mix solution was adjusted to target 10,000 nuclei per sample and loaded on a standard Chromium Controller (10X Genomics, 1000204) according to manufacturer protocols. We applied 11 PCR cycles to generate cDNA. Library construction was conducted using Chromium Single Cell 3’ Library Construction Kit v3 (10X Genomics, 1000121). All reaction and quality control steps were carried out according to the manufacturer’s protocol and with recommended reagents, consumables, and instruments. We chose 11 PCR cycles for library generation. Quality control of cDNA and libraries was conducted using a Bioanalyzer (Agilent) at the Stanford Protein and Nucleic Acid Facility. Illumina sequencing of the resulting libraries was performed by Novogene (https://en.novogene.com/) on an Illumina NovaSeq S4 (Illumina). Base calling, demultiplexing, and generation of FastQ files were conducted by Novogene.

### QUANTIFICATION AND STATISTICAL ANALYSIS

#### Bulk-seq quantification, quality control

Raw sequence reads were trimmed to remove adaptor contamination and poor-quality reads using Trim Galore! (v.0.4.4, parameters: --paired --length 20 --phred33 --q 30). Trimmed sequences were aligned using STAR (v.2.5.3, default parameters). Multi-mapped reads were filtered. Read quality control and counting were performed using SeqMonk v.1.48.0 and RStudio v.3.6. Data visualization and analysis were performed using custom Rstudio scripts and the following Bioconductor packages: Deseq2 ^33^, topGO, destiny and org.Mm.eg.db. Finally, we excluded pseudogenes and predicted genes from the count matrix to focus predominantly on well-annotated, protein-coding genes. In total, all of the following analyses were performed on the same set of 21,076 genes.

To assess the quality of our dataset, the count matrix was analyzed using Seurat’s built-in, default dimensionality reduction workflow ^81^ (Normalization: ‘LogNormalize’; Variable feature discovery: selection.method=’vst’, features=2000). Umaps were calculated using Seurat’s built-in functions, based on the first 40 principle components (PC) dimensions (Figure 1C; Figure S2B-D). A shared-nearest-neighbors graph was constructed using the first 40 PC dimensions before clustering samples using Seurat’s built-in FindClusters function with a resolution of 0.8 to identify samples that would not cluster with their region of origin.

We corroborated the Seurat-based quality assessment by loading and normalizing the count matrix using DEseq2 before conducting the built-in variance stabilizing transformation ^33^. We then performed hierarchical sample-to-sample clustering using Ward’s clustering algorithm across all 21,076 genes (Figure S2E). To detect whether samples within a given tissue would show profound clustering by age, we finally calculated diffusion maps using the R package destiny with default parameters (Figure 1D).

For bar graph visualization of gene expression (e.g. Figure 1E), we used DEseq2-normalized counts after calculating factors and dispersion estimates across all regions using the factor design ∼age + region. Trajectories were smoothed via triangular moving average across the interval between 3 and 28 months. This quantification and smoothing was solely used for visualization and was not the basis for any statistical testing in this study.

#### Bulk-seq differential expression

To identify significant differential expression changes with age, we used the raw count matrix as recommended for the DEseq2 standard analysis pipeline. Factors and dispersion estimates were calculated for each region separately. We conducted differential expression analysis comparing samples from 3 months to each consecutive time point, using sex as covariate. This is consistent with previously published differential expression analyses performed across whole organs in mice^8^. *P* values were adjusted for multiple testing, and genes with an adjusted *P* value of less than 0.05 were determined to be statistically significant. Finally, we required a gene to reach statistical significance (after multiple testing correction) in at least 2 pairwise comparisons (e.g. 3 months vs 18 months and 3 months vs 21 months) to be called a differentially expressed gene (DEG). We chose this criterion to retain only genes with robust differential expression patterns across age groups. We recognize that this tends to select against genes that are differentially expressed very late in life (i.e. 3 months vs 28 months). To demonstrate the validity of using sex as a model covariate in the differential gene expression analysis, we performed gene-wise likelihood-ratio tests (LRT analysis, as implemented in DESeq2 7). This assesses the goodness of fit between a ‘complete’ model formula (expr. ∼ age + sex + age:sex interaction) and the model formula implemented in our study (expr. ∼ age + sex). This analysis was run across ages 3 to 21 months, due to the lack of female samples for ages 26 and 28 months. If aging trajectories would be reasonably similar between sexes, then the LRT would indicate a significantly better goodness of fit for the complete model in only very few genes, if any (i.e. the interaction term improves the fit). In addition, we repeated the differential expression analysis for the age groups 3 to 21 months, for which we had data from both sexes.

Differential expression for the rejuvenation interventions was performed as described for the aging cohorts, except that no additional filter of 2 pairwise comparisons was employed.

#### DEG Gene Ontology functional enrichment

Unless stated otherwise, we performed functional enrichment analysis for DEGs using the Biocunductor package topGO as described in detail before ^8, 57^. Unless stated otherwise, the set of expressed genes (defined as passing the independent filtering criterion of DEseq2 ^33^) was used as background for all functional enrichment analyses involving expression data. Top-ranked, representative Gene Ontology (GO) terms were selected and visualized using the CellPlot package. The full-length GO terms were shortened to fit into the figure format.

#### Bulk-seq GWAS gene enrichment and expression distribution analysis

We analyzed if DEGs of a given region would be enriched for disease-associated genes using a previously assembled list of GWAS hits for several neurodegenerative diseases ^63^. The analysis was focused on Alzheimer’s disease and Parkinson’s disease (both age-related forms of dementia) and multiple sclerosis as we had observed several white matter-related effects in our dataset. We refer to these as ‘disease-associated genes’. Disease-associated genes that were expressed in a given region (defined as passing the independent filtering criterion of DESeq2 ^33^) were analyzed. To determine if disease-associated genes were enriched among the DEGs of a given region, we used a one-sided hypergeometric test with expressed genes as background. Resulting *P* values were corrected for multiple testing. We chose the anterior hippocampus region as representative for the hippocampus, and further excluded the entorhinal cortex (too few DEGs) and olfactory bulb. For each disease, we plotted the enrichment and the relative composition of disease-associated DEGs with respect to their regulation (i.e., up- or down-regulated) using the CellPlot package. We clustered regions for using a pairwise Jacquard Distance matrix, so that regions with overlapping diseases-associated DEGs will cluster together. Gene overlaps with a Jaccard index ≥ 0.25 were indicated with an arc.

Additionally, we performed a systematic analysis of expression shifts that could affect a given GWAS homologue’s distribution across the brain. To this end, we focused on sets of GWAS-DEGs in each region and ranked their mean expression at young (3 months) and very old (26 months) age, respectively. For instance, *Irf8*, one of the GWAS-DEGs associated with MS, became differentially regulated in multiple regions, including the corpus callosum (FigS18 B). However, the actual rank of the top *Irf8*-expressing regions (caudate putamen, SVZ and corpus callosum) stayed relatively constant . In contrast, the expression of *Nlrc5*, a gene from the same set, was only the 8th-highest in the corpus callosum at young age, but became the highest across all tissues with age (FigS18 B). Here, the differential regulation with age led to a significant redistribution of where the gene is predominantly expressed. Expanding the analysis to all GWAS-DEGs in a given tissue/disease set with least 15 genes to focus on (to ensure statistical power), we tested if there would be a systematic shift in rank-based expression using paired two-sided Wilcoxon rank-sum tests.

#### Bulk-seq correlation of gene expression with age

For each region separately, we probed the expression of each gene (using DEseq2-normalized counts) for positive or negative correlation with age using Spearman’s method and tested for significant association. *P* values were adjusted for multiple testing using the Benjamini-Hochberg method. Genes with Spearman’s rho ≥ 0.5 or ≤ -0.5, respectively, and padj ≤ 0.05 were called as significantly age-correlated in a given region. The total number of age-correlated genes was used to evaluate the impact of aging on a given region.

The sex-specific age-correlations in the hypothalamus were performed similarly, by subsetting the dataset to the ages 3 to 21 months (for which data of both sexes was available). Correlation with age was then calculated for each gene based on the male or female samples only. Criterion for age-correlated genes remained the same.

#### Weighted gene co-expression network analysis (WGCNA)

Network analysis was performed with the Weighted Gene Correlation Network Analysis (WGCNA) ^82^ package to identify significant modules that were associated with a specific aging group and brain region. Modules were independently detected in each brain region. For each brain region the soft-thresholding (ß value) was set based on scale-free topology (R2>0.8) to construct a correlation adjacency matrix. ß values 18, 10, 9, 8, 12, 4, 4, 5, 7, 9, 24, 14, 4 and 13 were used for the corpus callosum, cerebellum, motor cortex, entorhinal cortex, anterior hippocampus, posterior hippocampus, hypothalamus, medulla, olfactory bulb, choroid plexus, pons, SVZ, thalamus and visual cortex respectively. The ‘blockwiseModules’ function was used to construct the network. Biweight midcorrelation (‘bicor’) was used to compute the correlation between each pair of genes. Network analysis was performed with the “signed” network. The “deepSplit” argument value was 2 and a minimum cluster size was 25. (blockwiseModules parameters: datExpr=(datExpr), maxBlockSize=22000, networkType=“signed”, corType=“bicor”, power=ß, saveTOMFileBase=(file=’TOM_signed’), minModuleSize=25, deepSplit=2, saveTOMs=TRUE). The average linkage hierarchical clustering of the topological overlap dissimilarity matrix (1-TOM) was used to generate the network dendrogram. The hybrid dynamic tree-cutting method was used to define modules. Modules were summarized by their first principal component (ME, module eigengene) and modules with eigengene correlations >0.9 were merged.

Module-aging group associations were evaluated using a linear model within each brain region. Significance values were corrected for multiple testing using Benjamini-Hochberg method. Results from module-eigengene association tests are shown in Table S4. Genes within each module were prioritized based on their module membership (kME), defined as correlation to the module eigengene. The top ‘hub’ genes for several of the modules are shown in supplementary Table S4. Cell type enrichment analyses were performed using several mouse derived cell type specific expression datasets ^83–85^. Enrichment was performed for cell type specific marker genes using Fisher’s exact test, followed by Benjamini-Hochberg-correction for multiple testing. The WGCNA results were assembled in summarizing figures that can be browsed through our interactive shiny app website (https://twc-stanford.shinyapps.io/spatiotemporal_brain_map/).

#### Estimating the variance of the data depending on metadata

To estimate the variance in the data depending on age, tissue or gender we made use of principal variance component analysis (PVCA) as implemented in the Bioconductor Package pvca and described in detail in ^8^. PVCA combines the strength of principal component analysis and variance components analysis (VCA). Originally it was applied to quantify batch effects in microarray data. In our case, however, we do not provide experimental batches but rather groups of meta data as input

#### Gene signature generation and score calculation

Gene *signatures* are used in this study to quantify the expression of a gene set, thus representing the aggregated expression of multiple genes in a given transcriptome (e.g. a regional bulk-seq transcriptome or a single-nuclei transcriptome). The resulting value is defined as a *score*. Throughout the manuscript we generated signatures and quantified scores using the VISION (v.3.0) package as detailed in the original study ^25^. Notably, VISION z-normalizes signature scores with random gene signatures to account for global sample-level metrics (such as total number of counts/UMIs, which can be affected by age ^86^). While VISION was originally intended for the analysis of signatures in single-cell data we found its analysis workflow applicable for bulk, spatial and single-cell/-nuclei datasets. We note that due to differences in baseline expression across regions or cell types as well as the z-normalization mentioned above, VISION scores can be negative. However, our analyses are focused – unless stated otherwise – on the relative score changes (i.e. increase or decrease relative to 3 months) occurring with age in a given region or cell type.

#### Bulk-seq marker genes and score calculation

Seurat’s FindAllMarkers function was run using the ‘DESeq2’ test with parameters and Bonferroni correction for multiple testing to identify region-specific marker genes (*P* value of less than 0.05; Figure S3). For each region, we constructed unsigned signatures ^25^ based on a given region’s significant marker genes. For each signature, we calculated scores across a publicly available spatial transcriptome dataset from 10X Genomics (https://www.10xgenomics.com/resources) and compared the patterns to structural annotations in Allen Mouse Brain Atlas.

#### Bulk-seq Common Aging Score (CAS) calculation and CAS velocity comparison

We ranked genes on the basis of their regulation across regions, to summarize in how many regions a given gene would be called as a DEG (i.e. reach statistical significance in at least two comparisons between samples from 3 months and any following age group). We included only the anterior hippocampus region in the selection of cross-region DEGs to prevent a potential bias towards aging effects in the hippocampus. This led to the identification of 82 genes that were marked as DEG in at least 10 out of 14 regions (15 regions minus the posterior hippocampus region). We constructed a signed gene signature^25^ based on 75 up- and 7 down-regulated DEGs. We used the signature to calculate CAS for each single-region transcriptome. To quantify a region’s score increase over time (aging velocity), we constructed a linear model with the design: score ∼ age + region + age:region (score explained by a two factor model including interaction term) using the linear model function in R. We used the lstrends function of the lsmeans package^87^ that utilizes least-square means to estimate and compare the slopes of fitted lines for each region. We subsequently used Tukey’s range test across all possible region-to-region comparisons to assess which regions exhibited statistically significant (*P* value < 0.05) slope differences. In addition, we repeated the analysis resolved for sex-specific effects across the 3, 12, 15, 18 and 21 months groups (for which we had both male and female samples). We assessed if there was a differential aging velocity between sexes across all regions (Figure 2I), for which analyzed a linear model with the design: score ∼ age + sex + age:sex + region. We further performed the same analysis iteratively for each region individually (Figure S4D) using a model with the design: score ∼ age + sex + age:sex. We corrected the resulting *P* values for each region-wise analysis using the Benjamini-Hochberg method.

#### Comparing CAS velocity with STARmap single-cell composition

We obtained meta data from Shi et al. ^41^ via the <u>Single Cell Portal</u> where the authors had quantified spatial distribution of major cell types across the entire mouse brain using their previously published, imaging-based STARmap method^88^. This atlas contains data on 422,766 single cells of 27 major cell types quantified across 73 brain structures and several sagittal sections. We aggregated the 73 reported brain structures into 10 regions that we considered meaningful equivalents of regions profiled in our Bulk-seq study (e.g. data from CA1, CA2, CA3, dentate gyrus etc. were grouped together into a ‘hippocampus’ region). We calculated for each of the 10 regions the relative abundance of each of the 27 cell types and then correlated these with the regions’ respective CAS slopes. If, for example, the relative abundance of microglia across brain regions at young age was responsible for the observed differences in CAS slopes, then we would assume a significant correlation between those two metrics across the analyzed brain regions. We did not find any significant relationship (as tested with spearman correlation and linear regression) between relative cell abundance and CAS increase over time for any of the investigated cell types (Fig. FigS6).

#### Microarray analysis of microglia

We obtained annotated and pre-processed microarray data from ^52^ (GSE62420) using limma^89^, GEOquery^90^ and GEO’s online GEO2R tool (https://www.ncbi.nlm.nih.gov/geo/info/geo2r.html). Pairwise-differential testing between age groups of the same region was performed using empirical bayes moderation as implemented in limma with multiple testing correction.

#### Organ-specific aging signature identification and velocity comparison

To explore the feasibility of detecting gene expression patterns with organ-specific regulation during aging, we re-analyzed a previously published bulk RNA-seq dataset of 17 mouse tissues profiled across ten age groups (n = 4 males; aged 1, 3, 6, 9, 12, 15, 18, 21, 24 and 27 months; n = 2 females; aged 1, 3, 6, 9, 12, 15, 18 and 21 months)^8^. The dataset comprised the following organs: bone, brain, brown adipose tissue (BAT), gonadal adipose tissue (GAT), heart, kidney, limb muscle (muscle), liver, lung, bone marrow (marrow), mesenteric adipose tissue (MAT), pancreas, skin, small intestine (intestine), spleen, subcutaneous adipose tissue (SCAT), and white blood cells (WBC). We obtained pre-processed data as described in the original study and performed differential expression analysis accordingly^8^. We identified age-related DEGs in the same manner as described for the bulk-seq data: we used the raw count matrix as recommended for the DEseq2 standard analysis pipeline. Factors and dispersion estimates were calculated for each tissue separately. We conducted differential expression analysis comparing samples from 3-months-old mice to each consecutive time point, using age and sex as covariates. *P* values were adjusted for multiple testing, and genes with an adjusted *P* value of less than 0.05 were determined to be statistically significant. Finally, we required a gene to reach statistical significance (after multiple testing correction) in at least 2 pairwise comparisons (e.g. 3-months-old vs 12 months-old and 3-months-old vs 21 months-old) to be called a differentially expressed gene (DEG). We analyzed age groups that would be comparable to the age groups profiled in our study (3, 12, 15, 18, 21, 24 and 27 months). We ranked genes on the basis of their regulation across organs, to summarize in how many organs a given gene would be called as a DEG (i.e. reach statistical significance in at least two comparisons between samples from 3-months-old mice and any following age group). DEGs that were only detected in a single organ were assembled into signed, organ-specific aging signatures using VISION ^25^, comparable to the CAS. For organs that exhibited fewer than 25 unique DEGs we did not construct a signature. For each organ-specific signature, we performed the following analysis: We first tested for each organ separately, if the respective signature would show a significant correlation with age using linear models with the design: score ∼ age. Organs that showed no significant (*P* val < 0.05, *t*-test) association with the age were excluded. Next, we constructed a linear model with the design: score ∼ age + organ + age:organ (organ-specific score explained by a two factor model including interaction term) using the linear model function in R. We used the lstrends function of the lsmeans package^87^ that utilizes least-square means to estimate and compare the slopes of fitted lines for each organ. We subsequently used Tukey’s range test across all possible organ-to-organ comparisons to assess which organs exhibited statistically significant (*P* value < 0.05) slope differences. Notably, we asked if the organ where the signature was identified (the ‘reference’ organ) would show a significantly higher slope compared to all other organs. The summarized results are displayed in the heatmap in Figure S9.

#### Bulk-seq region-specific aging signature identification and velocity comparison

We ranked genes on the basis of their regulation across regions, to summarize in how many regions a given gene would be called as a DEG (i.e. reach statistical significance in at least two comparisons between samples from 3-months-old mice and any following age group). DEGs that were only detected in a single region were assembled into signed, region-specific aging signatures using VISION ^25^, comparable to the CAS. We excluded the posterior hippocampus region in the selection of region-specific DEGs. Further, there were less than 20 unique DEGs found for the entorhinal cortex, which we considered too small to construct a signature. For each region-specific signature, we performed the following analysis: We first tested for each region separately, if the respective signature would show a significant correlation with age using linear models with the design: score ∼ age. Regions that showed no significant (*P* val < 0.05, *t*-test) association with age were excluded. Next, we constructed a linear model with the design: score ∼ age + region + age:region (region-specific score explained by a two factor model including interaction term) using the linear model function in R. We used the lstrends function of the lsmeans package^87^ that utilizes least-square means to estimate and compare the slopes of fitted lines for each region. We subsequently used Tukey’s range test across all possible region-to-region comparisons to assess which regions exhibited statistically significant (*P* value < 0.05) slope differences. Notably, we asked if the region where the signature was identified (the ‘reference region’) would show a significantly higher slope compared to all other regions. The summarized results are displayed in the heatmap in Figure S10.

#### 10X Visium mapping, embedding, clustering and region identification

Space Ranger analysis pipelines were utilized to align image and FASTQ files, detect tissue and fiducial frames, count barcodes/UMIs. Throughout the manuscript we refer to the barcoded areas from a given dataset as ‘spots’. Spots with less than 5 UMIs were removed as well as all spots at the outline of the tissue as these can be affected by RNA diffusion. We integrated all six sample-wise datasets (two from 6 months, two from 18 months and two from 21 months), using Seurat’s built-in SCTransform and integration workflow^81^, with 2000 genes set as integration features. Integrated datasets were then used as input for spot embedding and clustering. A shared-nearest-neighbors graph was constructed using the first 30 PC dimensions before clustering spots using Seurat’s built-in FindClusters function with a resolution of 0.8 and default parameters. Umaps and tSNEs were calculated using Seurat’s built-in functions, based on the first 30 PC dimensions.

Count data was subsequently normalized and scaled using SCTransform across all spots to allow for visualization of expression values and differential gene expression analysis.

We chose a data-driven approach to group spots and map them to anatomical structures of the brain (Figure S5A): Transcriptional clustering yielded 29 clusters and we used Seurat’s FindAllMarkers function (parameters: min.pct=0.1, thresh.use=0.1 assay=’SCT’, only.pos=TRUE) to identify cluster markers. We compared the expression of marker genes to in-situ hybridization (ISH) data from the Allen Mouse Brain Atlas^26^ and visual landmarks from the H&E microscopy images (e.g. *cornu Ammonis* and dentate gyrus of the hippocampus; cell-sparse structure of the white matter fiber tracts). To enable comparisons with the regions isolated for Bulk-seq, we additionally grouped annotated clusters into meaningful region-level sets, guided by their anatomical location and the hierarchical ordering of structures in the Allen Mouse Brain Atlas. Ontology and nomenclature of clusters is indicated in Fig S5B.

#### 10X Visium differential expression analysis and comparison with Bulk-seq data

Given comparable representation of clusters across all samples and age groups, we considered differential expression analysis across age groups feasible. We analyzed differential expression in the white matter and cortex cluster as we considered them comparable to the corpus callosum (high CAS velocity) and motor cortex (low CAS velocity) region from the bulk-seq dataset. Differential gene expression of genes comparing 10X Visium data from 6 months to 21 months was done using the ‘DESeq2’ algorithm implemented in Seurat on Spatial count data. Seurat natural log(fold change) > 0.2 (absolute value), adjusted *P* value (Benjamini-Hochberg correction) < 0.05, and expression in greater than 10% of spots in both comparison groups were required to consider a gene differentially expressed. To test for a potential association between gene-expression changes measured in 10X Visium and bulk-seq data, we considered only genes that changed significantly in both datasets. For both regions, we confirmed significant overlap between the DEGs found in Bulk-seq and 10X Visium dataset (Fisher’s exact test, *P* Value < 0.05). Next, we plotted log2 fold expression changes during aging as measured via Visium versus log2 fold expression changes on the bulk-seq level. The distribution of genes among the four resulting quadrants was tested for directionality using Fisher’s exact test.

#### Visium CAS calculation

Calculation of CAS and CAS velocities for Visium data was carried out in a similar manner as described above for bulk-seq data: CAS for each Visium spot were calculated and score increase over time was calculated for region clusters with equivalent Bulk-seq regions: White matter (compared to corpus callosum), cortex (compared to motor cortex), striatum (compared to caudate putamen), hippocampus, hypothalamus, choroid plexus and thalamus. To quantify a region’s score increase over time (aging velocity), we constructed a linear model with the design: score ∼ age + region + age:region and carried out slope estimation and differential analysis as described above. We acknowledge that this analysis does not account for biological replicates but treats each spot belonging to the same region as replicate. We therefore visualized CAS in Visium for each replicate, to demonstrate that age-related changes in CAS supersede the intra-replicate CAS differences.

#### Nuc-seq mapping, embedding, clustering, sample demultiplexing and cell type identification

Cell Ranger (v.6.1.2) analysis pipelines were utilized to align reads to mm10 reference genome and count barcodes/UMIs. To account for unspliced nuclear transcripts, reads mapping to pre-mRNA were counted. Throughout the manuscript we use nuclei and ‘cells’ synonymously. Outliers with a high ratio of mitochondrial (more than 5%, fewer than 400 features) relative to endogenous RNAs and homotypic doublets (more than 6,000 features in hippocampus; more than 7,000 features in caudate putamen) were removed in Seurat^81^. We integrated all sample-wise datasets, using Seurat’s built-in SCTransform and integration workflow^81^, with 500 genes set as integration features. Integrated datasets were then used as input for cell embedding and clustering. A shared-nearest-neighbors graph was constructed using the first 12 PC dimensions before clustering spots using Seurat’s built-in FindClusters function with a resolution of 0.4 and default parameters. Umaps and tSNEs were calculated using Seurat’s built-in functions, based on the first 12 PC dimensions. A given nuc-seq sample represented nuclei from a male and female animal (of the same age) that were pooled in equal numbers during the nuclei isolation steps. To demultiplex a sample by sex, we calculated the ratio of counts belonging to female-(*Xist*, *Tsix*) and male-specific (*Ddx3y*, *Eif2s3y*, *Uty*, *Kdm5d*) genes, and identified nuclei with a log2 cutoff of 1 and -1 as female- and male-derived nuclei, respectively. Ambiguous nuclei, which had reads from both female and male nuclei, were removed from the analysis. Count data was subsequently normalized and scaled to allow for visualization of expression values and differential gene expression analysis. Seurat’s FindAllMarkers function (parameters: min.pct=0.15, thresh.use=0.15 assay=’SCT’) was run to identify cluster markers. Clusters were annotated based on marker genes. Finally, nuclei were manually inspected using known cell type-specific marker genes and nuclei expressing more than one cell type-specific marker were defined as doublets and removed^91^.

#### Publicly available scRNA-seq data embedding

We re-analyzed two previously published single-cell RNA-seq datasets: (1) Droplet-based scRNA-seq of freshly dissected SVZ at young and old age (n = 3 males per age; aged 3 and 28 months; all C57BL/6JN strain)^27^; (2) Smart-seq2-based ^49^ scRNA-seq of freshly isolated cells from the myeloid and non-myeloid fraction of the striatum, cerebellum, hippocampus and cortex at young and old age (n = 4 males per age; aged 3 and 24 months; all C57BL/6JN strain) ^19^. For the SVZ data, we obtained and analyzed pre-processed count matrices. For visualization purposes, we integrated all sample-wise datasets, using Seurat’s built-in SCTransform and integration workflow^81^, with 2000 genes set as integration features. Integrated datasets were then used as input for cell embedding. Umaps and tSNEs were calculated using Seurat’s built-in functions, based on the first 12 PC dimensions. Cell annotations were transferred from the original study. Count data was subsequently normalized and scaled to allow for visualization of expression values.

For the second dataset, we obtained and analyzed pre-processed count matrices. We followed previous analyses on the same dataset^73^. For visualization purposes, we integrated all sample-wise datasets, using Seurat’s built-in SCTransform workflow^81^. Integrated datasets were then used as input for cell embedding. Umaps and tSNEs were calculated using Seurat’s built-in functions, based on the first 12 PC dimensions. Cell annotations were transferred from the original study. Count data was subsequently normalized and scaled to allow for visualization of expression values.

#### Differential expression in scRNA- and Nuc-seq data

Differential gene expression of genes comparing young and old samples was done using the MAST^92^ algorithm, which implements a two-part hurdle model. Seurat natural log(fold change) > 0.2 (absolute value), adjusted *P* value (Benjamini-Hochberg correction) < 0.05, and expression in greater than 10% of cells in both comparison young and old samples.

#### Signature calculations in scRNA- and Nuc-seq data

Calculation of CAS and CAS velocities for scRNA- and Nuc-seq data was carried out in a similar manner as described above for 10X Visium data: CAS for each cell were calculated and CAS increase over time was calculated for cell types. To quantify a cell type’s score increase over time (aging velocity), we constructed a linear model with the design: score ∼ age + cell type + age:cell type and carried out slope estimation and differential analysis as described above. We acknowledge that this analysis does not account for biological replicates but treats each cell belonging to the same cell type as replicate. To account for this, we further calculated the cell type-median CAS for each biological replicate and tested for differential CAS regulation with two-tailed t-test on per-replicate median of CAS. For the comparison of CAS increase with age across microglia from different brain regions (Figure 4A-C), we used two-sided Wilcoxon rank-sum tests to test for CAS differences between microglia from the same age group.

For region-specific signatures, we performed a per-cell type slope quantifications as detailed for the CAS and clustered the resulting slope estimates using hierarchical clustering (Figure 5H,K).

#### Single-nuclei dispersion score

We employed a previously published strategy to quantify if a given DEG detected in bulk data would be expressed in a specific cell type^8^. For each gene in each brain region, we selected cells expressing the gene (log-CPM expression > 0). Next, we assigned to the cells the log-CPM expression values of the gene as weights. Based on these, we calculated the weighted center of the cells in the single-cell landscape defined by the UMAP embeddings. We defined the ‘single-cell dispersion’ of the gene as the weighted mean distance of the cells from their weighted center. Finally, we introduced region specific factors to account for differences between brain region specific embeddings. Per region, we set pseudo log-CPM count 1 to all cells and calculated the dispersion of them. We normalized the dispersion scores by these region-specific factors.

## Acknowledgments

We thank the members of the Wyss-Coray laboratory for feedback and support, and H. Zhang, D. Berdnik, K. Dickey and D. Channappa for laboratory management. We further thank the Neuroscience Microscopy Service center and director G. Wang for microscopy training and assistance in acquiring images for the Visium experiments. We wish to thank S. Buschbaum, L. Drews, J. Fröhlich, O. Hendrich, R. Hoppe and A. Pahl for preparing the murine tissue material from the C3B6F1 cohort. We thank A. Bastian for advice in establishing the Visium experiments. We thank the Gökce lab for providing pre-processed SS2 glia data. The research leading to these results has received funding from the European Research Council under the European Union’s Seventh Framework Programme (FP7/2007-2013) / ERC grant agreement n° 268739 (L.P.), the Max Planck Society (L.P.), the Schaller-Nikolich Foundation (A.Ke.), the Wu Tsai Neurosciences Institute and Fondation Bertarelli (T.W.-C.), the Simons Foundation (T.W.-C.), the Cure Alzheimer’s Fund (T.W.-C.), a National Institute of Aging grant R01-AG072255 (T.W.-C.), the Milky Way Research Foundation (T.W.-C.), the American Heart Association-Allen Initiative in Brain Health and Cognitive Impairment (T.W.-C.), The Phil and Penny Knight Initiative for Brain Resilience (T.W.-C.), Michael J. Fox Foundation for Parkinson’s Research grants 125491594 (A.Ke and F.K.) and MJFF-021418 (T.W.-C., A.Ke. and F.K.).

## Author contributions

O.H. and T.W.-C. conceptualized the study. O.H. designed and led experiments, conducted bioinformatic analyses and prepared the manuscript. A.G.F. and O.H. selected regions and devised the brain dissociation method. J.H. and J.R.H aided in region selection and method design. O.H., I.H.G., C.M., A.G.F., M.A., B.K. and A.Ka. conducted the tissue collection. A.G.F. dissociated and organized brain regions, and extracted RNA. N.L. and O.H. established the bulk-seq workflow. A.G.F., M.A. and O.H. processed libraries for bulk-seq. B.K. and O.H. conducted the 10X Visium experiments. I.H.G., C.M. and A.Ka. aided with the H&E workflow. M.A., C.M. and A.G.F. performed the nuc-seq experiments. S.B. and L.P. provided hippocampus tissue from C3B6F1 mice. P.M.L., B.L., R.P., F.K. and A.Ke. assisted with analysis and bioinformatics procedures. B.K. developed the searchable web interface (Shiny app). O.H. and T.W.-C. edited the manuscript with input from all authors. All authors read and approved the final manuscript.

## Declaration of interests

The authors declare no competing interests.

(F) Smoothed line plot displaying the number of DEGs for pairwise comparisons, referenced to data at 3 months. Positive (negative) values represent upregulated (downregulated) genes, gray lines represent non-labelled regions. DEGs that reached significance in ≥ 2 pairwise comparisons were included. (G) Heat map of the data in (F). (H) Number of genes that significantly correlate with age (Spearman’s rho ≥ 0.5), colored by up- and down-regulation. (I) Networks of the most highly connected genes (‘eigengenes’) of three exemplary modules with significant age-association identified in corpus callosum, motor cortex and thalamus. Networks display connections of the corresponding topological overlap above a threshold of 0.08. (J) Chord diagram representation of genes shared in age-associated modules across regions. Modules with significant enrichment of cell type markers are displayed. Modules and associated genes are listed in Table S4.

## Supplementary Figures

**Figure S1.**
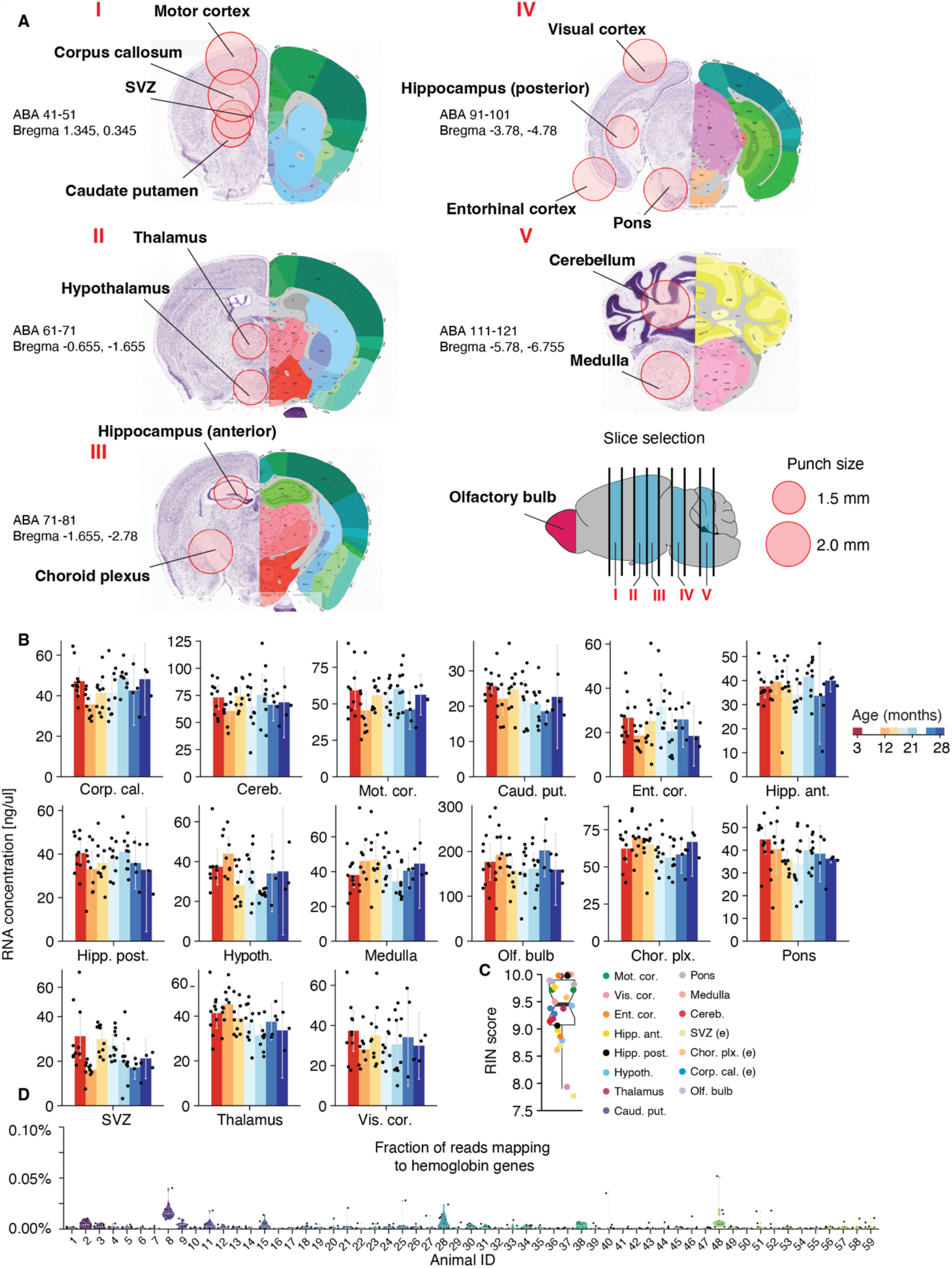
Capturing transcriptional heterogeneity of the brain with high RNA integrity (A) Location of tissue punches overlaid on the Allen Institute Mouse Coronal Reference Atlas. Equivalent regions were taken from both the left and right hemisphere. Depth of each brain slice is indicated as: ABA, Allen brain atlas coronal layer number range, and Bregma, stereotaxic coordinates of mm. from bregma. The following regions required overlapping punches and were thus sequentially collected: (1) motor cortex, (2) caudate putamen, (3) subventricular zone, (4) corpus callosum. (B) RNA concentration measured with Quant-iT broad range RNA Assay Kit in all regions, colored by age. Data are mean ± s.e.m. (C) RNA integrity number, determined by Bioanalyzer eukaryote total RNA, for randomly-selected samples, colored by region. Each region is represented by at least 3 samples. RINs above 7 indicate good quality RNA. (D) Violin plot of fraction of reads mapping to hemoglobin genes for each sample, grouped by animal.

**Figure S2.**
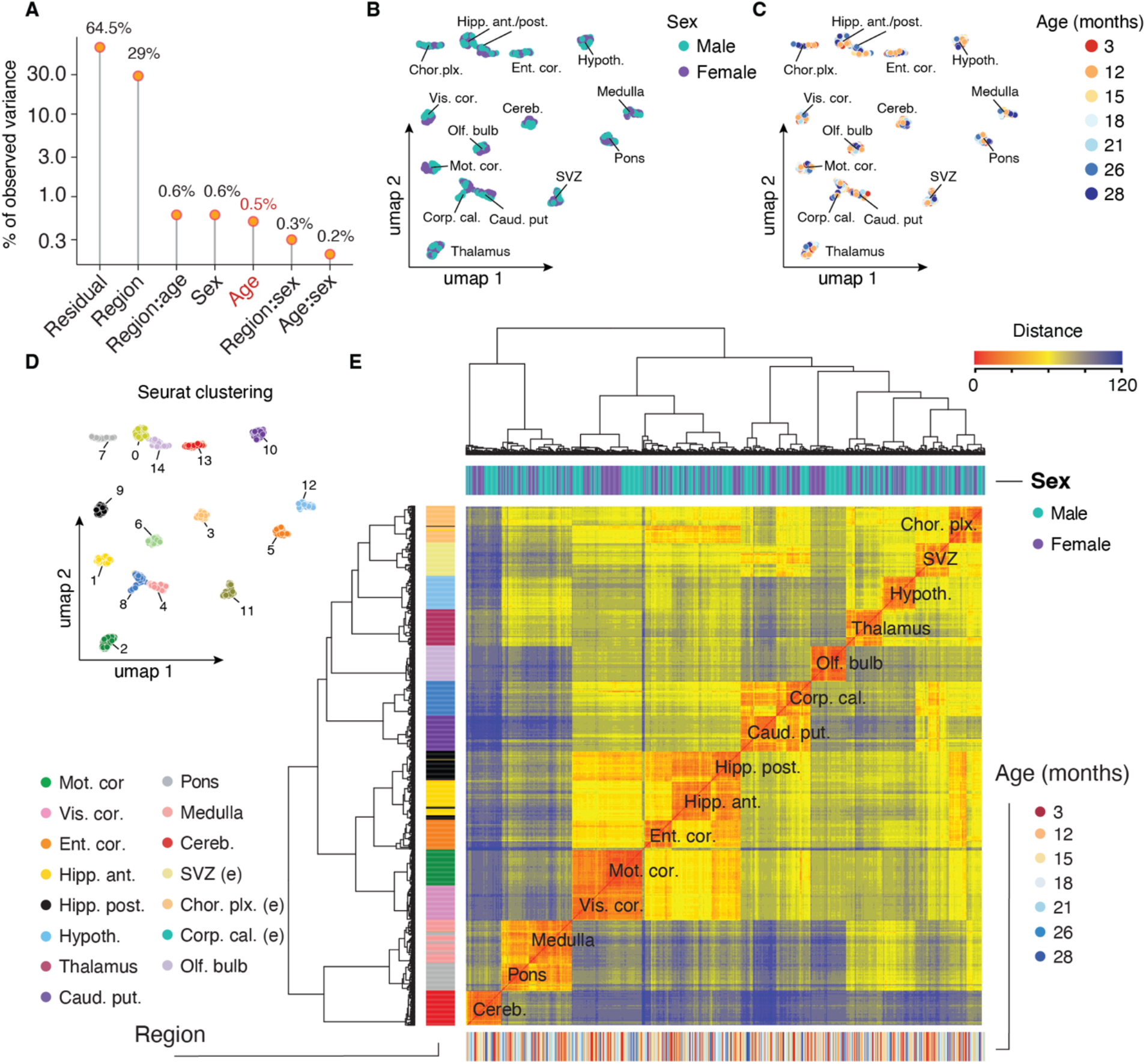
Gene expression variance analysis across regions (A) Visualization of the principal variance component analysis, displaying the gene expression variance explained by residuals (that is, biological and technical noise) or experimental factors such as region, age, sex and respective interactions. Highlighted is variance explained by age. (B,C,D) UMAP representation of bulk RNA-seq samples (n = 847 total samples) based on the first 40 principal components colored according to (B) sex, (C) age, and (D) Seurat clustering with default settings. (E) Hierarchical clustering of all samples using Ward’s algorithm. Samples are annotated by region, sex and age.

**Figure S3.**
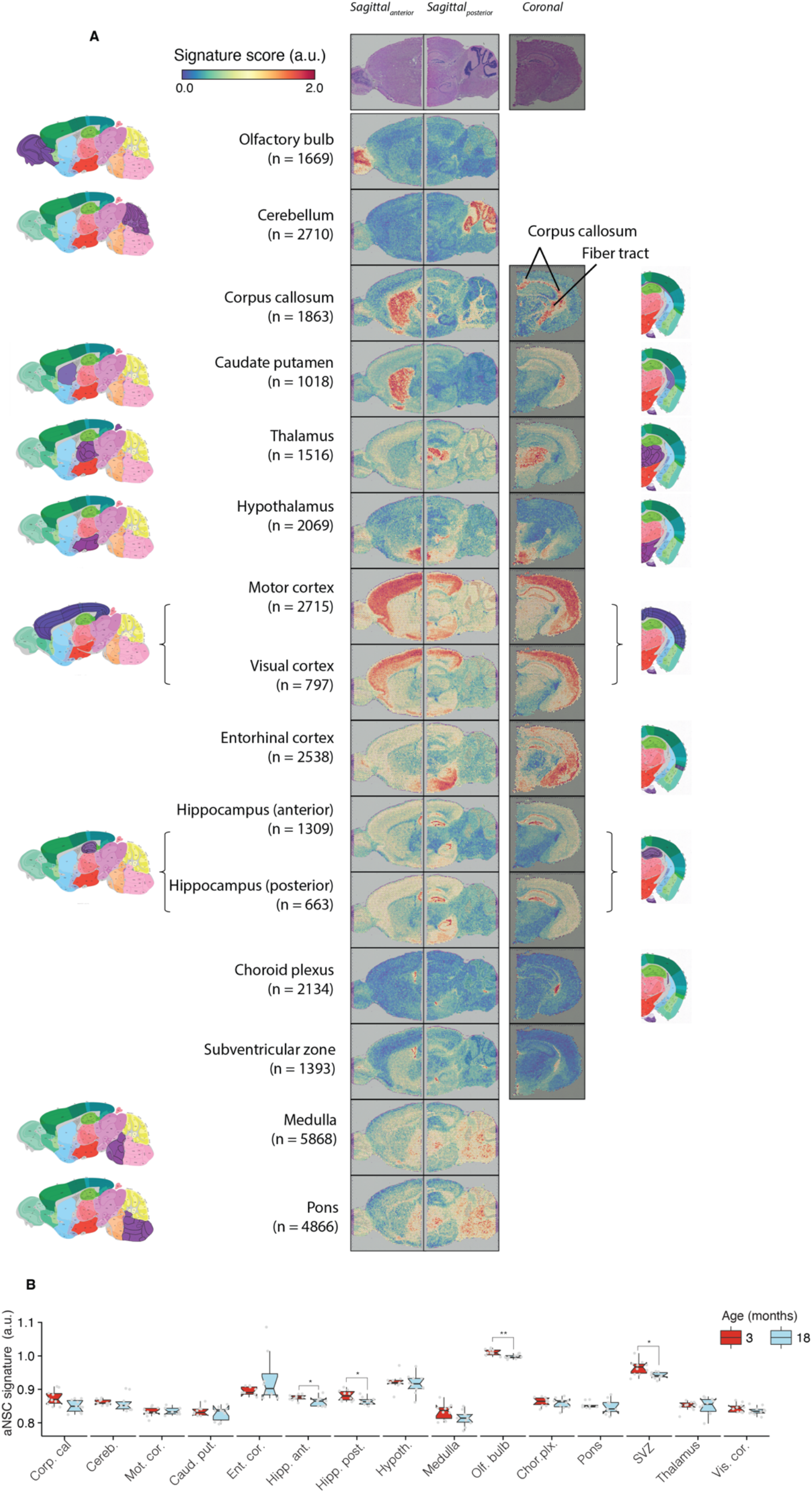
Collected brain regions reflect transcriptomes of anatomically distinct structures (A) Spatial expression of gene signatures derived from marker genes enriched in regions from Fig S1. For each region, marker gene analysis was performed and enriched genes were grouped into a marker signature. Each signature was quantified in publicly available Visium spatial transcriptomics data from a young mouse brain. Dataset contained anterior and posterior sagittal sections, as well as a coronal section. Number of marker genes that make up the respective signature is indicated in parentheses and marker genes are listed in Table S1. Quantification of gene signatures in spatial transcriptome yields scores for each spatial spot. Scores for each signature were compared to the Allen Institute Mouse Sagittal and Coronal Reference Atlases where possible. Positions of regions that we aimed to capture are highlighted in purple. (B) Expression of activated neural stem cell (aNSC)-derived marker gene signature in young (3 months) and aged (18 months) samples. Signature exhibits the highest baseline expression in the olfactory bulb and subventricular zone (SVZ) derived samples. A significant drop in the signature score is notable in olfactory bulb, SVZ and both anterior and posterior hippocampus regions. *P* values calculated with two-tailed t-test on per-replicate median of signature score, adjusted for multiple testing. *** p < 0.001, ** p < 0.01, * p < 0.05

**Figure S4.**
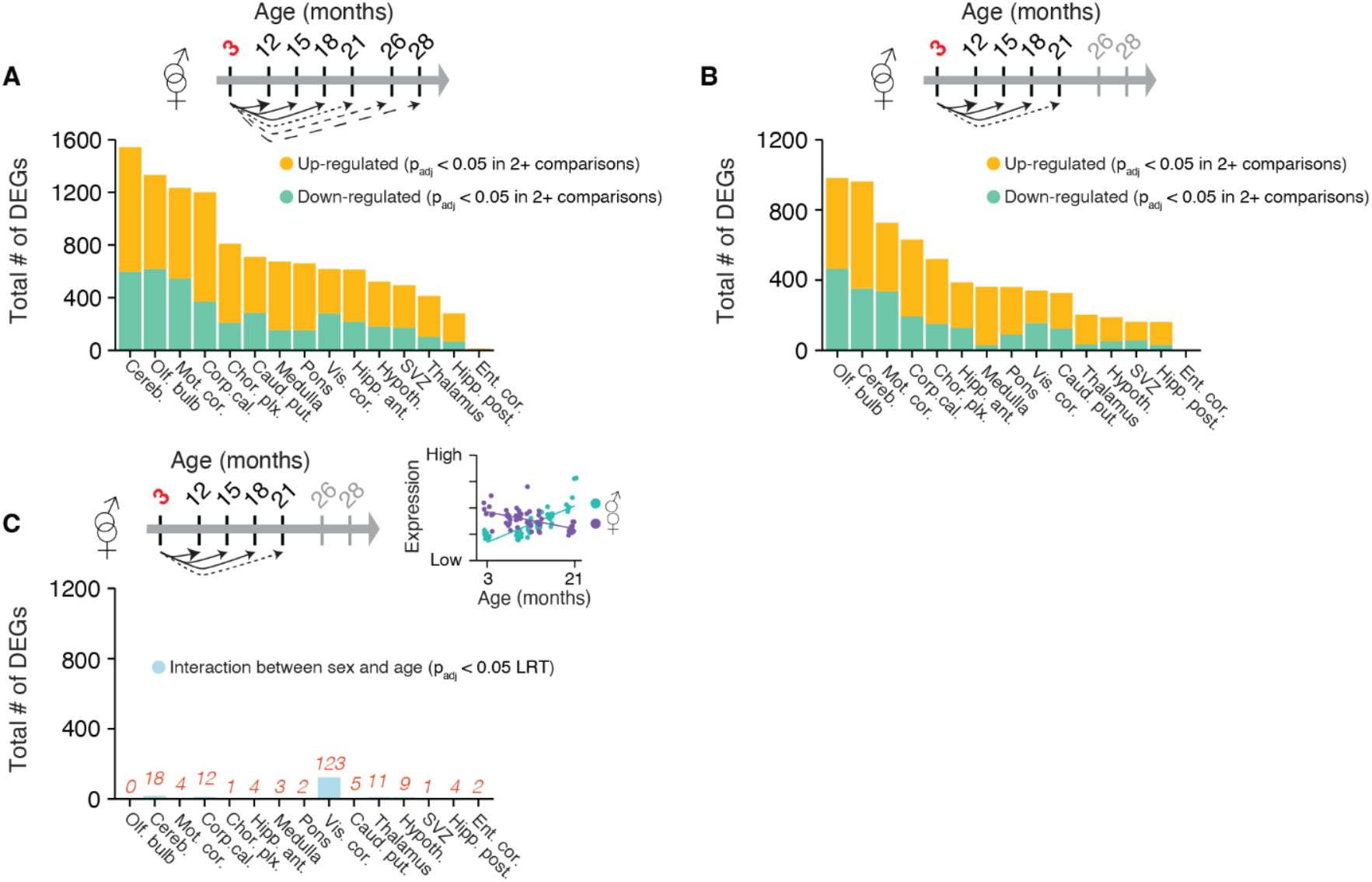
Validation of differential gene expression analysis (A) Number of differentially expressed genes in 2+ age comparisons (12 mo., 15 mo., 18 mo., 21 mo., 26 mo. and 28 mo.) relative to 3 mo. Split by region, colored by up- and down-regulation. padj ≥ 0.5 in 2+ comparisons. (B) Number of differentially expressed genes in 2+ age comparisons (12 mo., 15 mo., 18 mo., 21 mo.) relative to 3 mo. Split by region, colored by up- and down-regulation. padj ≥ 0.5 in 2+ comparisons. (C) Total number of genes exhibiting a significant interaction between sex and age split by region. padj < 0.05 LRT^33^.

**Figure S5.**
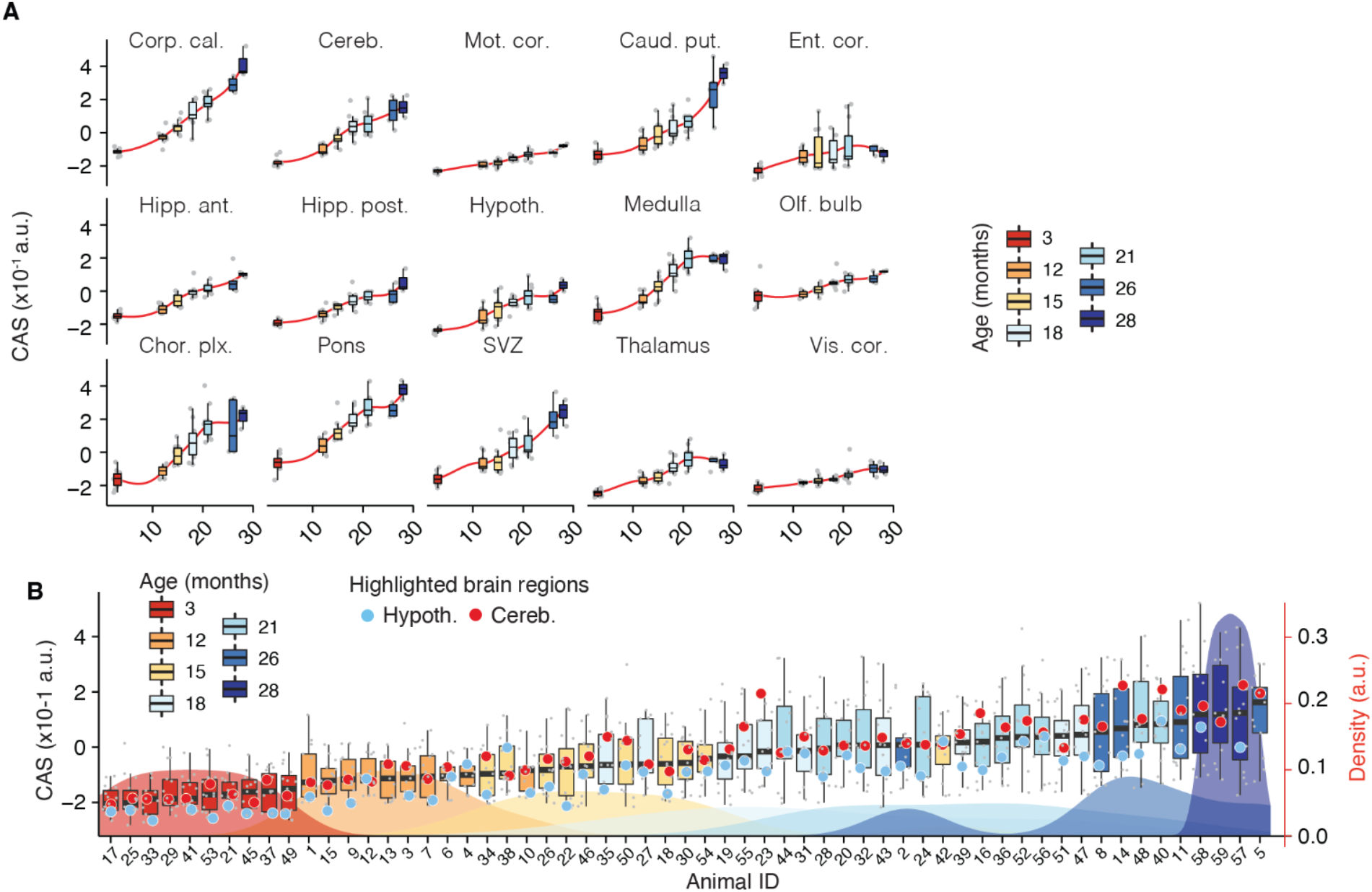
A common RNA aging signature quantifies the region-specific pace and magnitude of transcriptional shifts in the brain (A) CAS trajectories of all regions, colored by age. Red lines indicate averaged-smoothed values. (B) CAS for each sample grouped by animal. Each boxplot represents distribution of CAS across the 15 regions as measured by bulk RNA-seq. Animals are ranked by median score and colored by chronological age. Distribution of chronological age groups is indicated in the background.

**Figure S6.**
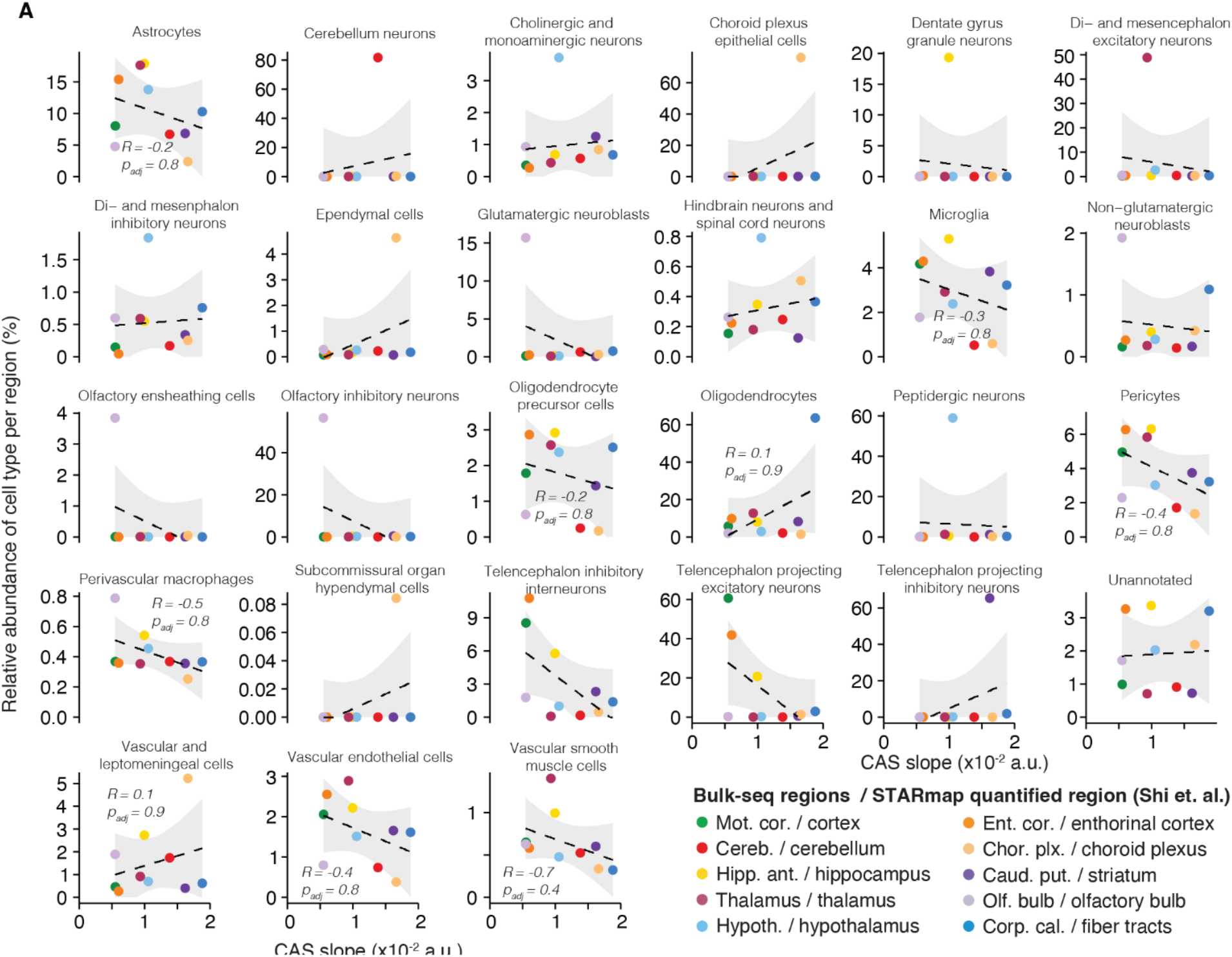
CAS increase is not associated with regions’ cell type composition (A) The correlation of the relative abundance of 27 cell types with the regions’ respective CAS slopes. Each plot shows the relative abundance of a different cell type in each of the regions, (determined from the equivalent region from ^41^, with CA1, CA2, CA3, dentate gyrus etc. grouped together into a ‘hippocampus’ region) plotted against the CAS slope (determined from our Bulk dataset). Each point represents a different region. Significance of this relationship tested through spearman correlation and linear regression, with no significant trends found.

**Figure S7.**
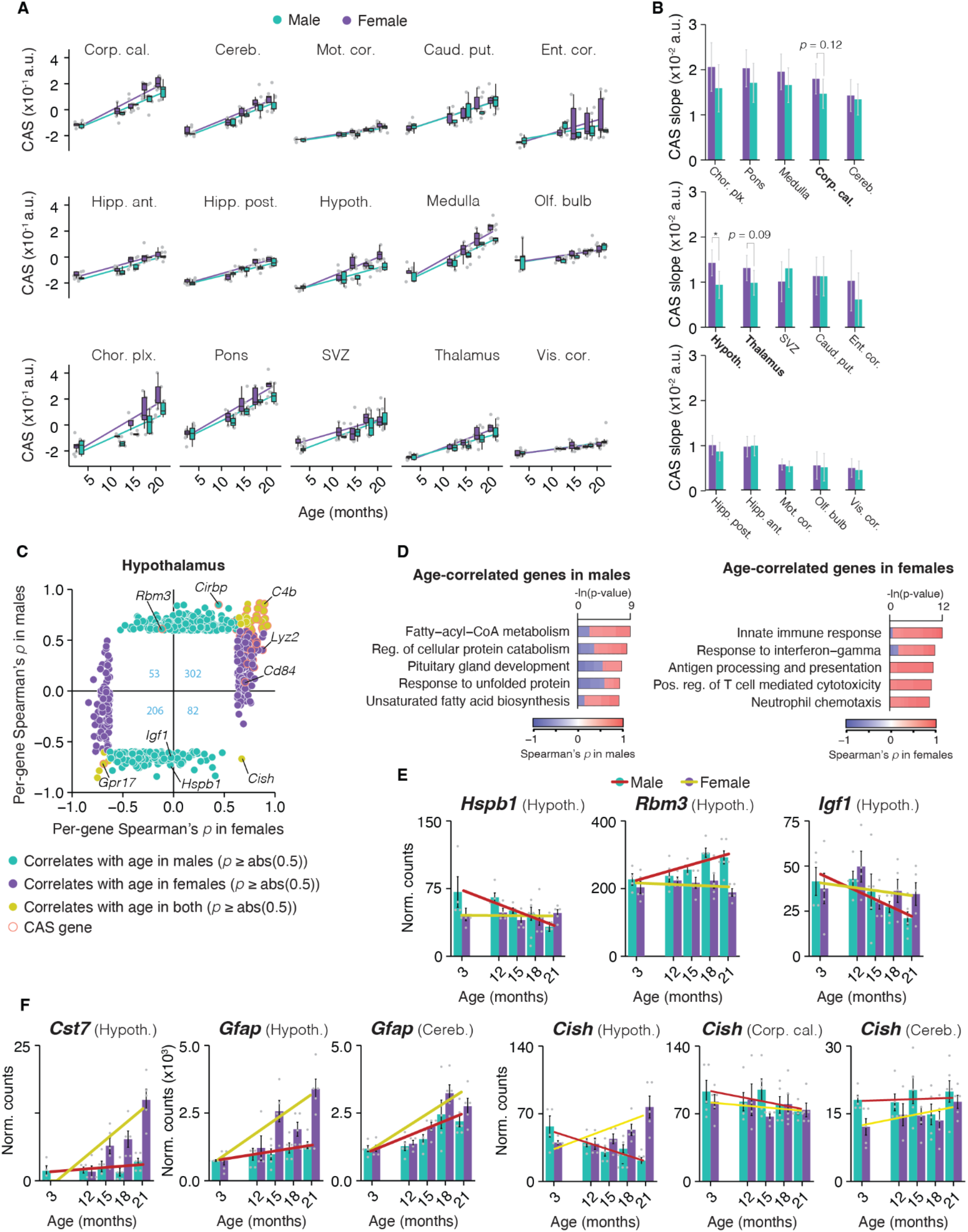
Hypothalamus aging is accelerated in females (A) CAS trajectories of all regions for the interval of 3 to 21 months, colored by sex. (B) Sex-specific slope of linear regressions in (A), colored by sex. Data are mean ± 95% confidence intervals. Pair-wise, two-sided Tukey’s HSD test. For each region, we calculated a separate model limited to data from the respective tissue only. *** p < 0.001, ** p < 0.01, * p < 0.05. The highest (least significant) Pval is indicated. (C) Scatterplot representation of genes that significantly correlate with age (Spearman’s rho ≥ 0.5) in males, females or both. The number of overlapping DEGs in each quadrant is indicated in blue. (D) Representative GO analysis of genes with significant age-correlation in males or females. Lengths of bars represent negative ln-transformed Padj using two-sided Fisher’s exact test. Colors indicate gene-wise age-correlation coefficient. (E) Bulk expression of genes with age-correlation in males, split by male and female samples. Lines indicate linear fit of gene expression resolved by sex. Data are mean ± s.e.m. (F) same as (E) for genes with significant age-correlation in females. Expression in hypothalamus, cerebellum and corpus callosum are shown to demonstrate the specificity of sexual dimorphisms in the hypothalamus.

**Figure S8.**
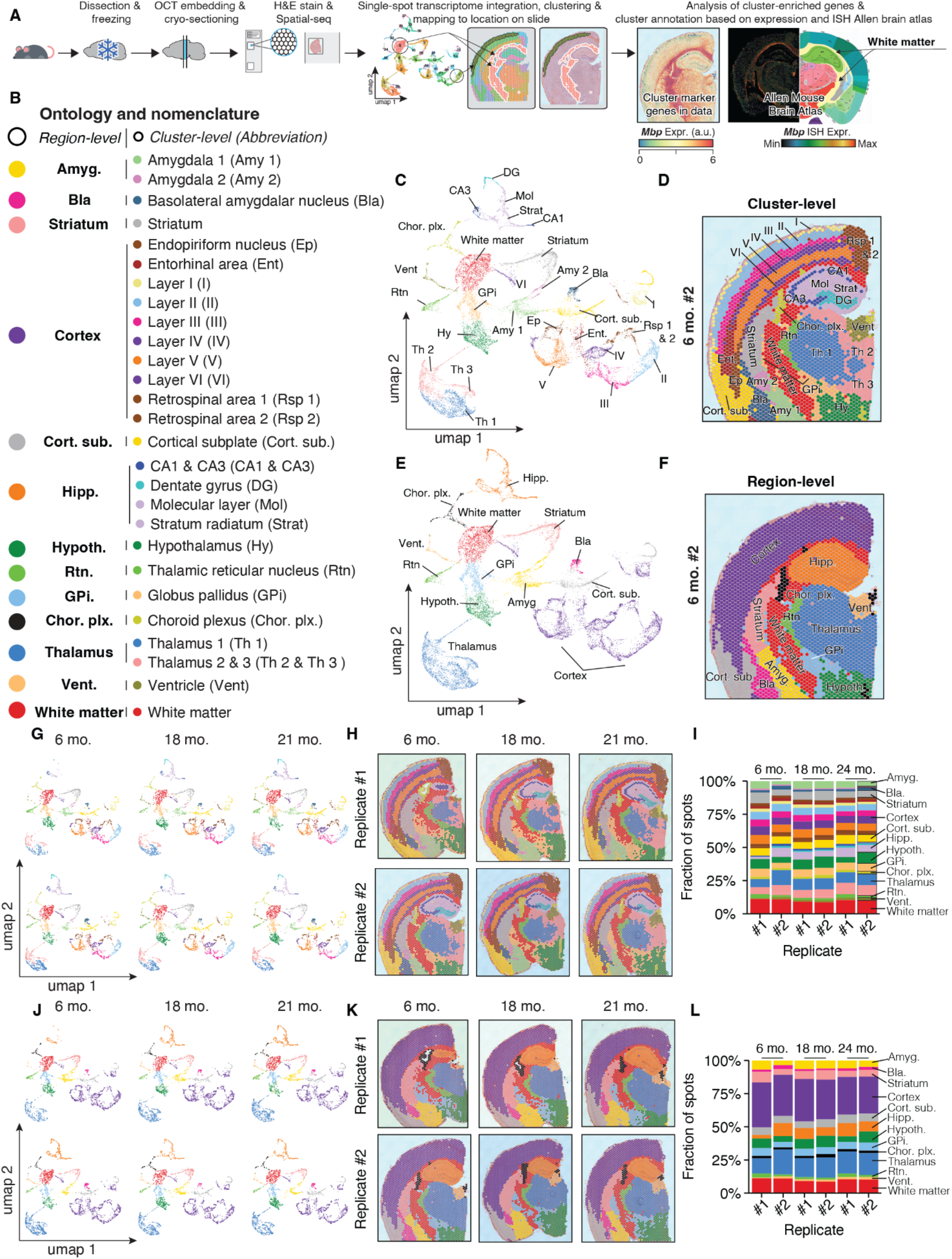
Robust capture of spatial transcriptomes across age (A) 10X Visium processing and analysis overview. Whole brains were frozen prior to OCT embedding and cryo-sectioning. Coronal sections were placed on a 10X Visium Spatial Gene Expression slide, followed by H&E staining and spatial reverse-transcription reaction. Single-spot transcriptomes were integrated, clustered with default settings and visualized as UMAP. Clustered spatial spot transcriptomes were mapped to their original location. To annotate the clusters, their marker genes (Table S6) were visualized, compared to the Allen Brain Atlas^26^. (B) Complete data description and abbreviations of ontology and nomenclature for spatial transcriptome data. Regional-level annotated manually, and cluster-level determined by Seurat clustering. (C,D) Representative spatial transcriptome data (6 months replicate #2), colored by cluster-level annotation and represented as (C) UMAP and (D) spatial transcriptome. (E,F) Representative spatial transcriptome data (6 months replicate #2), colored by region-level annotation and represented as (E) UMAP and (F) spatial transcriptome. (G,H,I) Cluster-level annotation across replicates and datasets represented as (G) UMAP and (H) spatial transcriptome. (I) Fraction of spots corresponding to each cluster. (J,K,L) Region-level annotation across replicates and datasets represented as (J) UMAP and (K) spatial transcriptome. (L) Fraction of spots corresponding to each region.

**Figure S9.**
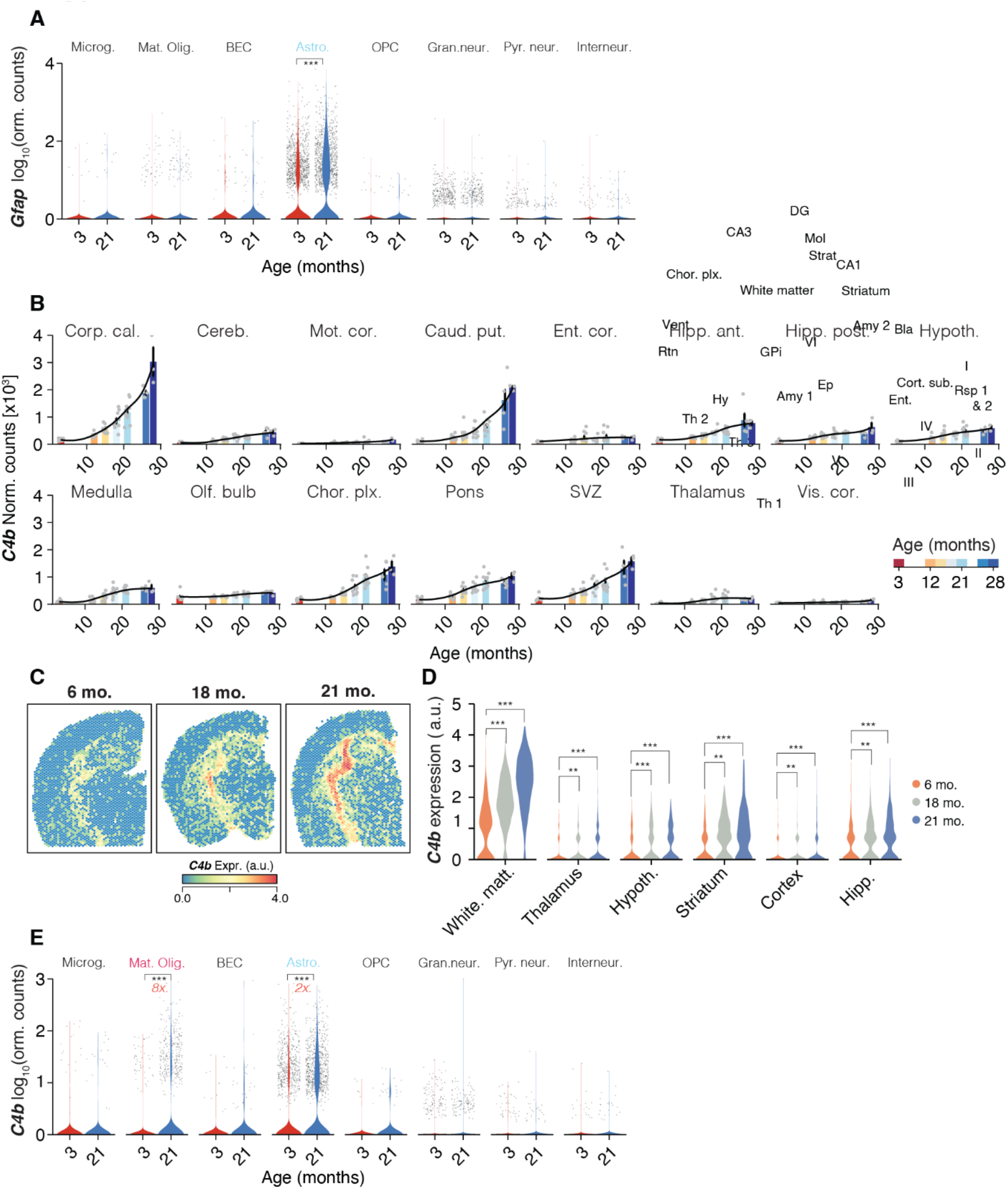
Cell type-specific quantification of *Gfap* and *C4b* (A) Violin plot of *Gfap* expression across hippocampal cell types. Points indicate nuclei-wise expression levels, and the violin indicates average distribution of expression split by age. (B) *C4b* expression across all bulk regions, colored by age. Black lines indicate averaged-smoothed gene expression. Data are mean ± s.e.m. (C) *C4b* expression in spatial transcriptome across age. (D) Violin plot of *C4b* expression in exemplary region-level clusters of spatial transcriptome data, according to Fig S5. (E) Violin plot of *C4b* expression across hippocampal cell types. Points indicate nuclei-wise expression levels, and the violin indicates average distribution of expression split by age. (MAST, Benjamini–Hochberg correction; false discovery rate (FDR) < 0.05 and logFC > 0.2 to be significant). *** p < 0.001, ** p < 0.01, * p < 0.05.

**Figure S10.**
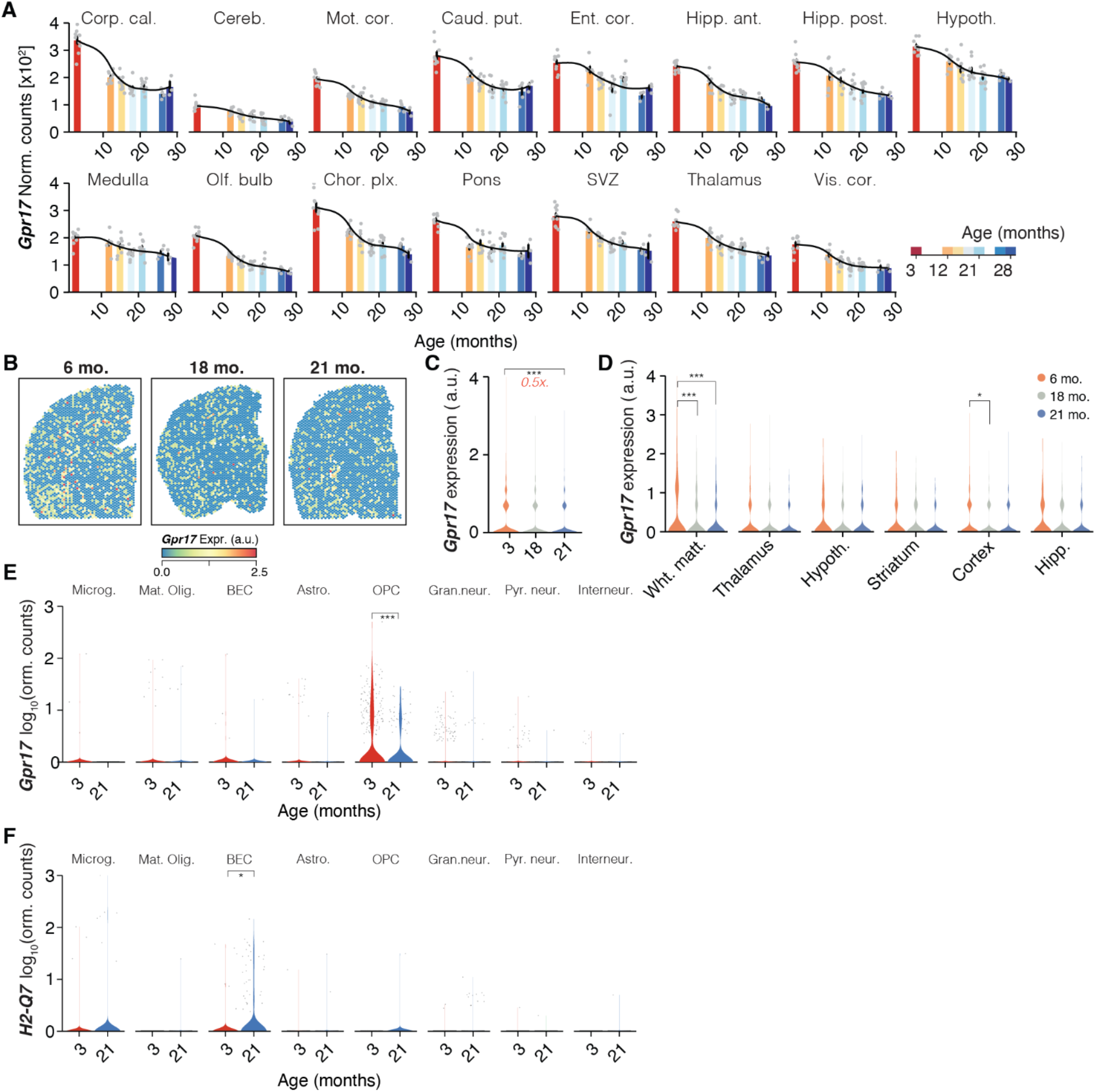
Cell type-specific quantification of *Gpr17* and *H2-Q7* (A) *Gpr17* expression across all bulk regions, colored by age. Black lines indicate averaged-smoothed gene expression. Data are mean ± s.e.m. (B) *Gpr17* expression in spatial transcriptome across age. (C) Violin plot of *Gpr17* expression full spatial transcriptome data across ages. (D) Violin plot of *Gpr17* expression in exemplary region-level clusters of spatial transcriptome data, according to Fig S5. (E,F) Violin plot of (E) *Gpr17* and (F) *H2-Q7* expression across hippocampal cell types. Points indicate nuclei-wise expression levels, and the violin indicates average distribution of expression split by age. (MAST, Benjamini–Hochberg correction; false discovery rate (FDR) < 0.05 and logFC > 0.2 to be significant). *** p < 0.001, ** p < 0.01, * p < 0.05.

**Figure S11.**
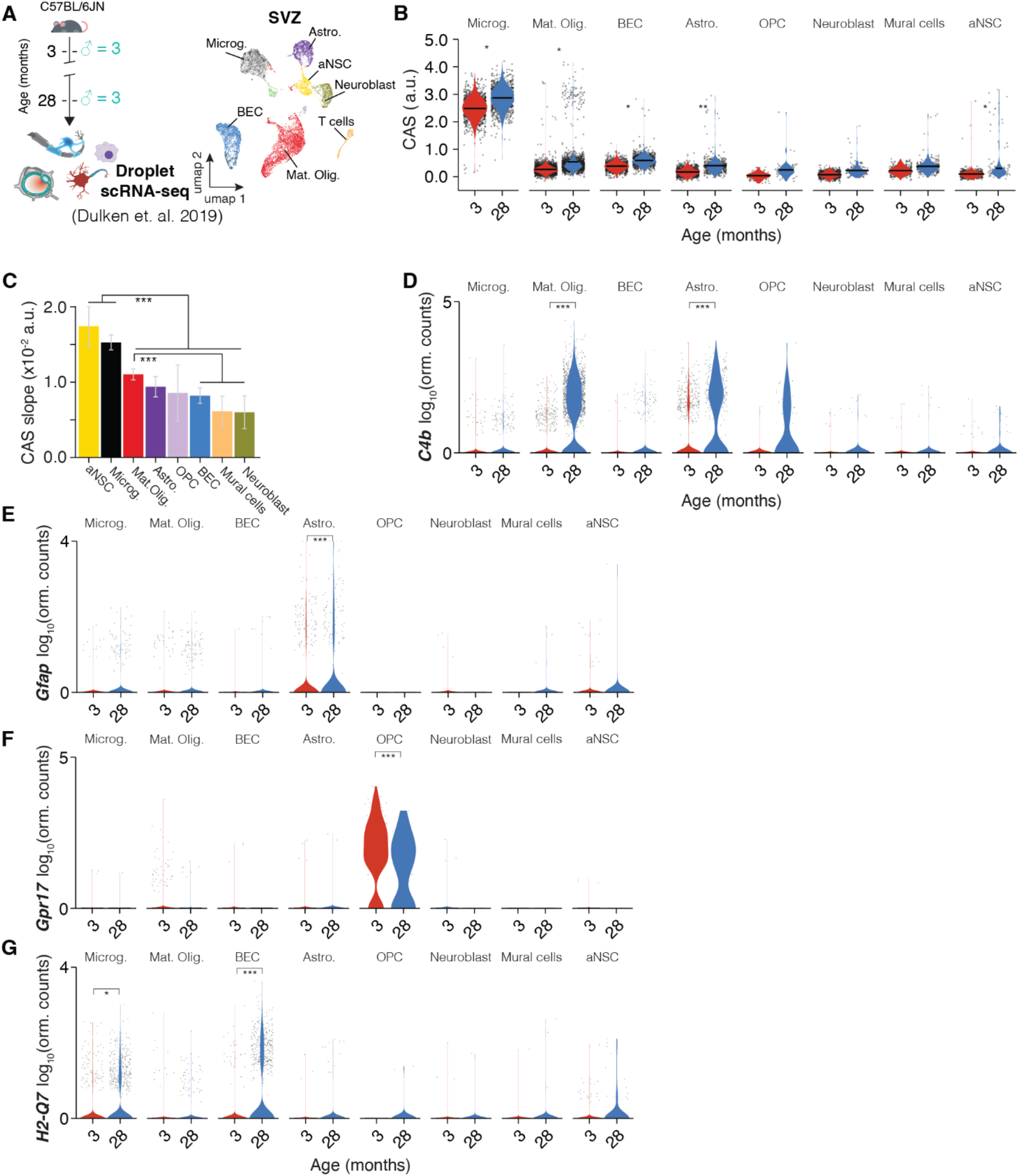
CAS analysis of SVZ scRNA-seq data (A) Meta-analysis of droplet scRNA-seq data from^27^ of cells from the SVZ. UMAP representation of all cell populations (n = 15,684 cells). (B) Violin plot representing CAS across cell types in the SVZ. Points indicate nuclei-wise expression levels, and the violin indicates average distribution of expression split by age. P values calculated with two-tailed t-test on per-replicate median of score. *** p < 0.001, ** p < 0.01, * p < 0.05. (C) CAS slope of linear regressions in (B), colored by cell type. Data are mean ± 95% confidence intervals. Two-sided Tukey’s HSD test, adjusted for multiple testing, *** p < 0.001, ** p < 0.01, * p < 0.05. The highest (least significant) Pval is indicated. (D-G) Violin plot representing (D) *C4b*, (E) *Gfap*, (F) *Gpr17* and (G) *H2-Q7* expression across cell types in the SVZ. Points indicate nuclei-wise expression levels, and the violin indicates average distribution of expression split by age. (MAST, Benjamini–Hochberg correction; false discovery rate (FDR) < 0.05 and logFC > 0.2 to be significant). *** p < 0.001, ** p < 0.01, * p < 0.05.

**Figure S12.**
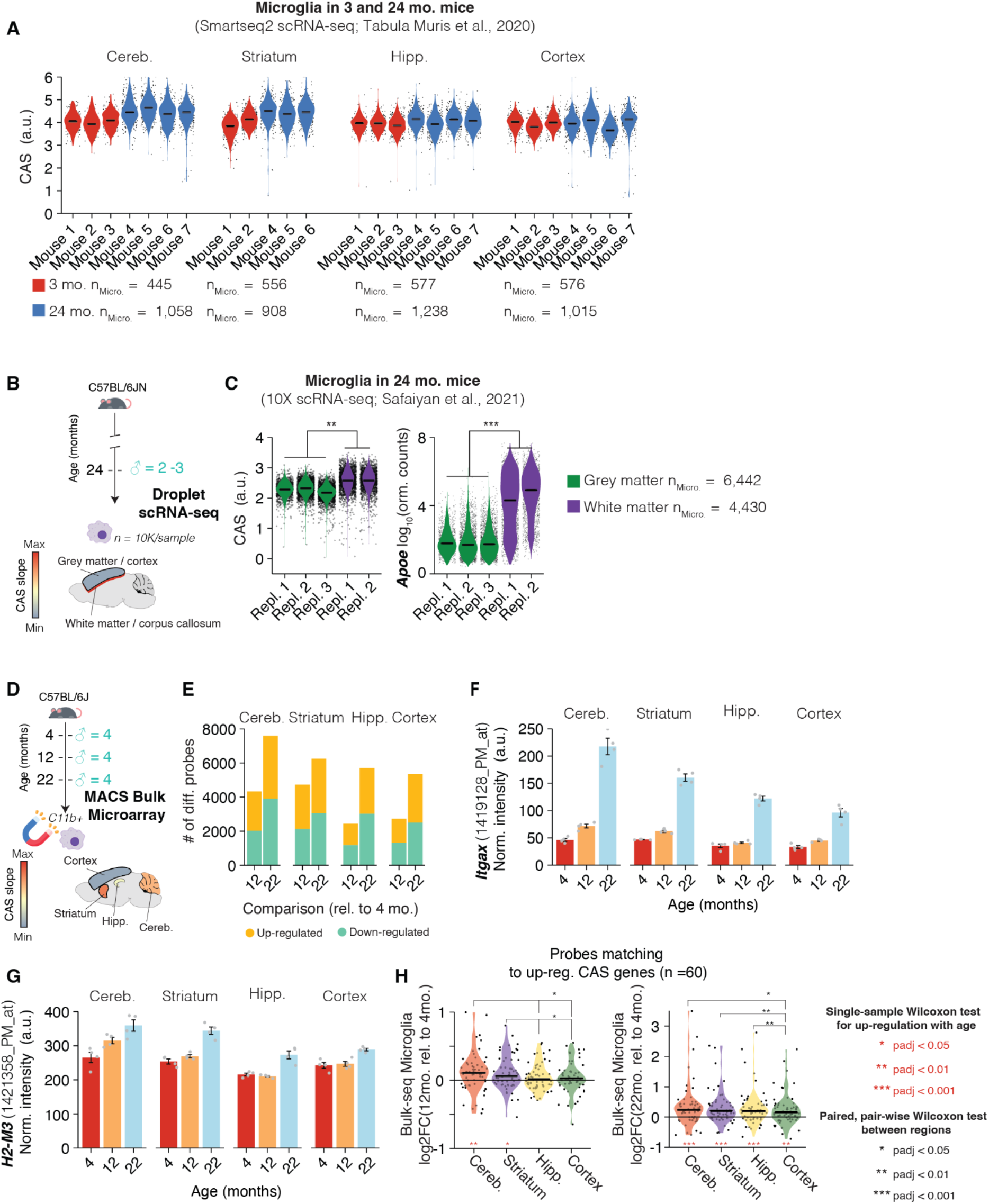
Region-specific aging of microglia is replicated across datasets (A) Meta-analysis of scRNA-seq data from ^19^ of microglia from cerebellum, striatum, hippocampus and cortex (n = 2-4 males at 3 mo. and 24 mo.). Points indicate nuclei-wise CAS, and the violin indicates average distribution of expression split by age and replicate. P values calculated with two-tailed t-test on per-replicate median of score. Benjamini–Hochberg correction; false discovery rate (FDR) < 0.05 and logFC > 0.2 to be significant). *** p < 0.001, ** p < 0.01, * p < 0.05. (B) Overview of meta analysis of scRNA-seq data from ^51^. Corpus callosum, optic tract, and medial lemniscus dissected to represent white matter, while prefrontal cortex dissected to represent gray matter (n = 2-3 males at 24 mo.). Regions colored according to CAS slopes in Figure 2G. (C) Violin plot representing CAS (left) and *Apoe* expression (right) across scRNA-seq microglia data from white matter (n=6,442 microglia) and gray matter (n=4,430 microglia). Points indicate cell-wise CAS, and the violin indicates average distribution split across tissue type and replicate. P values calculated with two-tailed t-test on per-replicate median of score. Benjamini– Hochberg correction; false discovery rate (FDR) < 0.05 and logFC > 0.2 to be significant). *** p < 0.001, ** p < 0.01, * p < 0.05. (D) Meta analysis of MACS bulk microarray data from ^52^ of microglia from the cerebellum, striatum, hippocampus, and cortex experiment overview. (n = 4 males at 4, 12, and 22 mo.). Regions colored according to CAS slopes in Figure 2G. (E) Number of probes that significantly change in expression with age in 12 mo. and 22 mo. relative to 4 mo. split by region and age, colored by up- and down-regulation. (F) Microarray expression levels of CAS gene *Itgax* in purified microglia split across regions and age. n=4 independent samples, each from tissue pooled from eight mice. Data shown mean ± SD (G) Microarray expression levels of *H2-M3* in purified microglia split across region and age. 4 independent samples, each from tissue pooled from eight mice. Data shown mean ± SD (H) Violin plot representing log2FC of up-regulated CAS genes included among the tested microarray probes split by region in 12 mo. relative to 4 mo. (left) and 22 mo. relative to 4 mo. samples. Points indicate probe-wise log2FC levels, and the violin indicates average distribution of log2FC split by region. Between-region significance determined by single-sample Wilcoxon test for up-regulation with age. In red *** padj < 0.001, ** padj < 0.01, * padj < 0.05. Between-age significance determined by paired, pair-wise Wilcoxon test between regions. In black *** padj < 0.001, ** padj < 0.01, * padj < 0.05.

**Figure S13.**
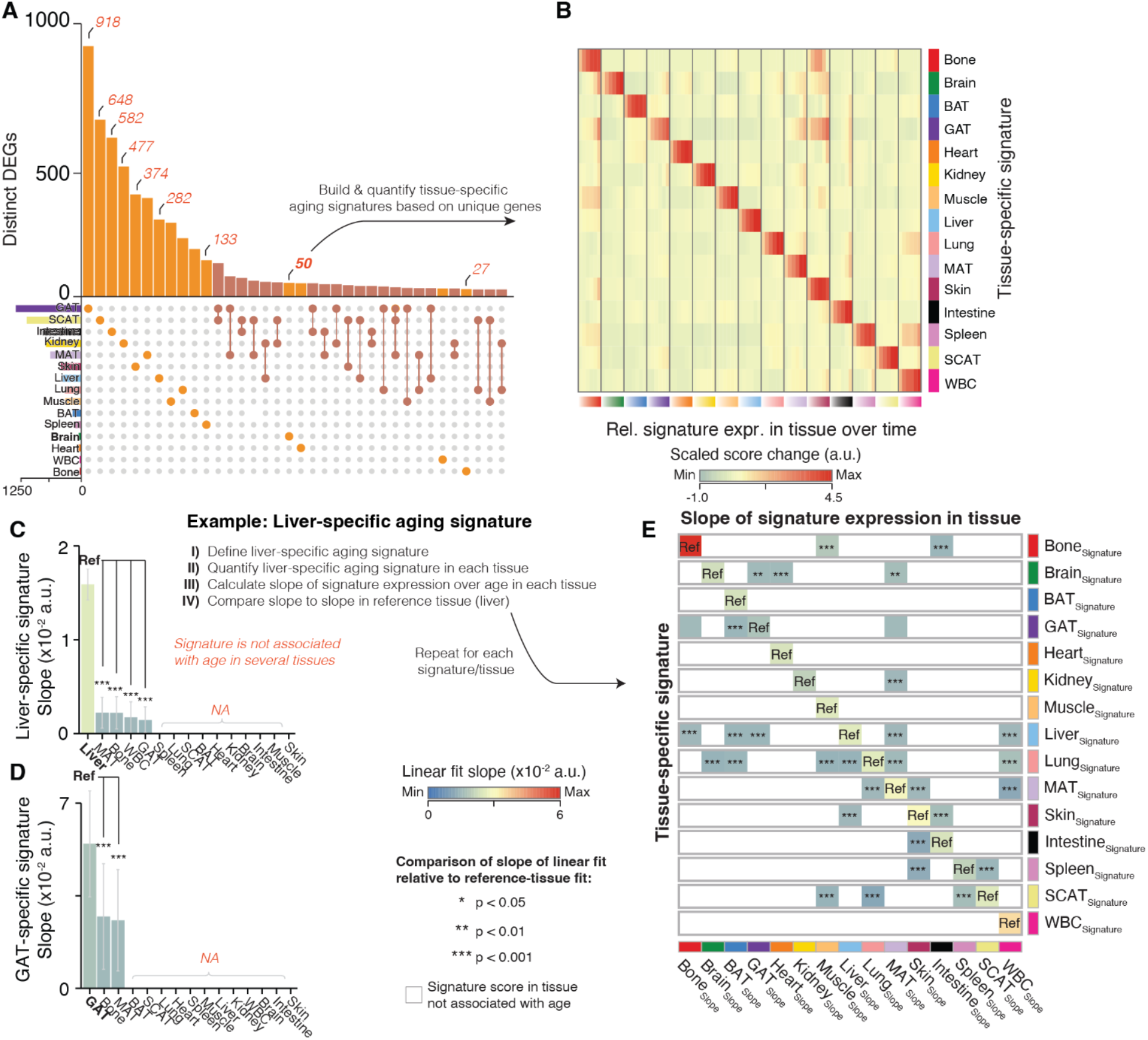
Identifying unique transcriptional signatures of aging across peripheral organs (A) Meta-analysis from *(8*) for cross-organ specificity of DEGs. UpSet plot showing a matrix layout of DEGs shared across and specific to a tissue. Each matrix column represents either DEGs specific to a tissue (single circle with no vertical lines) or DEGs shared between tissue, with the vertical line indicating the tissues that share that given DEG. Top, bar graph displays the number of DEGs in each combination of tissue. Left bottom, bar graph displays the total number of DEGs for a given tissue. Only gene sets ≥ 25 genes are shown. Unique gene sets were used to construct tissue-specific aging signatures. (B) Tissue-wise score changes with age relative to 3 months (column-wise from left to right) for tissue-specific signatures. Score changes are z-scaled within a row. (C) Slope of linear regressions for the liver-specific signature, colored by slope. The liver signature was quantified in each organ and tested for correlation with age. Tissues indicated with NA exhibited no significant association with age. Subsequently, the slope of the linear fit for each organ that exhibited association with age was compared against the reference tissue, where the signature was first identified in (i.e., the liver). Pairwise comparisons across all tissues were run and Pval corrected for multiple testing using Tukey’s method. Comparisons relative to the reference tissue were inspected to determine if the tissue-specific signature exhibits an age-related increase distinct to the reference tissue. Data are mean ± 95% confidence intervals. Two-sided Tukey’s HSD test, adjusted for multiple testing, *** p < 0.001, ** p < 0.01, * p < 0.05. The highest (least significant) Pval is indicated. (D) Analysis as in (C) for the gonadal adipose tissue (GAT) – specific signature. (E) Summarized results for each tissue-specific signature analysis. Reference tissues are indicated with ‘Ref” and slopes for each tissue with significant age-association in a given signature is indicated by color. Tissues that exhibited no association with age (NA) for a given signature were left blank. Adjusted *P* values for slope comparison of a given tissue with the reference tissue are indicated.

**Figure S14.**
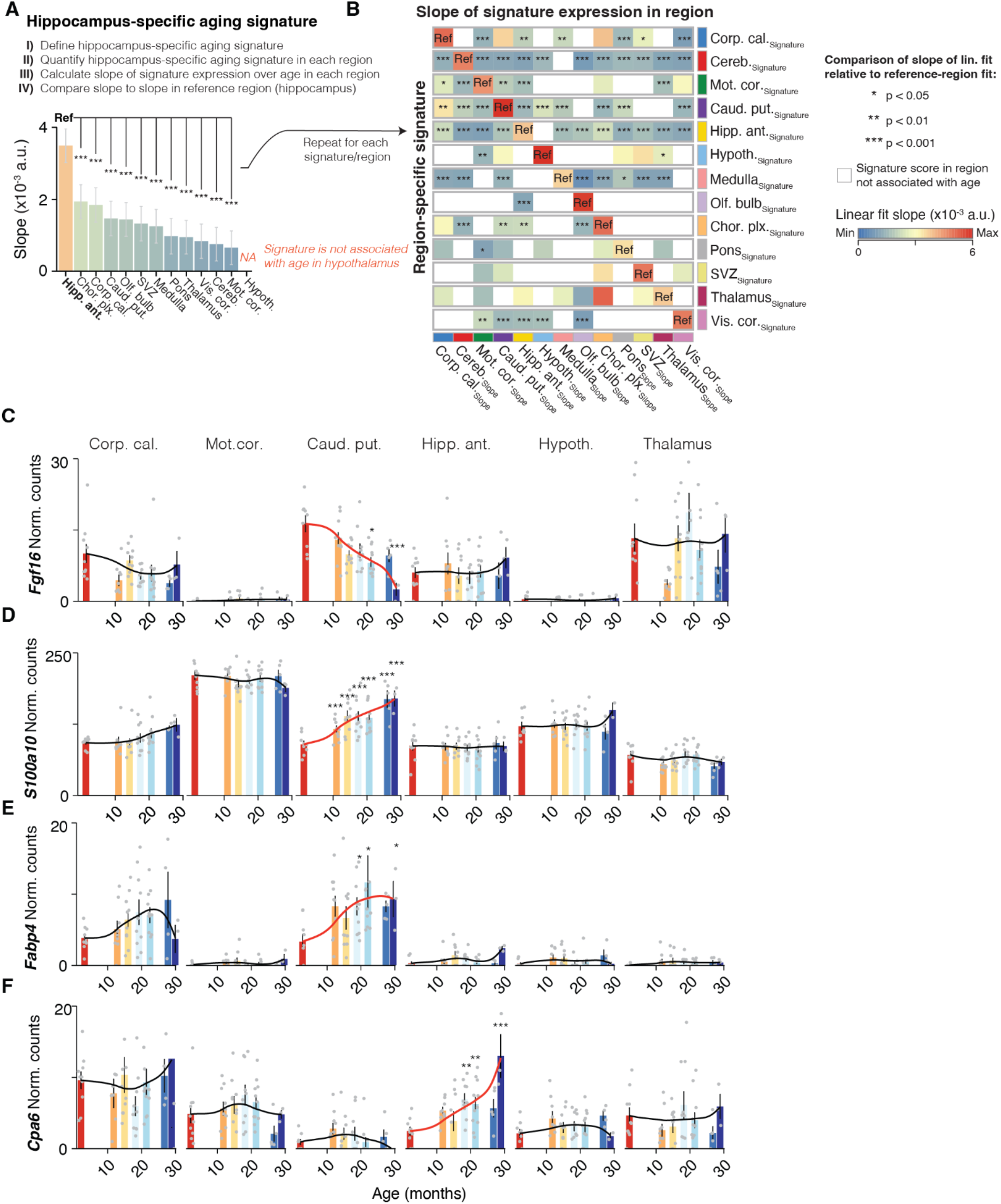
Identifying unique transcriptional signatures of aging across brain regions (A) Slope of linear regressions for the hippocampus-specific signature, colored by slope. The hippocampus signature was quantified in each region and tested for correlation with age. Regions indicated with NA exhibited no significant association with age. Subsequently, the slope of the linear fit for each region that exhibited association with age was compared against the reference region, where the signature was first identified in (i.e., the hippocampus). Pairwise comparisons across all regions were run and *P* values corrected for multiple testing using Tukey’s method. Comparisons relative to the reference region were inspected to determine if the region-specific signature exhibits an age-related increase distinct to the reference region. Data are mean ± 95% confidence intervals. Two-sided Tukey’s HSD test, adjusted for multiple testing, *** p < 0.001, ** p < 0.01, * p < 0.05. The highest (least significant) Pval is indicated. (B) Summarized results for each region-specific signature analysis. Reference regions are indicated with ‘Ref” and slopes for each region with significant age-association in a given signature is indicated by color. Regions that exhibited no association with age (NA) for a given signature were left blank. Adjusted *P* values for slope comparison of a given region with the reference region are indicated. *** p < 0.001, ** p < 0.01, * p < 0.05. (C-G) Bulk expression across proximal brain regions of genes (C) *Fgf17*, (D) *S100a10*, (E) *Fabp4*, (F) *Cpa6* that exhibit distinct changes in caudate putamen or hippocampus. Black lines indicate averaged-smoothed gene expression. The trajectory with significant age effect is highlighted. Data are mean ± s.e.m.

**Figure S15.**
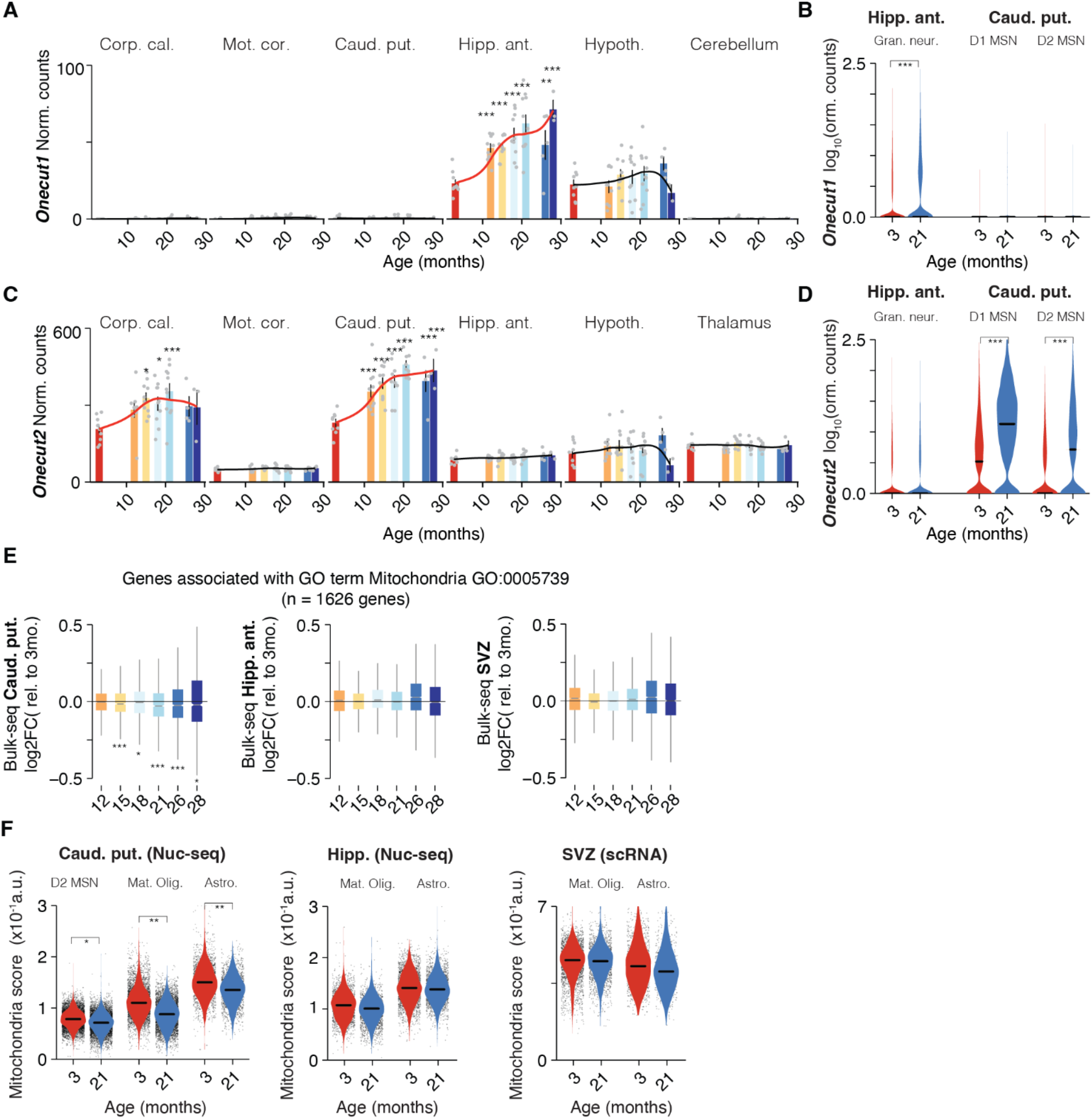
Region-specific expression of Onecut transcription factors during aging (A) Bulk expression across proximal brain regions of *Onecut1*. Black lines indicate averaged-smoothed gene expression. The trajectory with significant age effect is highlighted. Data are mean ± s.e.m. (B) Violin plot of *Onecut1* expression across neuronal cell types in hippocampus and caudate putamen. Points indicate nuclei-wise expression levels, and the violin indicates average distribution of expression split by age. (MAST, Benjamini–Hochberg correction; false discovery rate (FDR) < 0.05 and logFC > 0.2 (log2FC > 0.7) to be significant). *** p < 0.001, ** p < 0.01, * p < 0.05. (C) Same as (A) for *Onecut2*. Data are mean ± s.e.m. (D) Same as (B) for *Onecut2*. (E) Distribution of gene-wise expression changes in the caudate putamen (left), anterior hippocampus (center) and SVZ (right) with age relative to 3 months for genes associated with the GO term ‘Mitochondria’ (n = 1,626 genes). Two-sided Wilcoxon rank-sum test, adjusted for multiple testing. *** p < 0.001, ** p < 0.01, * p < 0.05. (F) Violin plot representing mitochondria RNA score across D2 MSN, mature oligodendrocytes and astrocytes in caudate putamen (left). Same for astrocytes and oligodendrocytes in the anterior hippocampus (center) and SVZ (right). Points indicate nuclei-/cell-wise expression levels, and the violin indicates average distribution of expression split by age. P values calculated with two-tailed t-test on per-replicate median of score. *** p < 0.001, ** p < 0.01, * p < 0.05.

**Figure S16.**
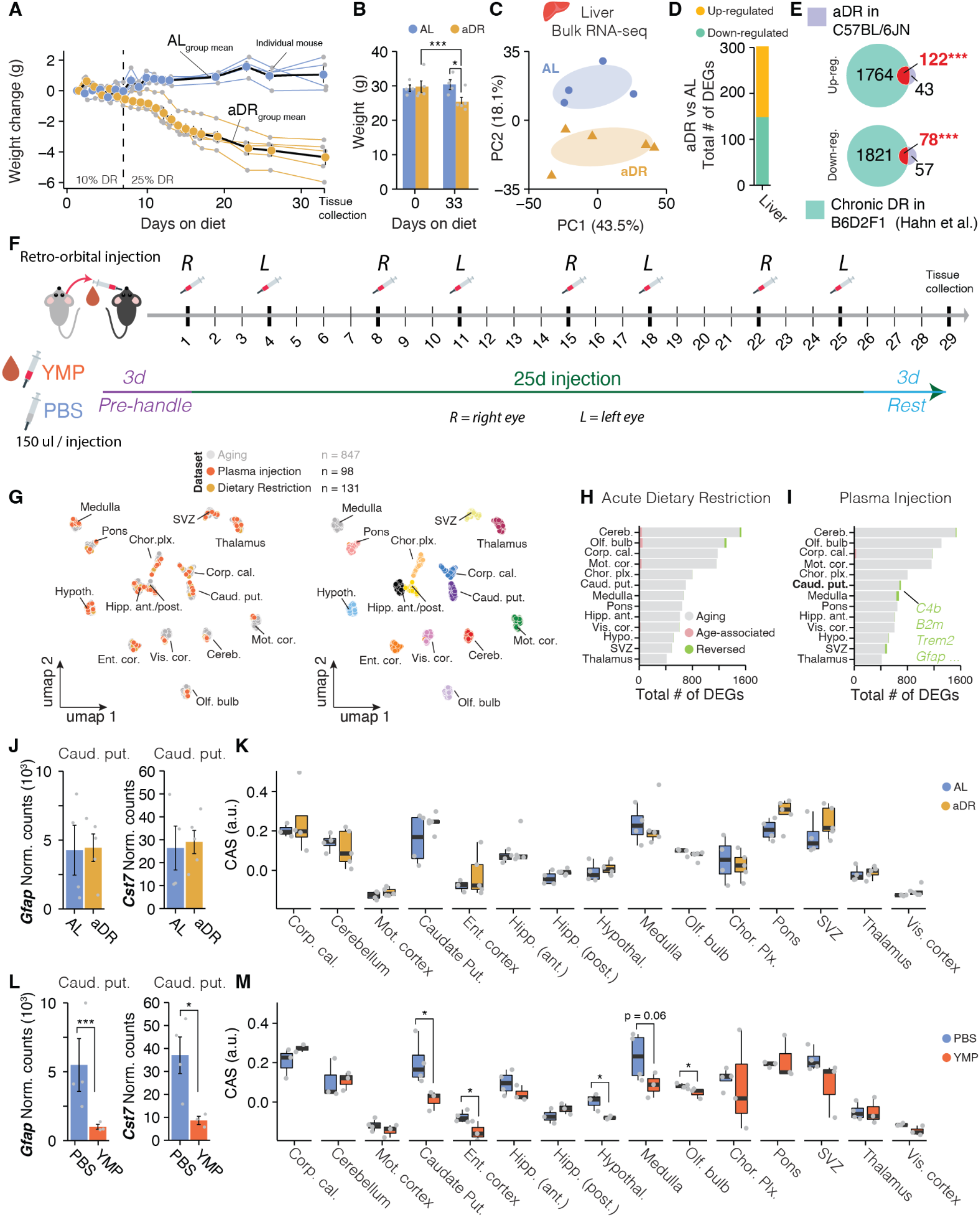
Mapping the shared and distinct expression patterns of dietary restriction and young mouse plasma injection. (A) Body weight change for AL and aDR-fed animals (n = 9). Each animal’s individual trajectory is depicted with gray points. Per-group averages are indicated by trajectories colored in black. Start of the complete 25% aDR phase is indicated. (B) Body weights for each animal at the beginning and end of the aDR phase. P values calculated with two-tailed, paired t-test. Bonferroni correction; padj < 0.05. *** p < 0.001, ** p < 0.01, * p < 0.05. (C) PCA of Bulk-seq data of liver tissue from the aDR and AL-fed mice. (D) Bar graph indicating the number of detected DEGs in liver tissue of aDR-fed mice. (E) Venn diagrams depicting the overlap of DEGs in liver of C57BL/6JN and chronically DR-fed B6D2F1 mice (data from ^57^). (F) Injection paradigm for YMP-treated animals. Injections were conducted retro-orbitally every 3-4 days. Injections were alternated between left and right eye. (G) UMAP representation of co-integrated brain region transcriptomes from rejuvenation cohorts and aging time course (n = 131 total samples from DR/AL fed mice; n = 98 samples from YMP/PBS mice; n = 847 samples from aging cohort; from Figure 1), based on the first 40 principal components. Cells are colored by experiment (left) or region (right). (H,I), Barplots showing the proportion of genes that are differentially expressed age which are reversed by (H) aDR or (I) YMP-injection. Region, and whether the genes increase or decrease with age. (J) *Gfap* and *Cst7* expression in caudate putamen. Differential expression relative to control group is indicated. Data are mean ± s.e.m. Two-sided Wald test, adjusted for multiple testing. *** p < 0.001, ** p < 0.01, * p < 0.05. (K) Boxplot representation of CAS across all regions. P values calculated with two-tailed t-test on per-replicate median of score. *** p < 0.001, ** p < 0.01, * p < 0.05. (L) same as (J) for YMP-treated mice. (M) same as (K) for YMP experiments.

**Figure S17.**
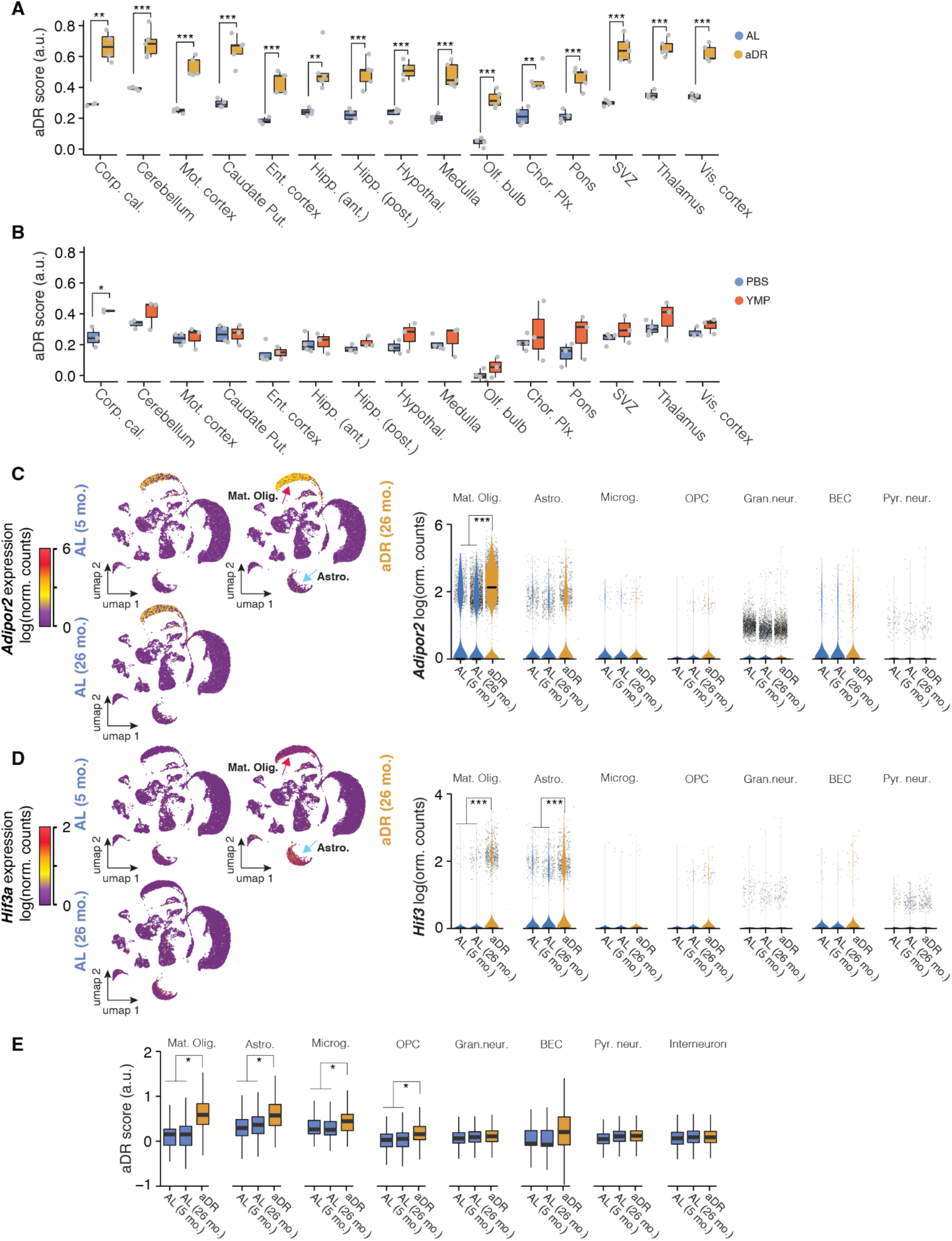
Acute DR causes brain-wide expression shifts in glia (A) CAS shifts in response to aDR across all regions. P values calculated with two-tailed t-test. Bonferroni correction; padj < 0.05. *** p < 0.001, ** p < 0.01, * p < 0.05. (B) same as (A) for YMP experiments. (C,D) Expression of (C) *Adipor2* and (D) *Hif3a* mapped onto aDR hippocampus Nuc-seq data. Quantification and statistical analysis is plotted on the right as violin plots. Points indicate nuclei-wise expression levels, and the violin indicates average distribution of expression split by age. (MAST, Benjamini–Hochberg correction; false discovery rate (FDR) < 0.05 and logFC > 0.2 to be significant). *** p < 0.001, ** p < 0.01, * p < 0.05. (E) Boxplot representation of common aDR score across all cell types. P values calculated with two-tailed t-test on per-replicate median of score. *** p < 0.001, ** p < 0.01, * p < 0.05.

**Figure S18.**
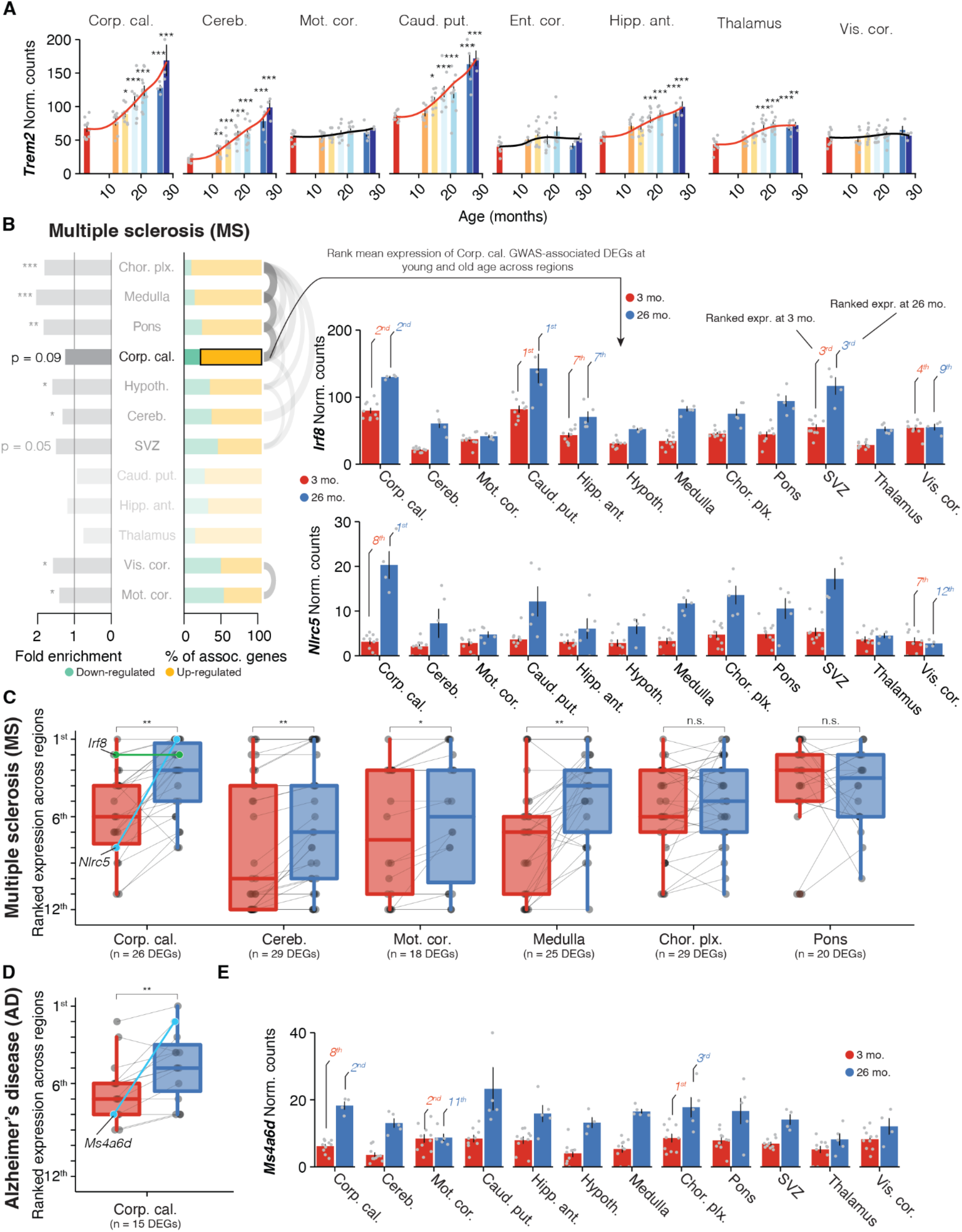
Expression of disease variant homologues across the brain is shifted as a result of age-related expression changes (A) *Trem2* expression across all bulk regions, colored by age. Red lines indicate averaged-smoothed gene expression. Data are mean ± s.e.m. (B) Analysis outline to probe brain-wide expression of GWAS genes at young and old age. Enrichment analysis of region-resolved DEGs for human GWAS variants for multiple sclerosis from Fig. 8 is depicted. The expression of corpus callosum DEGs associated with MS GWAS hits were ranked across brain regions at young (3 months) and old (26 months) of age. Expression of two exemplary genes, *Irf8* (top) and *Nlrc5* (bottom) across bulk regions at 3 mo. and 26 mo is depicted. The rank of mean expression for selected regions is highlighted. (C) Box plots displaying ranked expression changes from 3 mo. to 26 mo. of up-regulated GWAS-DEGS for multiple sclerosis across regions, filtered for regions with at least 15 genes to focus on trends with specific statistical power. Irf8 (green) and Nlrc5 (teal) rank changes in the corpus callosum are highlighted. Significance determined by paired, pairwise Wilcoxon test between regions. *** padj < 0.001, ** padj < 0.01, * padj < 0.05. (D) Box plot displaying ranked expression changes from 3 mo. to 26 mo. of up-regulated GWAS-DEGS for Alzheimer’s in the corpus callosum. *Ms4a6d* (teal) highlighted. Significance determined by paired, pairwise Wilcoxon test between regions. *** padj < 0.001, ** padj < 0.01, * padj < 0.05. (E) *Ms4a6d* expression across bulk regions at 3 mo. and 26 mo. The rank of expression for selected regions is highlighted.

## Supplementary table legends

Table S1. Bulk-seq region marker genes (separate file)

Marker gene analysis for bulk-seq regions from Figure 1C and Figure S3A.

Table S2. DEG detection across regions (separate file)

DEGs and the number of tissues in which they were detected. Further indicates CAS and region-specific DEGs from Figure 2A,B and Figure 6A.

Table S3. Age-correlated genes (separate file)

List of genes with significant correlation with age from Figure 1H.

Table S4. WGCNA modules (separate file)

List of modules as detected by WGCNA from Figure 1 I,J. Enriched Gene Ontology terms and cell type markers are indicated. Modules with significant age-related change are noted.

Table S5. Functional enrichment of CAS genes (separate file)

Complete list of enriched Gene Ontology terms for CAS genes from Figure 2C.

Table S6. 10X Visium cluster marker genes (separate file)

Marker gene analysis for 10X Visium clusters from Figure S7.

Table S7. DEGs detected in bulk- and 10X Visium (separate file)

Comparison of age-related DEGs found in bulk-seq and 10X Visium data of corpus callosum/white matter cluster and motor cortex/cortex cluster, from Figure 3C.

Table S8. Functional enrichment of caudate putamen-specific DEGs (separate file)

Complete list of enriched Gene Ontology terms for caudate putamen-specific DEGs from Figure 6E.

Table S9. Cell dispersion analysis for hippocampus and caudate putamen (separate file)

Quantification of cell-specificity of DEGs found in hippocampus or caudate putamen from Figure 6G

Table S10. aDR- and YMP-induced DEGs detected across regions (separate file)

aDR- or YMP-induced DEGs and the number of tissues in which they were detected. Further indicates aDR and region-specific DEGs from Figure 7A,B,J.

Table S11. **Functional enrichment of aDR-induced DEGs in the cerebellum (separate file)** Complete list of enriched Gene Ontology terms for aDR-induced DEGs in the cerebellum from Figure 7I.

Table S12. Functional enrichment of YMP-induced DEGs in the SVZ (separate file)

Complete list of enriched Gene Ontology terms for YMP-induced DEGs in the SVZ from Figure 7I.

Table S13. Signature genes of rejuvenated SVZ and aged aNSCs (separate file)

List of genes utilized to build the YMP signature for the SVZ (bulk) and signature of aged aNSCs based on DEGs found in ^27^

Table S14. DEGs with human GWAS homologue (separate file)

List of DEGs with human disease GWAS homologue for AD, PD and MS from Figure 6. Colored cells indicate regions where we discovered a significant enrichment of disease-related genes.

